# Target Specific Conformational Flexibility and Electrostatic Complementarity Modulate LTNF10 Peptide Binding to Phospholipase A2 Variants from Snake Venoms

**DOI:** 10.64898/2026.10.30.761852

**Authors:** Aayushi Gupta, Mohan Maruthi Sena, Bipin Singh

**Affiliations:** Centre for Life Sciences, Mahindra University, Hyderabad, India; Department of Molecular Oncology, Cancer Institute (WIA), Chennai, India

**Keywords:** Bioactive Peptides, Antivenom, Molecular dynamics simulation, Free energy calculations, Snakebite envenoming, Snake Venom Toxins

## Abstract

Phospholipase A2 is one of the primary cytotoxins present in snake venoms, causing tissue damage through the hydrolysis of phospholipids in the cellular membrane. We prioritised and investigated in detail the binding and conformational dynamics of a ten-residue short peptide (LTNF10) from Lethal Toxin Neutralizing Factor from Opossum against snake venom Phospholipase A2 (svPLA_2_) variants. LTNF10 was prioritised from the curated list of peptide inhibitors. A charge-modulating mutation from aspartic acid to lysine at the fifth position in LTNF10 was observed to enhance the binding affinity to svPLA_2_ variants from *Elapids*. The mutant LTNF10 shows selectively enhanced binding affinity toward Group I svPLA_2_ from Elapidae. Free energy landscapes of LTNF10 and mutant LTNF10, and binding pocket-volume analysis of svPLA_2_ revealed that Lys5 substitution increases conformational flexibility of mutant LTNF10, allowing it to form favourable binding conformations, while svPLA_2_ shows a reduction in pocket volume. Conversely, the mutant LTNF10 shows reduced binding affinity in Group II svPLA_2_ from *Viperidae* while demonstrating reduced flexibility and a less significant decrease in binding pocket volume in svPLA_2_. These findings demonstrate that active-site charge microenvironments, binding pocket and structural differences in catalytically relevant loop regions influence LTNF10 binding to svPLA_2_. The differential binding affinity was primarily driven by improved non-covalent interactions, specifically reduced electrostatic repulsions and enhanced electrostatic interactions in Group I svPLA from *Elapids.* This was influenced by the enhanced flexibility of mutant LTNF10, suggesting the role of a complex scenario of enthalpy-entropy compensation. The understanding of dynamic and structural mechanisms of LTNF10 and svPLA_2_ interactions and binding offers key insights for designing target-specific peptide therapeutics against diverse svPLA2 toxins.

## Introduction

Snakebite envenoming remains a high-impact and under-recognized public-health problem across large parts of the tropical and subtropical world. In June 2017, the World Health Organisation (WHO) formally designated snakebite envenoming as a highest priority neglected tropical disease (NTD) (1,2). It has been neglected largely because snake envenomation is a rural problem and multinational pharmaceutical industries prioritize profit-oriented endeavours. (3). The lives affected each year are estimated to be 1.8-2.7 million globally, out of which 138,000 cases are fatal.(4)58,000 cases of these are from India, and 400,000 people are left with permanent disability(5). According to recent research conducted across 11 Indian states covering over 60 million Indian population, the snakebite incident rate is 4.4 per 100,000 population per year(6). This disproportionate burden reveals the profound social and economic dimensions of the disease, weighing heavily on Indians.(7).

### Snake Venom Variation

Developing antivenom is complicated by the fact that venom composition varies within species and across regions, broadening the toxin target profiles that antibody therapeutics need to neutralise. Research on the spectacled cobra (*Naja naja*) has shown that biogeographical variation in venom profiles undermines the feasibility of a single pan India therapy (8). Subspecies level differences in the North African mountain viper (*Vipera monticola*) further illustrate how ecological and taxonomic context shape toxin repertoires (9). Such variation is driven by evolutionary and environmental pressures, with direct consequences for clinical treatment efficacy (10). Since antivenoms act by introducing antibodies that bind and neutralize venom components, shifts in toxin abundance, variation in sequence, and structure can reduce their binding efficiency, underscoring the need for regionally informed and molecularly precise strategies in snakebite therapy.

### Snake venom Phospholipase A as a target

Snake venom Phospholipase A (svPLA_2_) is a phospholipase that hydrolyzes the sn-2 ester bond of phospholipids, releasing lysophospholipids and free fatty acids (often arachidonic acid). The svPLA targets phospholipid membranes rather than a single receptor. This reaction destabilizes membranes and triggers downstream toxic effects such as neurotoxicity, myotoxicity, and anticoagulation. svPLA s act through conserved catalytic residues and calcium binding, but their toxic manifestations diverge between elapids (neurotoxic, Group I) and vipers (myotoxic, Group II). This dual functionality underscores their evolutionary adaptation to prey immobilization and defense. Elapid PLA (Group I) are highly enzymatic and often neurotoxic, while viperid PLA (Group II) show greater structural diversity, combining enzymatic activity with non enzymatic myotoxic effects. The two variants of svPLA_2_ include Asp49 variants from Group I, which are catalytically active and responsible for the enzyme’s primary function of phospholipid hydrolysis. In contrast, Lys49/Ser49 variants from Group II are catalytically inactive but still retain significant myotoxic (muscle-damaging) and cytotoxic effects(11,12).

The catalytic region of His48–Asp49 is central to the enzymatic mechanism, where His48 acts as a general base to activate water for nucleophilic attack, while Asp49 stabilizes the transition state and coordinates the essential calcium ion. The calcium binding loop (Gly30–Gly33, Tyr28) ensures proper substrate orientation, and hydrophobic channel residues (Trp31, Tyr52, Lys56) guide phospholipids into the active site. Calcium plays a dual role of stabilizing the negative charge on the phosphate group and enhancing nucleophilicity, making it indispensable for enzymatic activity(13,14). In elapids (Group I PLA) like cobras and kraits, these enzymes retain the Asp49 catalytic residue and a conserved calcium binding loop. They are highly enzymatically active and frequently associated with neurotoxic effects, disrupting presynaptic membranes and blocking neurotransmission. Their structural motifs favor precise substrate binding and efficient catalysis. While in vipers (Group II PLA) like the Sindh saw-scaled viper and Russell’s viper, this shows diversity. Some retain Asp49 and remain catalytically active, while others carry substitutions such as Lys49 and Ser49, which abolish enzymatic activity but confer direct myotoxic and cytolytic properties. Lys49 and Ser49 PLA s disrupt membranes through non enzymatic mechanisms, contributing to local tissue damage, edema, and necrosis. Thus, viperid PLA s combine enzymatic and non enzymatic toxicity, explaining their severe local effects (15,16).

svPLA enzymes are consistently among the most abundant components of snake venoms, accounting for 30–50% of total protein content in viperidae venoms and 10–30% in elapids, according to proteomic-based studies (13,18,19). They catalytically hydrolyze phospholipids in cell membranes, leading to local tissue necrosis, edema, anticoagulation, neurotoxicity, and systemic organ damage. Structurally, svPLA s are relatively conserved across species, with a well defined calcium binding loop and catalytic site that provide rational entry points for inhibitor design (13). Functionally, they act not only as enzymes but also as pharmacological effectors, amplifying venom pathology through inflammatory cascades.

Computationally designed peptides targeting conserved PLA motifs could provide scalable and precise interventions against one of the most abundant and toxic venom components (12).

### Current treatment against snakebite envenomation

Recent innovations in antivenom development are moving toward highly targeted molecular strategies that address the limitations of conventional polyclonal antibody-based therapies. Traditional antivenoms often suffer from restricted geographical efficacy due to venom variability, as well as batch to batch inconsistency and risks of adverse reactions. To overcome these challenges, a research work led by Laustsen and Karthik showcases how monoclonal antibodies can be designed to be capable of broadly neutralizing long chain three finger α neurotoxins, representing a step toward standardized, reproducible therapies with defined molecular specificity (21–24). In parallel, Baker’s Lab has developed an AI platform to design de novo small proteins that bind both short and long chain α neurotoxins with high affinity (24). These engineered proteins exemplify a modular approach that could be adapted across diverse venom repertoires, potentially enabling pan species coverage. On the other hand, Varespladib (LY315920), a small molecule which was originally investigated as a drug for the treatment of acute coronary syndrome and acute chest syndrome by targeting the precursor of inflammatory sPLA2 (secretory PLA2), was later repurposed for neutralisation of PLA2 enzymes in snakes. (24)

Together, these strategies highlight a shift from empirical serum production to rational, precision-engineered therapeutics, underscoring how antibody engineering, AI driven protein design and peptide therapeutics have the potential to reshape the future of snakebite treatment.(24)

### Peptide-based therapeutics

Peptide based strategies against snake venom have steadily evolved from receptor mimicking decoys to modern computationally designed scaffolds. Early work demonstrated that short synthetic peptides could block α neurotoxins by imitating the nicotinic acetylcholine receptor binding site, thereby preventing paralysis (25,26). Plant-derived phyto peptides have been investigated for their ability to neutralize venom toxins, offering a natural and potentially accessible therapeutic avenue (27). Peptide based approaches have progressed from conceptual receptor mimics to rationally engineered therapeutics, underscoring their potential as precise and scalable complements to conventional antivenoms (25,26).

LTNF10 peptide is derived from the lethal toxin neutralizing factor (LTNF) protein found in the blood serum of the opossum, a species uniquely known to survive fatal snake bites (28–31). This evolutionary adaptation provides strong evidence of functional relevance against venom toxins, positioning LTNF10 as a biologically validated scaffold for antivenom design (29,32).

Thus, peptide-based therapeutics represent another promising avenue in the development of next-generation antivenoms. Peptide-based approaches offer a more targeted solution. Ultimately, mitigating the global snakebite crisis requires a comprehensive strategy that extends beyond a single therapeutic solution. Introducing a computational approach to modifying and designing peptides can be an avenue to tackle current issues and provide a target-based approach. In this study, we evaluated a set of peptide candidates from diverse sources and optimized and validated the most promising candidate (LTNF10) against svPLA_2_s from Elapidae and Viperidae through rational mutagenesis, molecular dynamics (MD) simulations, and free energy calculations based on molecular interactions between LTNF10 and svPLA_2_s.

## Materials and Methodology

### Target Toxins and Peptide Candidates

Target protein data collection: All svPLA_2_ sequences and structures were collected using UniProt (33), RCSB (34), and Venomzone®2015 server (35)developed by the Swiss-Prot group at the SIB Swiss Institute of Bioinformatics (33,33–37). A total of 35 protein structures of svPLA_2_ were collected from RCSB across two venomous snake families, named Elapidae and Viperidae (Table S1). The four target svPLA2s were taken from the big four snake species, from Elapidae *Naja naja, Bungarus caeruleus,* and from Viperidae, *Echis carinatus* and *Daboia russelii*.

Peptide data collection - We obtained potential peptides by literature survey and open-source peptide sequence libraries such as PEPotec Immuno Peptide libraries by Thermo Fisher Scientific, PepTrack™ Peptide Libraries by JPT Peptide Technologies, Antimicrobial Peptide Database (APD), Phage display libraries, and in silico peptide libraries (Table S2) (38,38–40).

Peptide candidates were designed from proteins with potential anti-venom activity using the GenScript peptide library design platform (41) which enables systematic generation of short sequences. Peptide sequences were generated using an overlapping fragments strategy. Guided by this approach, we derived peptides from two source proteins previously implicated in toxin neutralization: Turmerin and Snakin Z (42). The library was screened computationally, and peptides were selected based on their predicted toxicity and hemolytic activity profiles, ensuring prioritization of candidates with favorable safety characteristics. From these templates, four 15 residue peptides (TP1, TP2, SP1, and SP2) were taken. These peptides were subsequently advanced for docking and binding affinity evaluation against representative svPLA enzymes.

From all the peptide candidates considered, we prioritized 7 peptides for further evaluation using multiple criteria, such as solubility using PeptiVerse (43), hemolytic activity using HemoPI2 (44), and toxicity using Tox-Pred3 (45). PeptiVerse is a unified AI powered platform that predicts therapeutic peptide properties using both amino acid sequences and chemically modified SMILES representations. It enables rapid evaluation of biophysical and developability features, helping researchers design and optimize peptide candidates for drug discovery. HemoPI2 is a robust machine learning–based platform developed to evaluate the hemolytic activity of peptides. This tool offers both classification and regression modules tailored to diverse experimental needs. Tox-pred3 predicts the toxicity of peptides based on a machine learning-based infrastructure that is trained on curated datasets of toxic and non-toxic peptides.

### Molecular docking

Peptide candidates with no three-dimensional structure available were modeled using trRosetta (46). trRosetta uses a deep-learning-based neural network trained on large protein datasets to predict peptide structure by calculating inter-residue distance distributions and orientations from sequence co-evolutionary couplings. These co-evolutionary couplings are then converted into geometric constraints that are fed into the Rosetta energy minimization pipeline. Selected peptide candidates from those that were prioritized were docked against svPLA_2_ corresponding to the big four snake species using Auto-Dock Crank Pep (ADCP) (47), a specialized docking program for peptides that uses a Monte-Carlo based folding algorithm to explore peptides conformational space during molecular docking.

The protonation states of amino acid residues were determined by in-built function in ADCP. ADCP achieves this by assigning standard protonation states at physiological pH (∼7.0). According to standard protonation, acidic residues are treated as deprotonated, basic residues as protonated, and neutral residues remain unchanged. Additionally, ADCP accounts for histidine tautomerism, allowing either HID (proton on ND1) or HIE (proton on NE2) to be assigned depending on the local environment. Three of the four systems investigated in present study (PDB IDs 4QEM, 2WQ5, 1U4J) are catalytically active, Ca² dependent svPLA2. However, the Ca^2+^ was absent in 4QEM, 1U4J and 2QHD. The 2QHD is a Ca^2+^ independent Ser49-svPLA2 (48), and Ca^2+^ was absent in its crystal structure. Structural work on this protein shows that its Ca² binding loop does not adopt a conformation favorable for Ca² coordination. Considering these facts, for a fair and consistent comparison and current modelling limitations, we have not included Ca^2+^ during system preparation for 2WQ5 (49). Crystal water was removed prior to docking as a standard preprocessing step for ADCP docking. There were no missing residues present in svPLA_2_, as per sequence information provided by corresponding UniProt records. (4QEM:P59071; 2QHD: P48650; 2WQ5:P15445; 1U4J: Q6SLM1).

The protein is treated as a rigid molecule, while the peptide is considered flexible. ADCP folds a peptide in the potential energy landscape created by the protein to predict the high-quality protein-peptide docked complex.

ADCP uses the AutoDock scoring function, adapted for peptides. ADCP scoring function includes van der Waals interactions, hydrogen bonds, electrostatic interactions, desolvation energies (hydrophobic effect) and a torsional entropy penalty, which accounts for peptide flexibility. Each docking run generates multiple peptide conformations, and the score is computed for each pose. Poses are grouped by similarity, and the lowest energy cluster centre is usually taken as the representative binding mode.

After the molecular docking process with prioritized peptide candidates, the interactions between the peptide and target site were analyzed using LigPlot+(50). LigPlot+ takes 3D protein–ligand structures, calculates hydrogen bonds and hydrophobic contacts, and produces 2D schematic diagrams. It uses minimum distance cutoff and angle between atoms from the ligand (peptide) and target protein residues as criteria to identify non-covalent interactions and then generates 2D schematic representations of binding site interactions between peptide and protein molecule (50).

### Peptide Modification

We performed positional scanning for all the positions in LTNF10 (LKAMDPTPPL) (28,29) using Foldx (51). Foldx uses an empirical force field approach and calculates ΔΔG based on physicochemical properties. It takes into consideration the van der Waals, electrostatics, hydrogen bonds, water bridges, entropy, and solvation factors (51). We verified FoldX results with mCSM-PPI2 (52) and MutaBind2 (53), where mutations at certain positions were preferred based on the molecular interactions and positional scanning. mCSM-PPI2 uses a machine learning-based approach that uses graph-based signatures and is trained on experimental free energy difference data (ΔΔG). It takes the local structural environment around the mutation site into consideration and uses patterns found in distances between residues along with chemical properties of accompanying residues as features (52). Meanwhile, MutaBind2 uses an ensemble approach that combines multiple algorithms and integrates structural and evolutionary features. It uses machine learning algorithms with structural descriptors and is specifically designed for the prediction of protein-peptide binding affinity upon single or multiple mutations (53). We optimized LTNF10 by introducing point mutations, leading to the mutant LTNF10 candidate peptide, referred to as LTNF10-MT in this study.

### Molecular Dynamics Simulations

MD simulations were performed for svPLA_2_s-LTNF10 and svPLA_2_s-LTNF10-MT complexes. All MD simulations were conducted using GROMACS 2022.3 version (54). A cubic box was used for the simulation, specifying a solute-box distance of 1.0 nm, which means that there is at least 2.0 nm between any two periodic images of our complex. CHARMM27 force field (55) with CMAP (grid-based energy correction map) corrections was used for all MD simulations. CHARMM27 is known to provide stable MD trajectories and the best fit to experimentally determined conformational flexibility parameters (56,57). TIP3P (58) Water molecules were incorporated to ensure compatibility with the CHARMM27 force field, which was parameterized using this model. Particle Mesh Ewald method (59) was used for long-range electrostatics. Following solvation of the system with TIP3P water molecules, the net charge of each svPLA - peptide complex was neutralised by replacing an appropriate number of water molecules with Na or Cl counterions using the GROMACS genion utility.

The systems were minimized using steepest descent minimization until they converged with a force tolerance of 1000 kJ/mol/nm. The first equilibration step involved a 1 ns simulation in the canonical ensemble (NVT) (60) using the modified Berendsen thermostat (61). The second equilibration phase consisted of a 1 ns simulation conducted under an isothermal–isobaric (NPT) ensemble (62,63), during which positional restraints were maintained on the protein backbone. Harmonic positional constraints (64) were applied to the toxin protein and peptide in each complex. The LINCS algorithm (65) was employed to constrain all bonds involving hydrogen atoms, ensuring numerical stability throughout the simulation. Pressure was maintained at 1 bar using the Parrinello-Rahman barostat (63), while temperature for all equilibrations was kept constant at the initial (lowest) TTMD (Thermal Titration Molecular Dynamics) temperature value through the v-rescale method (66), which is a modified version of the Berendsen thermostat. All MD simulations were performed in triplicate for 100ns under periodic boundary conditions. Further, we performed extended MD simulations for 1U4J svPLA_2_-LTNF10-MT (Rep-3) and 4QEM svPLA_2_-LTNF10-MT (Rep-2) to 200ns to ensure equilibration was achieved within 100ns (Figure S30 and Table S23). A continuous MD trajectory of 200 ns was generated using the previous 100ns MD run and the checkpoint files generated by GROMACS, which led to exact continuation rather than two independent MD runs.

### Molecular Simulation Analysis

Analysis of MD simulation trajectories for computing RMSF, SASA, DSSP, hydrogen bonds, and PCA was done using the built-in features available with the GROMACS (54) simulation package. For PCA, MM/PBSA and binding-pocket volume analyses, only the equilibrated segment of each MD trajectory containing 1001 frames, identified from the plateau region of the Cα-RMSD profile of the svPLA2 target (Figures S20–S23), was analysed. The g_mmpbsa (67) is an external software package used for computing the binding free energy difference for protein-ligand complexes. It uses a post-processing approach to calculate binding free energy based on snapshots from the MD simulation trajectory. It consists of three main energy components, which are molecular mechanics energy (MM), polar solvation energy (PB), and non-polar solvation energy (SA). Molecular mechanics energy includes van der Waals and electrostatic interactions, polar solvation energy includes ligand interactions and behaviour in water, and non-polar solvation energy includes the hydrophobic effect.

g_mmpbsa (67) integrates the Molecular Mechanics Poisson–Boltzmann Surface Area (MM/PBSA) approach directly into the GROMACS environment, making binding free energy calculations both accessible and reproducible. Algorithmically, it operates by extracting representative snapshots from molecular dynamics trajectories and evaluating the free energy of the complex, receptor, and ligand separately. In the updated g_mmpbsa package developed by Rashmi Kumari, solvation and non-solvation energies are calculated through a refined MM-PBSA framework that integrates GROMACS trajectories with APBS-based continuum electrostatics. Specifically, the solvation energy is decomposed into polar and non-polar contributions. The polar term is obtained by solving the Poisson–Boltzmann equation to capture electrostatic interactions between the solute and solvent, while the non-polar term is estimated from solvent-accessible surface area (SASA)-dependent empirical models (68). In contrast, non-solvation energies encompass the molecular mechanics components that compose van der Waals and electrostatic interactions, which are directly derived from the force field used in GROMACS simulations. By combining these terms, g_mmpbsa provides a residue-level decomposition of binding free energy, enabling researchers to distinguish between solvent-mediated stabilization and intrinsic molecular interactions. This dual-layered calculation not only improves accuracy but also enhances interpretability, enabling analysis of energetic hotspots in protein–ligand complexes and guiding rational drug design. By averaging these values across many conformations, it provides a statistically robust estimate of binding free energy, with options for per residue decomposition to identify key stabilizing interactions at the residue level (69,70).

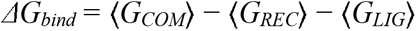

where each term on the right-hand side is given by:

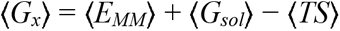

In turn, *ΔG_bind_*can also be expressed as:

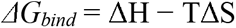

ΔH can be decomposed into its constituent terms:

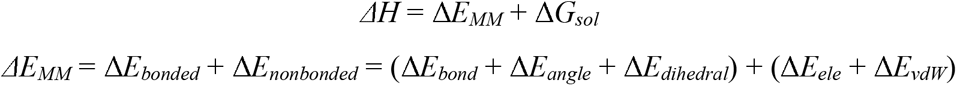

We also used two complementary methods for calculating binding affinity using PRODIGY (71), which is based on structural descriptors, and Boltz-2 (72), which is based on a deep learning biomolecular foundation model that jointly predicts 3D structures and binding affinities. On one hand, PRODIGY (71) predicts binding affinity between protein-protein complexes by computing structural features of protein-protein interfaces, which are based on intermolecular contacts, buried surface area, and residue composition. It counts the number and type of atomic contacts across the binding interface and the buried surface area (BSA) in the complex, where more contacts and larger BSA values contribute to stronger binding. Residue composition also influences binding affinity, where charged residues contribute to electrostatic interactions and apolar residues contribute to hydrophobic stabilization. PRODIGY uses statistical models built from structural descriptors (71). We used representative structures extracted from cluster analysis of MD simulation trajectories for binding affinity prediction using PRODIGY. On the other hand, Boltz-2 (73) calculates protein-peptide binding affinity by combining structural modeling and empirical scores using a foundational model trained on large biomolecular datasets. It is built on the previous models of Boltz-1 (74) and Alphafold3 (75,76) but adds controllability features for affinity prediction. It initially works with sequence information for modeling and integrates geometric features of the complex with statistical mechanics to predict binding affinity. It learns from large datasets of protein–ligand and protein–protein binding affinities.

Binding pocket volumes were calculated using CavitOmiX (Innophore GmbH, Graz, Austria), a standalone, cavity-detection plugin built on Innophore’s *Catalophore* point-cloud cavity technology (77). CavitOmiX identifies cavities using a custom grid-based implementation of the LIGSITE (78) algorithm in which the protein structure is mapped onto a three-dimensional grid, and empty (solvent) grid points are scanned along multiple fixed directions. The corresponding hydrophobicity module of the program VASCo (79) was used to analyse the hydrophobicity of the cavities.

The stable part of the molecular dynamics trajectory of svPLA2 (1001 frames, 10 ps interval) was first processed to extract individual frames as separate coordinate files using trjconv (54) from the stable part of the trajectories. A custom Python script was used to automate cavity detection.

## Results

### Selection of snake venom Phospholipase A2 candidates

Out of the 35 target structures available in RCSB PDB (34,36). We selected svPLA_2_s from the big four snake species, which consist of Elapidae species *Naja naja* and *Bungarus caeruleus* and Viperidae species *Echis carinatus* and *Daboia russelii.* These species are more significant in contributing to the fatality rate, among others, in India. The selection of svPLA_2_ structures was based on availability, recentness, and resolution. Selected target PDB IDs are shown in Table 1.

**Table 1.** Structural and taxonomic details of svPLA from big four snake species. The entries contain Group 1 PLA s from elapids (Indian krait, Indian cobra) and Group 2 svPLA s from vipers (Sindh saw scaled viper, Russell’s viper).

| **PDB ID** | **Species** | **Common name** | **Family** | **svPLA<sub>2</sub> family** | **Sequence length** | **Resolution** | **Reference** |
|---|---|---|---|---|---|---|---|
| 1U4J | *Bungarus caeruleus* | Indian krait | Elapidae | Group I | 118 | 2.18A | (80) |
| 2WQ5 | *Naja naja* | Indian cobra | Elapidae | Group I | 119 | 1.65A | (49) |
| 2QHD | *Echis carinatus* | Sindh saw-scaled viper | Viperidae | Group IIB | 122 | 1.95A | (48) |
| 4QEM | *Daboia russelli* | Russell's viper | Viperidae | Group IIA | 121 | 1.20A | (81) |

After prioritizing svPLA2 targets, we analyzed the selected peptides from various resources for molecular docking. Based on intermolecular interactions observed in svPLA2-peptide complexes, we prioritized LTNF10 as our primary peptide candidate. LTNF10 was further optimized to LTNF10-MT by introducing a point mutation at specific positions using positional scanning. We then performed MD simulation in triplicate for all the svPLA_2_ complexes with LTNF10 and LTNF10-MT. The complete workflow of the study is provided in Figure 2.

**Figure 1.**
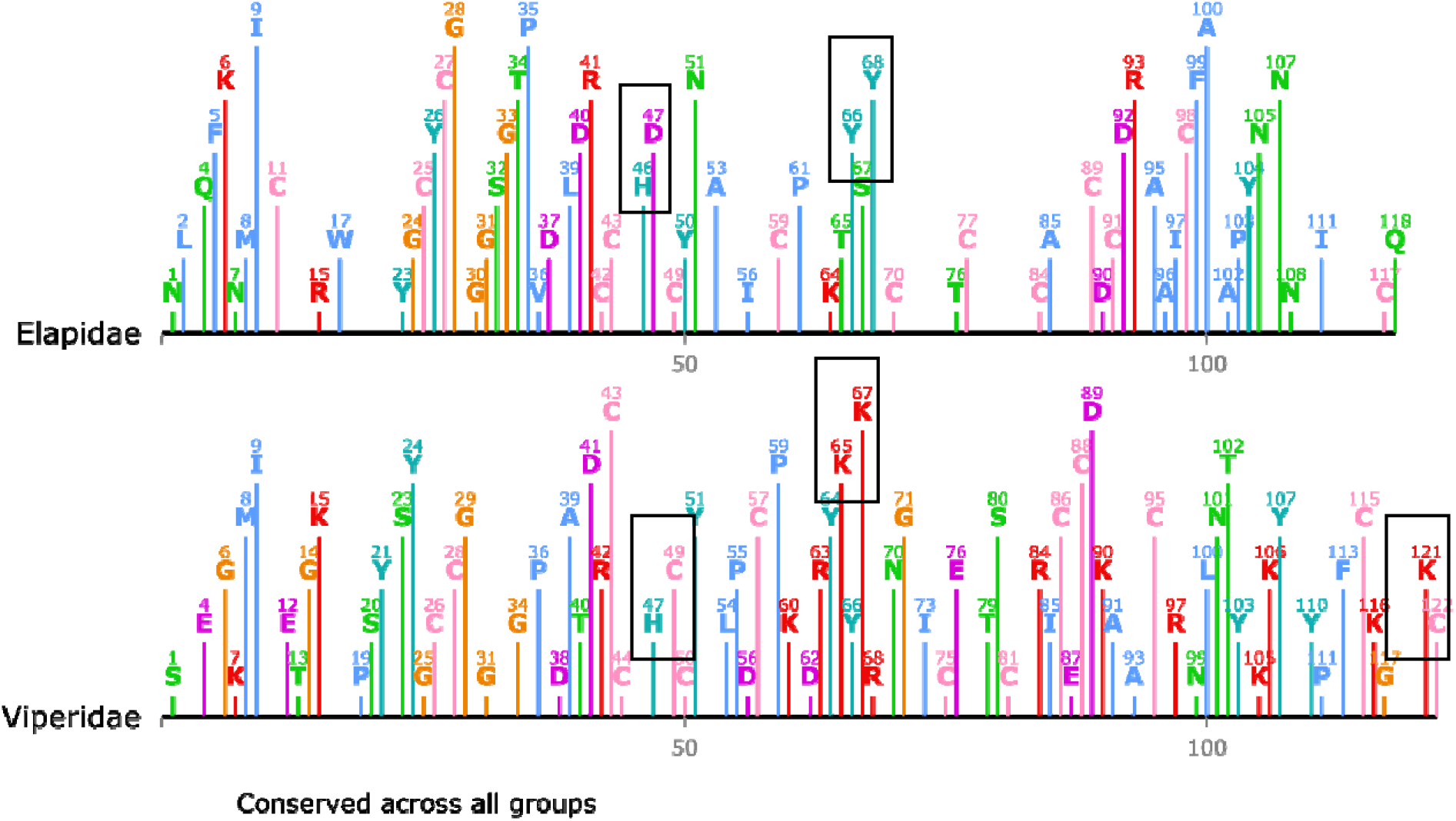
Comparative Evolutionary Profiles of svPLA Sequence Conservation in Elapidae and Viperidae Generated with Alvis(17). Sequence logos of aligned amino acid sequences from Elapidae (A) and Viperidae (B) families. The x-axis indicates amino acid positions, and the y-axis represents information content (bits). The height of each letter corresponds to its relative frequency at that position. The total stack height indicates how conserved that position is across the sequences. In the Elapidae family, we can see His46-Asp47 in the catalytic region, while in the Viperidae family, we see His47-Cys48. This difference can alter the enzymatic activity of svPLA_2_s. We observe Tyr66-Ser67-Tyr68 in Elapidae, while in Viperidae we see Lys65-Tyr66-Tyr67. This difference is attributed to the viper’s non-enzymatic membrane-penetrating activity. We also observe Lys121-Cys122 at the end in Viperidae, making its sequence longer than Elapidae, which also helps in anchoring to the target membrane.

**Figure 2.**
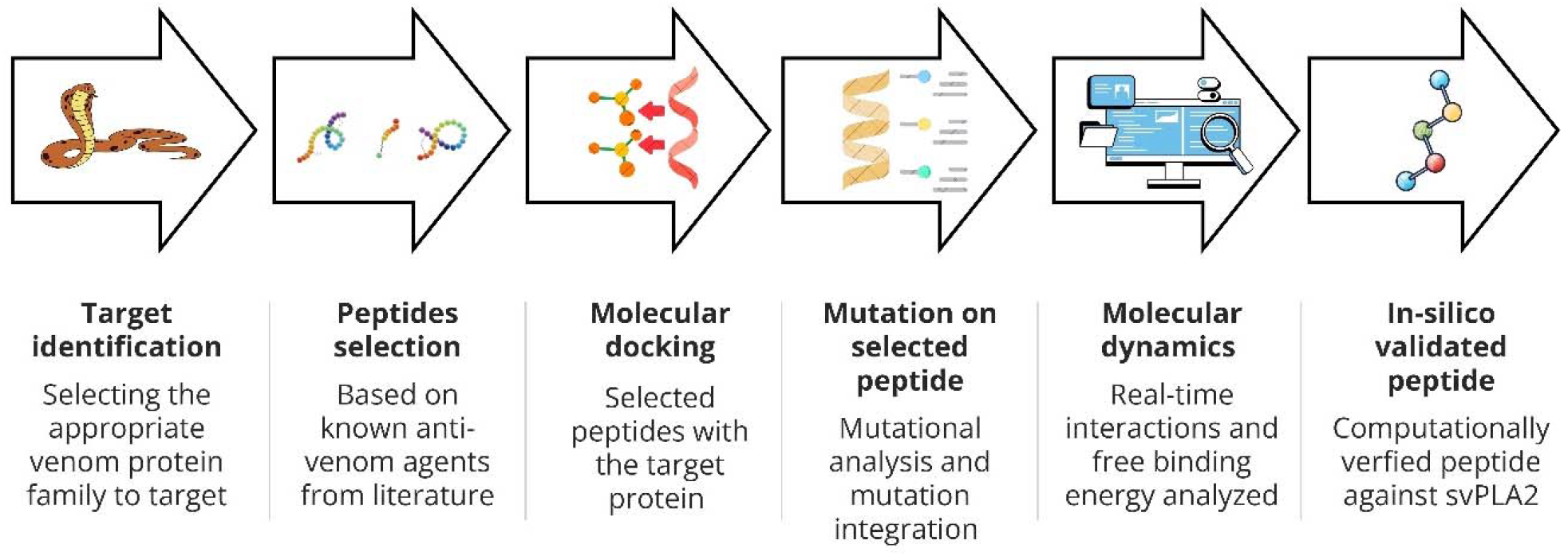
Computational workflow for peptide design against svPLA proteins. The schematic representation outlines the stepwise strategy employed to identify and optimize peptide inhibitors targeting the svPLA protein family. The process begins with target identification of svPLA enzymes, followed by selection of candidate peptides from literature reports of antivenom agents. These peptides are subjected to molecular docking to assess binding interactions with the target proteins. Selected candidate undergoes mutational analysis to improve stability and affinity, which are then evaluated through molecular dynamics simulation to capture real-time interactions and estimate binding free energies. The pipeline culminates in the identification of an in-silico validated peptide, computationally verified for its potential to neutralize svPLA activity. This integrative approach highlights the synergy of bioinformatics, docking, mutagenesis, and dynamics in rational peptide design for antivenom development.

### Physicochemical and structural analysis of selected peptide candidates

Among the peptide candidates, LTNF10 consistently exhibited the most favorable binding across all svPLA_2_-LTNF10 complexes, with affinity scores ranging from –6.4 to –8.2 kcal/mol, suggesting robust interaction with both svPLA_2_ from Elapids and Vipers. LTNF15 showed moderate binding, with weak or unfavorable scores against Elapids but improved affinity against Viperids (–2.9 and –3.5 kcal/mol). TP1 and TP2 displayed variable affinities, with TP1 showing strong binding to 2QHD (–6.4 kcal/mol) but weak interactions elsewhere, while TP2 bound moderately to 4QEM (–4.6 kcal/mol). SP1 and SP2 generally exhibited poor binding, with positive or near zero scores indicating weak or unstable complexes. JB006-free (82) consistently yielded positive scores (7.2–19.9 kcal/mol), reflecting poor binding and steric incompatibility due to its extended length. Overall, these observations highlight LTNF10 as the most promising peptide candidate, with consistent negative affinity scores across all svPLA enzymes, supporting its potential as a broadly effective peptide inhibitor.

Computational prediction showed LTNF10 is consistently non toxic and non hemolytic (a seen in Table 3), in contrast to other peptide candidates that displayed hemolytic potential. Thi favorable safety profile is key for translational potential, as it minimizes the risk of off target effects.

**Table 2.** Affinity scores (kcal/mol) of peptide candidates docked to svPLA from Elapidae (1U4J, 2WQ5) and Viperidae (2QHD, 4QEM) using AutoDock CrankPep (ADCP). The Table summarizes the predicted affinity scores of seven peptide candidates against four representative svPLA structures, with more negative affinity score values indicating stronger predicted binding.

| **S. No.** | **Peptide candidate** | **Sequence length** | **Affinity score from ADCP (kcal/mol)** |  |  |  |
|---|---|---|---|---|---|---|
|  |  |  | **1U4J** | **2WQ5** | **2QHD** | **4QEM** |
| 1 | LTNF15 | 15 | 4.7 | 1.8 | -2.9 | -3.5 |
| 2 | LTNF10 | 10 | -6.8 | -6.4 | -8.2 | -7.3 |
| 3 | TP1 | 15 | 0.7 | -1.2 | -6.4 | -0.5 |
| 4 | TP2 | 15 | -1.1 | 2.6 | 0.8 | -4.6 |
| 5 | SP1 | 15 | 2.2 | 6.3 | 8.6 | -1.2 |
| 6 | SP2 | 15 | 10.1 | 4.3 | 5.2 | 4.8 |
| 7 | JB006-free | 20 | 11.1 | 15.9 | 7.2 | 19.9 |

**Table 3.**
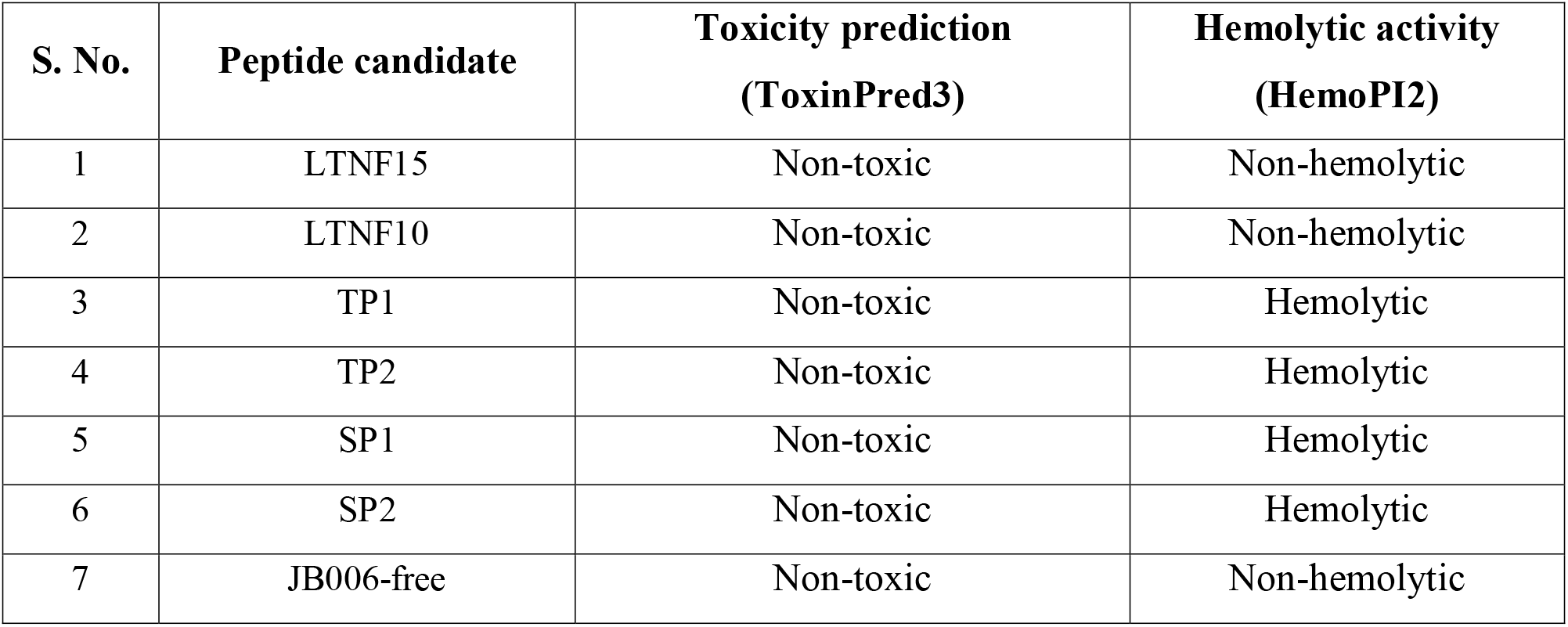
Predicted toxicity and hemolytic activity profiles of peptide candidates assessed using ToxinPred3 and HemoPI2.

The predicted three-dimensional structure of the LTNF10 peptide candidate from trRosetta i shown in Figure 3. An electrostatic interaction is observed between the positively charged side chain of Lys2 and the negatively charged side chain of Asp5, suggesting a stabilizing salt bridge within the short chain. These features highlight charge complementarity, and proline induced conformational constraints might influence the LTNF10 peptide structural stability and potential binding with svPLA_2_.

**Figure 3.**
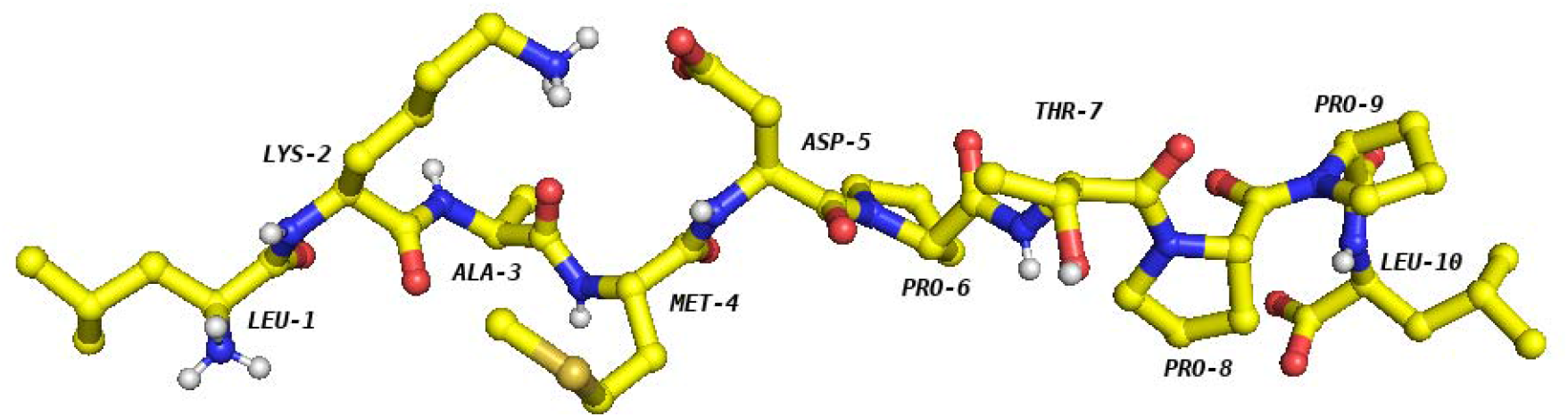
Structural representation of LTNF10 highlighting intramolecular interactions between Lys-2 and Asp-5.

Molecular docking studies show LTNF10 interacting with critical residues present in svPLA_2,_ shown in Figures 4-7, forming stabilizing molecular contacts. Unlike longer peptides derived from turmerin (TP1 and TP2), snaking-z (SP1 and SP2), and JB006-free, which may introduce structural incompatibility with svPLA_2_, LTNF10, with a small length and compact 3D structure, is easier to optimize for enhanced molecular interactions with svPLA_2_. (Figures S1-S19)

As shown in Figures 4-7, LTNF10 forms non-covalent interactions with critical binding hotspots in svPLA_2,_ which consists of a hydrophobic channel formed by Leu2, Ile9, Trp31, Tyr52 & Asn52/53, a calcium binding loop formed by Gly30–Gly33 & Tyr28, membrane anchoring residues consisting of Lys56, Lys69 & Tyr73 and the catalytic region of His48 & Asp49. In LTNF10 Leu1 forms hydrogen bond with Asp53 (Figure 4) and hydrophobic contact with Trp19 (Figure 6); Lys 2 forms hydrophobic contacts with Asn52 (Figure 5) & Asp49 (Figure 7); Ala3 forms hydrogen bonds with Lys69 (Figure 4) & Asn53 (Figure 6) and hydrophobic contact with Asn52 (Figure 5); Pro6 forms hydrogen bond with Gly32 (Figure 5) and hydrophilic contacts with His48 (Figure 4) & Asn (Figure 6); Pro8 forms hydrogen bond with Gly30 (Figure 4) and hydrophobic contact with Lys69 (Figure 7); Pro9 forms hydrogen bond with Tyr25 (Figure 4) and hydrophobic contact with Asp49 (Figure 7); and Leu10 forms hydrogen bond with Tyr25 (Figure 4) and hydrophobic contact with Gly30 (Figure 7). While these residues from LTNF10 have been shown to interact with significant residues from svPLA_2_, Met4, Asp5, and Thr7 in LTNF10 show low interactions and hydrophobic contacts with the significant residues in svPLA_2_. These three residues are centrally positioned within the binding arc and consistently observed in proximity to svPLA_2_ residues such as Gly31, Ser/Asp49, His48, Tyr52 and Lys69 (Figures 4-7). These contacts are weaker and more variable compared to the terminal anchors, indicating that positions 4, 5, and 7 are structurally permissive and viable for mutation. Their location allows side chain substitutions to potentially enhance electrostatic complementarity, introduce new hydrogen bonds, or reinforce hydrophobic packing without disrupting the native scaffold. Hence, Met4, Asp5 and Thr7 positions show potential for fine tuning the LTNF10-svPLA_2_ interface and improving intermolecular interactions. Therefore, with the favorable therapeutic profile, originating from a natural protein and favorable molecular interactions, LTNF10 was prioritized as the primary peptide candidate.

**Figure 4.**
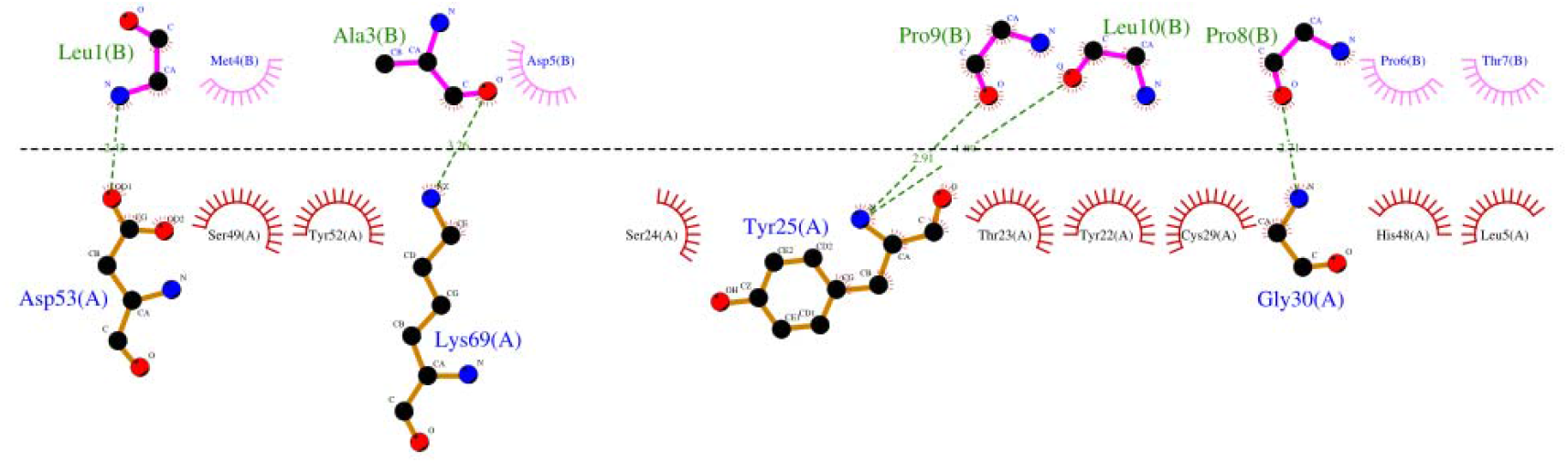
Hydrogen bonds and Hydrophobic contacts Stabilizing *Bungarus caeruleus* (1U4J) svPLA (Chain A)–LTNF10 (Chain B) Complex, highlighting Asp53(A)–Leu1(B), Lys69(A)-Ala3(B), Tyr25(A)-Pro9(B)/Leu10(B) and Gly30(A)-Pro8(B) hydrogen bonds and other depictions shows hydrophobic Interactions.

**Figure 5.**
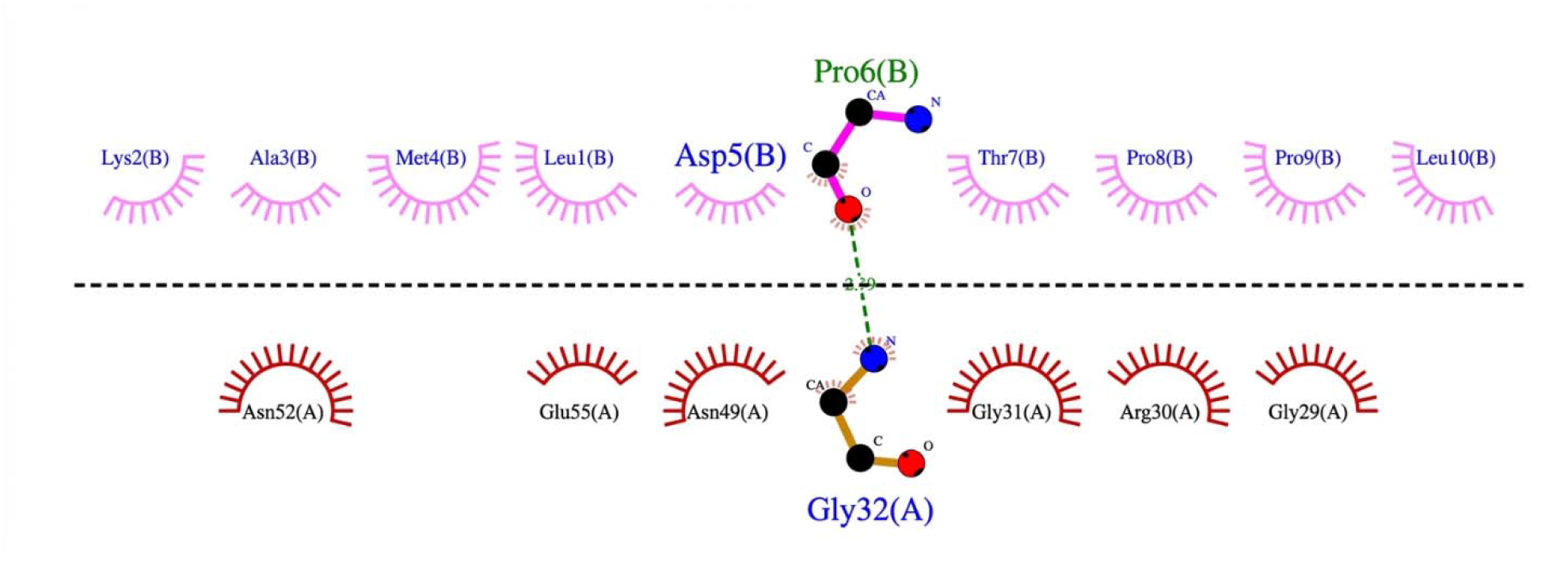
Hydrogen bonds and Hydrophobic contacts stabilizing *Naja naja* (2WQ5) svPLA (Chain A)–LTNF10 (Chain B) Complex, highlighting Gly32(A)-Pro6(B) hydrogen bond, with additional depictions of hydrophobic interactions contributing to peptide binding.

**Figure 6.**
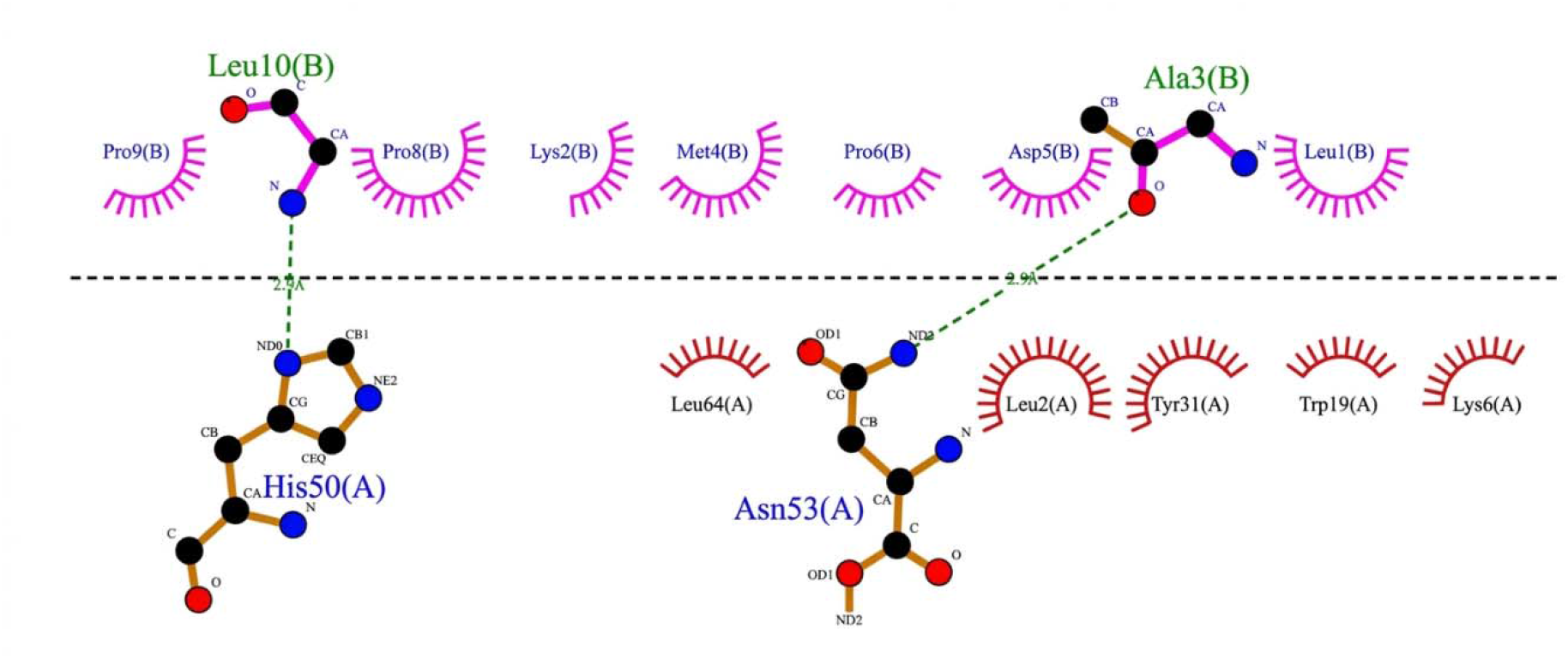
Hydrogen bonds and Hydrophobic contacts stabilizing *Echis carinatus* (2QHD) svPLA (Chain A)–LTNF10 (Chain B) Complex, highlighting His50(A)–Leu10(B) and Asn53(A)-Ala3(B) hydrogen bonds and other depictions show hydrophobic Interactions.

**Figure 7.**
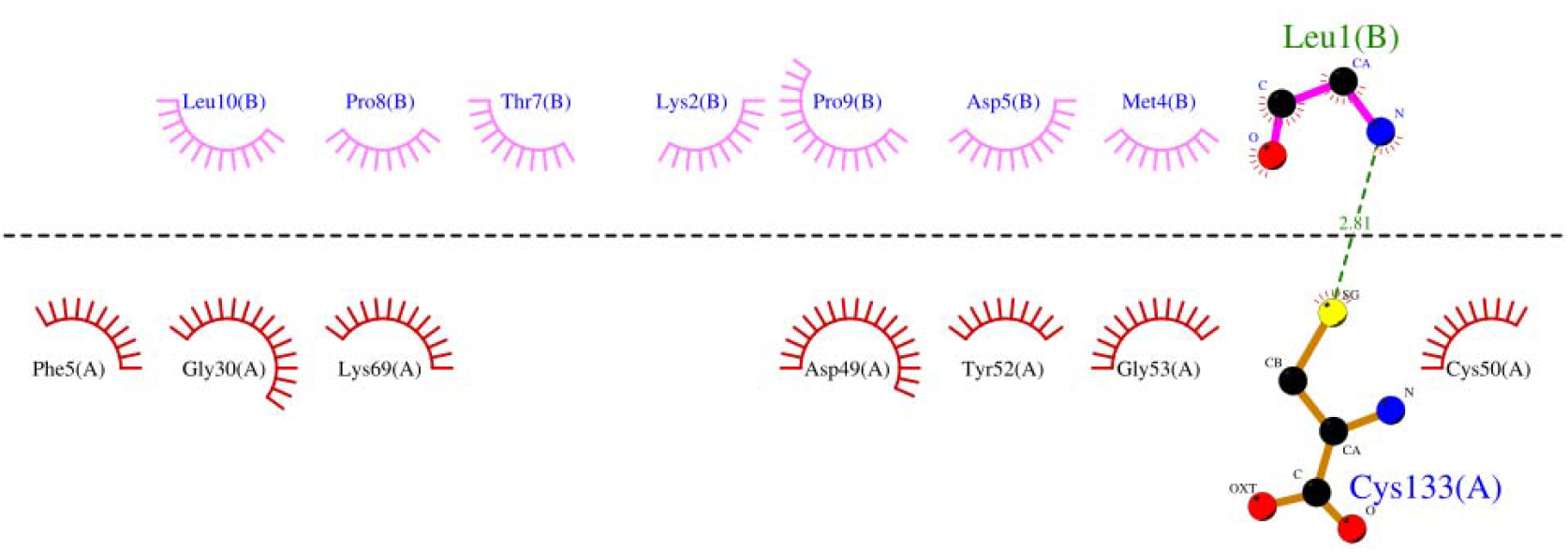
Hydrogen bonding and Hydrophobic contacts stabilizing *Daboia russelii pulchella* (4QEM) svPLA (Chain A)–LTNF10 (Chain B) Complex, highlighting Cys133(A)–Leu1(B) hydrogen bond and other depictions show hydrophobic Interactions.

### LTNF10 Optimization

To determine the suitable mutations in LTNF10, positional scanning was performed using FoldX (51) (Table S3), and identified stabilizing mutations were re-assessed using MutaBind2 (53) and mCSM-PPI2 (52). Significant point mutations identified in LTNF10 and its corresponding changes in protein-peptide complex stability (FoldX) or binding affinity (MutaBind2 and mCSM-PPI2) due to point mutations are shown in Figure 8.

**Figure 8.**
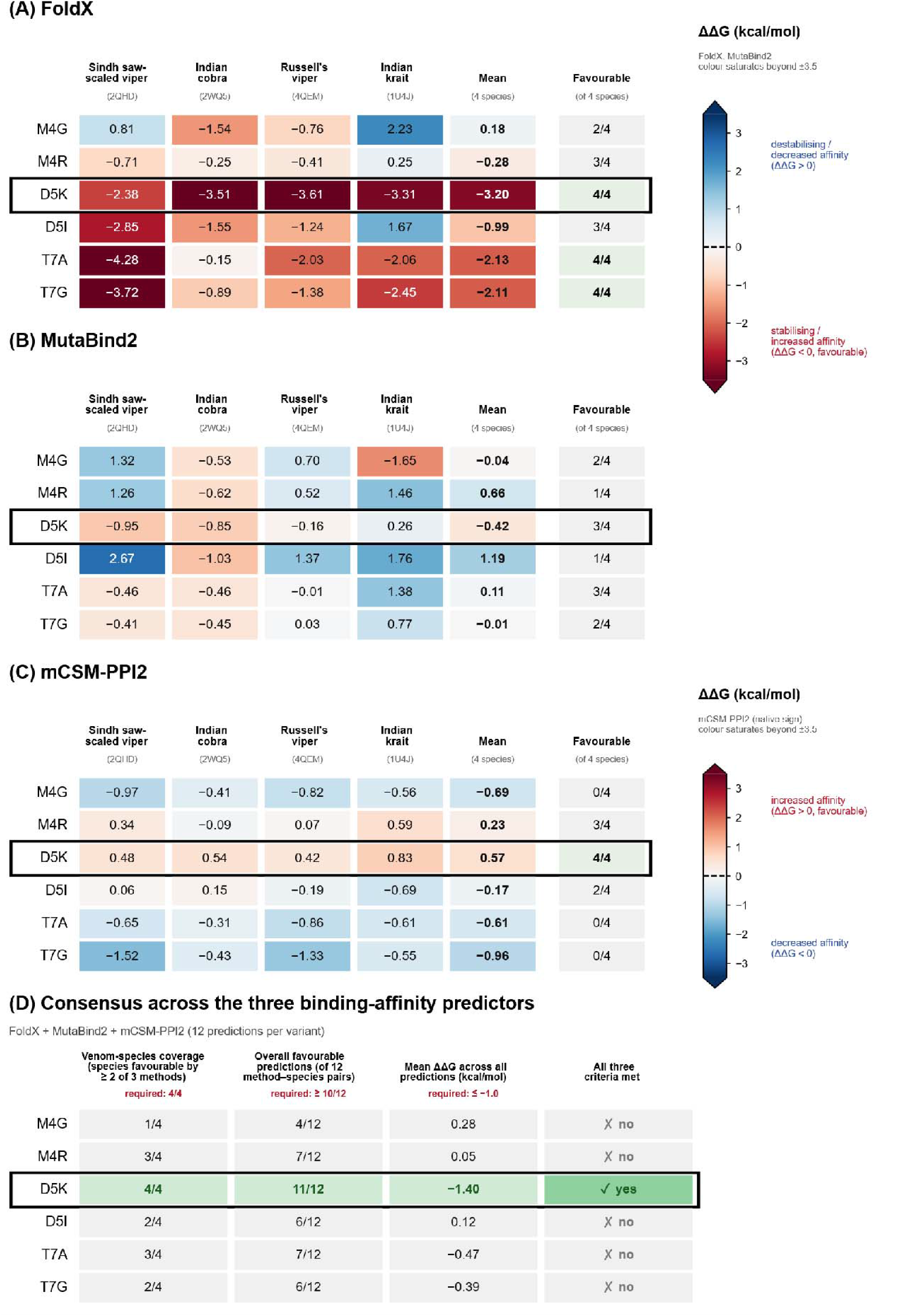
Heatmaps showing the effects of point mutations (M4G, M4R, D5K, D5I, T7A, and T7G) in LTNF10 on the stability and binding of the four target systems for each snake species svPLA_2_ with LTNF10. (A) FoldX, (B) MutaBind2 and (C) mCSM-PPI2 ΔΔG values for each mutation against the four svPLA targets, with the mean over the four targets and the number of targets with a favourable prediction. For FoldX and MutaBind2, negative values are favourable; mCSM-PPI2 positive values are favourable (separate colour scale). Red indicates favourable and blue unfavourable binding in all panels; (D) Consensus across the three predictors (with mCSM-PPI2 values sign-inverted so that negative ΔΔG is favourable for all methods).

ΔΔG predictions were computed to capture the effects of point mutations in LTNF10 on the stability and binding of the four target systems for each species svPLA_2_ with LTNF10 were made using three computational methods: FoldX (51), MutaBind2 (53), and mCSM-PPI2 (52) to capture differences in mutational impact on LTNF10-svPLA_2_ complexes. For FoldX positional scan results (ΔΔG), the standard threshold for a significant change in protein stability is ± 1.0 kcal/mol (51). The MutaBind2 server classifies a mutation as deleterious if ΔΔ G ≥ 1.5 or ≤ −1.5 kcal/mol (53). mCSM-PPI2 classifies a mutation as stabilizing if ΔΔ G ≥ 0.5 kcal/mol, as destabilizing if ΔΔ G ≤ −0.5 kcal/mol, and neutral if ΔΔ G ≤ 0.5 kcal/mol and ΔΔ G ≥ −0.5 kcal/mol (52). The threshold of −1 kcal/mol was chosen empirically for the mean calculated across the three methods after analysing the distribution of the computed values.

The negative values indicate favourable effects of mutation in FoldX and MutaBind2, while mCSM-PPI2 uses the opposite sign convention for a favourable change in binding affinity (ΔΔG from mCSM-PPI2 is inverted for the consensus analysis in Figure 8D for consistency in comparison across methods). A mutation was prioritized if it met the following criteria: (i) venom-species coverage, which means the number of venom species (out of four) for which at least two of the three methods predict a favourable effect; (ii) ΔΔG is favourable in at least 80% of the 12 predictions (≥10/12); and (iii) a mean ΔΔG ≤ −1.0 kcal/mol across all 12 predictions from the three methods used (Figure 8D).

The point mutations in LTNF10 show different ΔΔG patterns across the three methods used. For the M4G, FoldX and MutaBind2 values were observed to be favourable only against the Indian cobra (Figure 8A and 8B), while mCSM-PPI2 predicted reduced affinity at all four targets (Figure 8C). Therefore, the M4G mutation in LTNF10 did not qualify for the criterion (Figure 8D). For M4R, FoldX and mCSM-PPI2 were mildly favourable at three targets, but MutaBind2 predicted weaker binding at three of the four targets (Figure 8A-C). Therefore, the M4R mutation in LTNF10 did not qualify for the criteria (Figure 8D).

The T7A and T7G mutations reported favourable outcomes in FoldX, but the other two methods did not predict it to be favourable (Figure 8). FoldX values were favourable for all four targets for both mutations (Figure 8A). MutaBind2 was favourable for T7A at three targets but predicted weaker binding against the Indian krait, and for T7G, it was favourable only against the Sindh saw-scaled viper and the Indian cobra, with relatively weaker binding against Russell’s viper and the Indian krait (Figure 8B). mCSM-PPI2, however, predicted reduced affinity for all four targets for both mutations (Figure 8C). As a result, T7A had 3/4 species qualifying the criteria (i) (all except the Indian krait) and T7G at only 2/4 (Sindh saw-scaled viper and the Indian cobra) (Figure 8D). Overall, they had 7/12 and 6/12 favourable predictions, with mean ΔΔG across methods to be −0.47 and −0.39 kcal/mol (Figure 8D). Despite the large FoldX values, neither mutation met any of the stated criteria (i)–(iii).

Both D5K and D5I mutations were observed to have favourable FoldX values (Figure 8A), indicating enhanced stability, while mCSM-PPI2 did not support this stabilizing trend for D5I. D5I was favourable in FoldX against the two vipers and the Indian cobra but not the Indian krait (Figure 8A). MutaBind2 was favourable only against the Indian cobra and predicted weaker binding at the other three targets, and mCSM-PPI2 was marginally neutral against the Sindh saw-scaled viper and the Indian cobra (Figure 8B and 8C). D5I therefore did not qualify the criterion with 2/4 species qualifying, 6/12 favourable predictions and a mean ΔΔG of +0.12 kcal/mol across all 12 predictions (Figure 8D). D5K stood out from the rest of the candidates, with FoldX (Figure 8A) and mCSM-PPI2 (Figure 8C) consistently indicating neutral to favourable changes in stabilization, while MutaBind2 reported favourable ΔΔG at three targets (Figure 8B) and slightly unfavourable for Indian krait (Figure 8B). The divergence observed in MutaBind2 can be attributed to the inclusion of evolutionary substitution and continuum desolvation penalties (53), which penalize non-conservative charge reversals (Asp to Lys in case of D5K mutation in LTNF10) even when favourable outcomes were observed in FoldX and mCSM-PPI2. D5K was the only mutation candidate meeting all three criteria (4/4 species qualified, 11/12 predictions, mean −1.40 kcal/mol; Figure 8D).

In svPLA_2_, since Asp49 is a key residue in the binding region (Figure 14), making aspartate to lysine mutation in LTNF10 seems a favourable choice due to its central position in the LTNF10 sequence and electrostatic complementarity with Asp49. The positive charge of lysine in LTNF10 and the negative charge of Asp49 in svPLA_2_ will favour strong electrostatic interaction. This suggests that introducing a lysine at position 5 enhances electrostatic complementarity in elapids while simultaneously reinforcing viper toxin stability. Therefore, these results highlight D5K as the most rational mutation, supported by changes in stabilization score and affinity scores across LTNF10-svPLA_2_s complexes from both families Elapidae and Viperidae. After this validation, we optimized LTNF10 to LTNF10-MT by introducing a mutation at position 5 of LTNF10, which is Asp5 to Lys5 (D5K).

### Binding potential of LTNF10 and LTNF10-MT to svPLA_2_ from Elapidae and Viperidae

The binding free energy results from MD simulations (Table 4) highlight consistent differences in binding free energy between LTNF10-svPLA_2_ and LTNF10-MT-svPLA_2_ complexes, with LTNF10-MT (Indian cobra and Indian krait) showing markedly stronger binding free energy in Elapidae, while exhibiting reduced binding free energy in Viperidae when compared to LTNF10. These findings emphasise species specific variation in mutational effects on svPLA binding with the candidate.

**Table 4.** Binding free energy calculations (kJ/mol) for LTNF10 and LTNF10-MT with svPLA complexes by MMPBSA. This table presents MMPBSA calculations using MD simulation trajectories with averaged binding free energies for LTNF10 and LTNF10-MT with svPLA from four snake species: Indian cobra (2WQ5), Indian krait (1U4J), Sindh saw scaled viper (2QHD), and Russell’s viper (4QEM). Binding free energy values are in kJ/mol.

| **Species/PDB ID** | **LTNF10** |  |  |  | **LTNF10-MT** |  |  |  |
|---|---|---|---|---|---|---|---|---|
|  | Rep 1 | Rep 2 | Rep 3 | Average | Rep 1 | Rep 2 | Rep 3 | Average |
| **1U4J (Indian krait)** | -166.766 ± 84.228 | -124.524 ± 54.647 | -130.238 ± 55.330 | -140.509 ± 64.707 | -457.147 ±61.942 | -493.751 ±59.993 | -463.751 ±59.498 | -471.216 ± 61.227 |
| **2WQ5 (Indian cobra)** | -215.976 ± 48.762 | -214.152 ± 43.759 | -293.279 ± 43.592 | -241.13 ± 45.371 | -357.003 ± 62.412 | -420.435 ± 63.165 | -395.712 ± 57.779 | -390.716 ± 61.118 |
| **2QHD (saw-sindh viper)** | -699.383 ± 75.607 | -627.223 ± 88.064 | -794.262 ±61.727 | -706.956 ± 75.132 | -573.115 ± 81.874 | -539.10 ± 70.751 | -652.123 ± 77.630 | -588.112 ± 76.751 |
| **4QEM (Russell’s viper)** | -746.319 ± 46.029 | -668.978 ± 35.565 | -688.26 ± 44.538 | -701.185 ± 42.044 | -487.367 ± 63.476 | -418.498 ± 53.328 | -579.541 ± 60.705 | -495.135 ± 59.169 |

The molecular dynamics simulations across all four LTNF10-svPLA_2_ complexes revealed consistent and stable MD trajectory characteristics for svPLA_2_s. This is supported by RMSD (root mean square deviation), RMSF (root mean square fluctuation), and SASA (solvent accessible surface area) values across triplicate MD production runs, while we observed more flexible characteristics in LTNF10-MT peptides when bound with Elapidae (Figures S20-23 and Tables S4-S7).

We can see the contributions of significant amino acid residues in svPLA_2_s from Elapidae (top) and Viperidae (bottom) for both LTNF10 (green) and LTNF10-MT (purple) in Figure 11. Decomposition energy contributions by all possible significant residues from svPLA2s, along with LTNF10 and LTNF10-MT residue contributions, can be referred to in Tables S8-S15. Intermolecular interactions in Elapidae complexes are shown in Figure 9. In LTNF10-svPLA_2_ complexes from Elapidae (1U4J and 2WQ5), the point mutation significantly reshaped the interaction profile. In 1U4J svPLA_2_, the catalytic residues Tyr31 and Asp49 act as favorable contributors in LTNF10-MT (–11.01 and –183.06 kJ/mol) as compared to LTNF10 (–5.11 and – 42.99 kJ/mol), while the Lys66 residue, important for anchoring, shows unfavorable interactions (+61.35 vs. +1.77 kJ/mol) (Table S9). The LTNF10 dynamics revealed that Asp5 contributes unfavorably (+62.07 kJ/mol) and, upon mutation to Lys5 in LTNF10-MT, showed a favorable contribution (–67.51 kJ/mol). The Leu1 residue contribution shifted from destabilizing (+20.10 kJ/mol) to favorable (–57.51 kJ/mol) upon Asp5 to Lys5 mutation in LTNF10-MT (Table S12). In hydrogen bond analysis (Table S16), we observed shifts in energetic contribution, which is supported by stronger and more persistent hydrogen bonds in LTNF10-MT, where Lys5 forms hydrogen bonds with Tyr31, Asn53, and Asp49, and Leu1 forms a hydrogen bond with Tyr28, Cys29, and Asp49.

**Figure 9.**
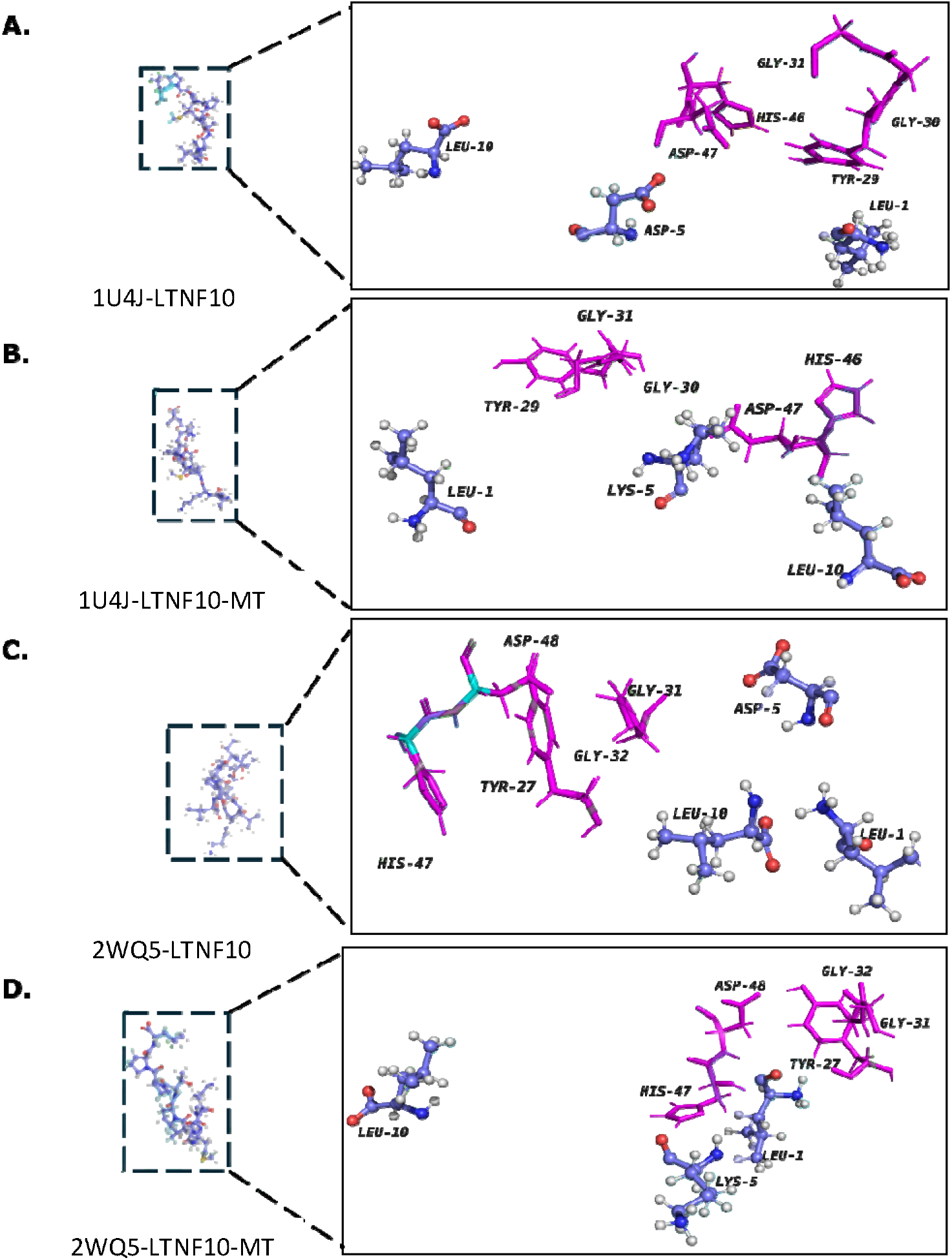
Structural comparison of LTNF10-svPLA_2_ complexes showing LTNF10 interactions in Elapidae complexes (1U4J and 2WQ5), highlighting key residues in LTNF10 and LTNF10-MT. Important residues from svPLA_2_ are highlighted in magenta, while key residues from LTNF10 and LTNF10-MT are highlighted in purple, represented by sticks and balls. A) shows the interactions between 1U4J svPLA_2_ and LTNF10; B) shows the corresponding mutant LTNF10-MT interaction with 1U4J svPLA_2_; C) shows the interactions between 2WQ5 svPLA_2_ and LTNF10; D) shows the corresponding mutant LTNF10-MT interactions with 2WQ5 svPLA_2_, revealing differences in residues contributing to intermolecular interactions between LTNF10-svPLA_2_ and LTNF10-MT-svPLA_2_.

**Figure 10.**
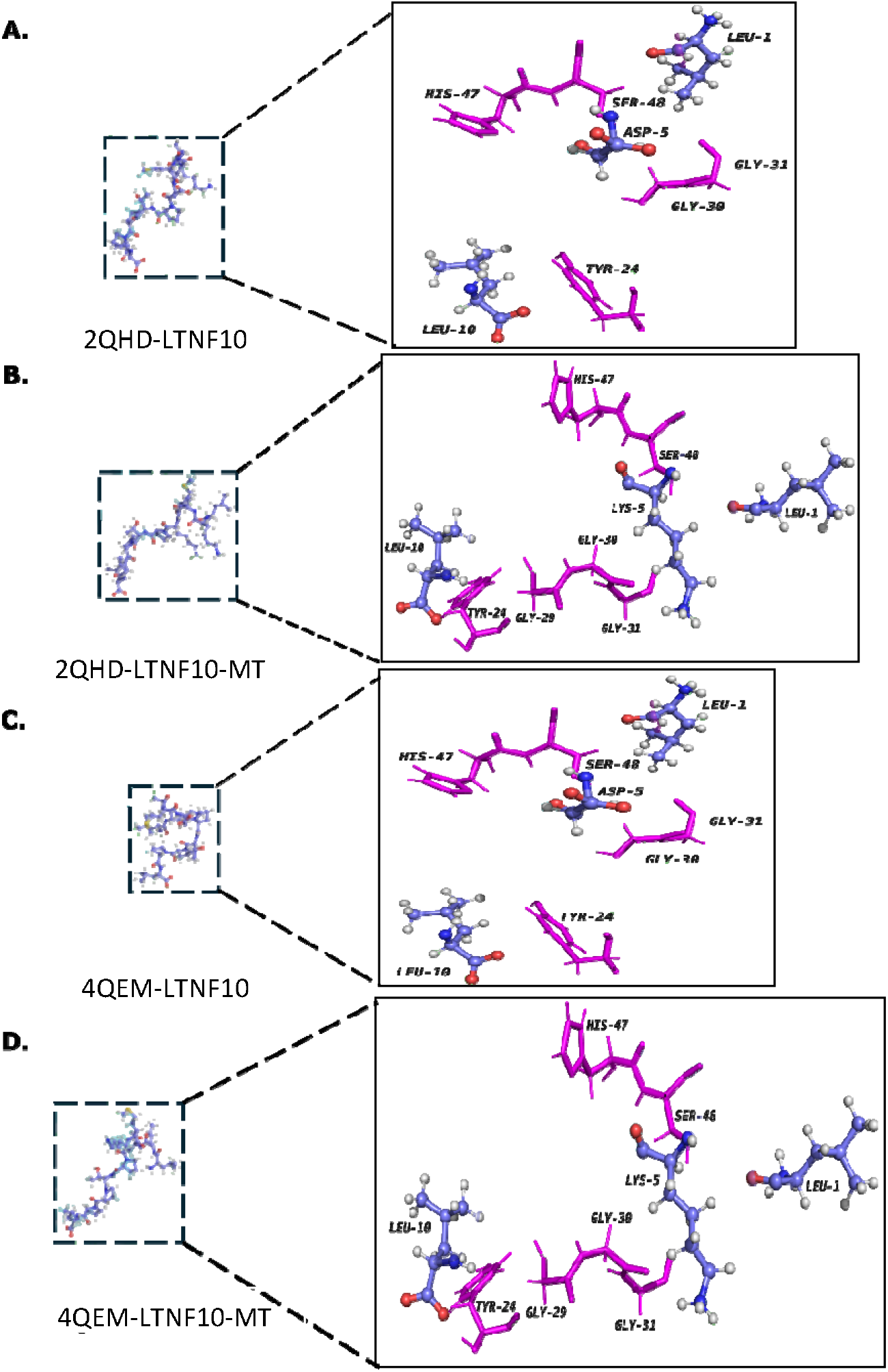
Structural comparison of LTNF10-svPLA_2_ complexes showing LTNF10 interactions in Viperidae complexes (2QHD and 4QEM), highlighting key residues in LTNF10 and LTNF10-MT. Important residues from svPLA_2_ are highlighted in magenta, while key residues from LTNF10 and LTNF10-MT are highlighted in purple, represented by sticks and balls. A) show the interactions between 2QHD svPLA_2_ and LTNF10; B) shows the corresponding mutant LTNF10-MT interaction with 2QHD svPLA_2_; C) shows the interactions between 4QEM svPLA_2_ and LTNF10; D) shows the corresponding mutant LTNF10-MT interactions with 4QEM svPLA_2_, revealing differences in residues contributing to intermolecular interactions between LTNF10-svPLA_2_ and LTNF10-MT-svPLA_2_.

**Figure 11.**
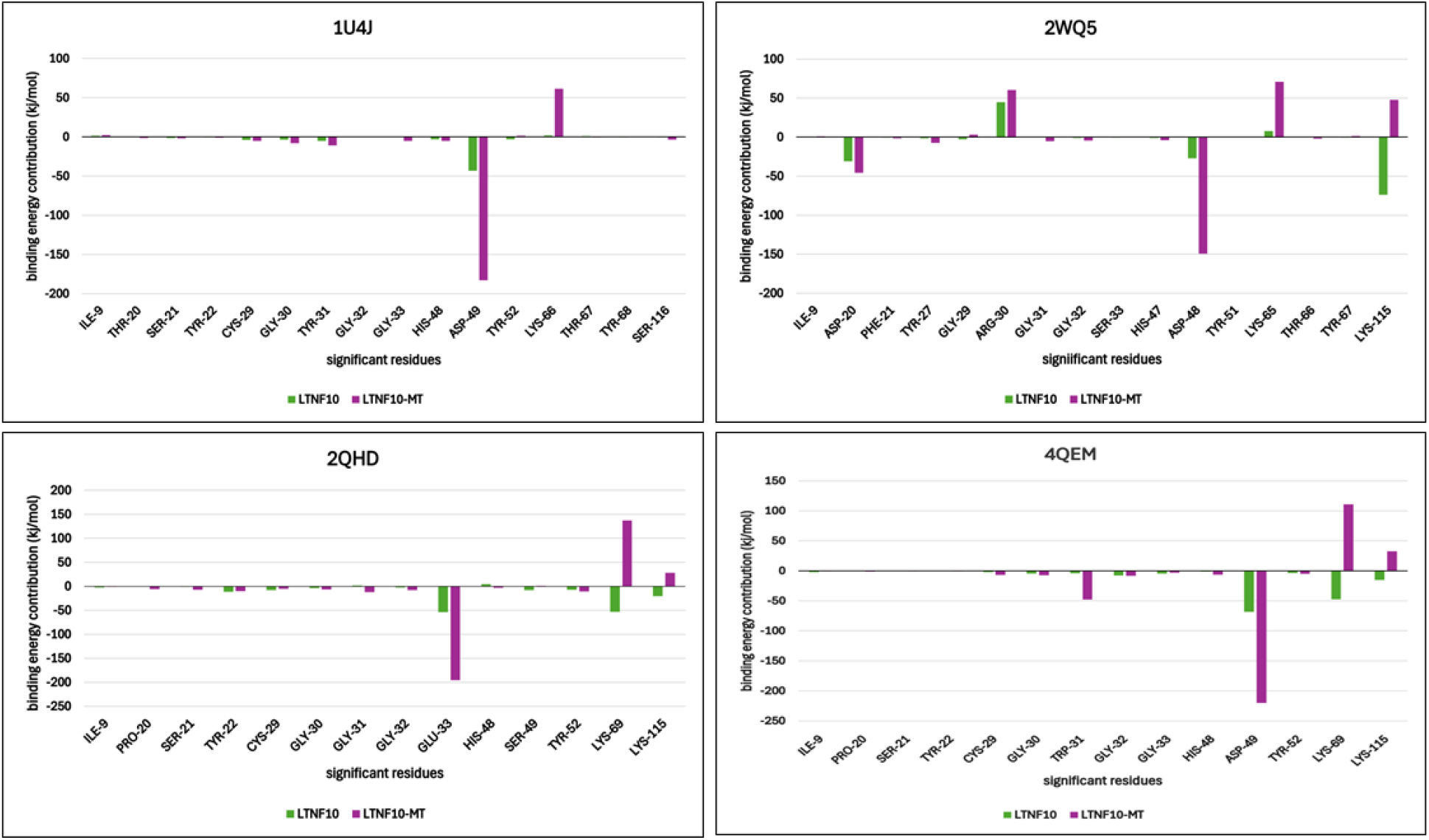
Residue-specific Binding Energy Contributions in LTNF10-svPLA and LTNF10-MT-svPLA_2_ complexes of Elapidae (1U4J and 2WQ5) and Viperidae (2QHD and 4QEM). Decomposition energy for significant amino acid residues is depicted in green for the svPLA_2_ complex with LTNF10 and purple for the svPLA_2_ complex with LTNF10-MT.

In 2WQ5 svPLA_2_, Asp48 contribution increased substantially (–27.27 to –149.11 kJ/mol), while Lys65 lost stabilizing potential (71.15 vs. 7.74 kJ/mol) upon mutation to LTNF10-MT (Table S9). The peptide dynamics support this trend, where Leu1 shows better stabilization (−68.15 to - 188.18 kJ/mol) upon mutation to LTNF10-MT, and Asp5 in LTNF10 mutating to Lys5 in LTNF10-MT shows more favorable contributions (–30.71 vs. +13.44 kJ/mol) (Table S13 and Figures 9-11). Hydrogen bond analysis (Table S17) showed Leu1 in LTNF10-MT engages more consistently with His48, Asp49, and surrounding residues, while Lys5 in LTNF10-MT improved contacts with Asp23, Gly29, and Arg94. Binding free energy calculations using MM/PBSA (Table 4) showed significantly enhanced binding free energy in the mutant variant LTNF10-MT-svPLA_2_ complex from 1U4J (−471.216 kJ/mol) and 2WQ5 (−390.716 kJ/mol) when compared to LTNF10-svPLA_2_ from 1U4J (−140.509 kJ/mol) and 2WQ5 (−241.13 kJ/mol). This trend of enhanced binding profiles is also supported by PRODIGY and Boltz-2 predictions of binding affinities (Table 5).

**Table 5.** Comparative average binding affinities of svPLA s from elapidae and viperidae predicted by PRODIGY and Boltz-2. This table summarises the average binding affinities of LTNF10 and LTNF10-MT with svPLA from elapidae (Indian krait, Indian cobra) and viperidae (Sindh saw-scaled viper, Russell’s viper). Values are reported as kcal/mol.

| Species / PDB ID | **PRODIGY<br>LTNF10<br>Avg (kcal/mol)** | **PRODIGY<br>LTNF10-MT<br>Avg (kcal/mol)** | **Boltz-2<br>LTNF10<br>Avg (kcal/mol)** | **Boltz-2<br>LTNF10-MT<br>Avg (kcal/mol)** |
|---|---|---|---|---|
| **1U4J<br>(Indian krait)** | −7.47 | −9.17 | −9.22 | −9.86 |
| **2WQ5<br>(Indian cobra)** | −8.67 | −10.60 | −8.91 | −9.95 |
| **2QHD<br>(Sindh saw-scaled viper)** | −8.47 | −7.93 | −9.30 | −9.26 |
| **4QEM<br>(Russell’s viper)** | −6.63 | −5.97 | −9.21 | −9.11 |

In contrast, the LTNF10-svPLA_2_ complexes from Viperidae (2QHD and 4QEM) showed little shift and sometimes opposing shifts as well. Intermolecular interactions in Viperidae complexes are shown in Figure 10. In 2QHD svPLA_2_, His48 and Asp49 showed minimal energetic changes, while Glu33 emerged as a significant stabilizer, with its contribution increasing from –55.99 to – 195.31 kJ/mol upon mutation from LTNF10 to LTNF10-MT (Table S10). The peptide dynamics upon mutation to LTNF10-MT (D5K) showed Leu1 becoming strongly stabilizing (−49.2 to – 160.71 kJ/mol), while Leu10 lost much of its stabilizing role (–152.55 to –46.49 kJ/mol), and Lys5, which is the mutated position, was shown to be less favorable than in Elapids (–98.48 to - 10.01 kJ/mol) (Table S14). Hydrogen bond analysis (Table S18) supported these findings, where Leu1, upon D5K mutation in LTNF10-MT, engaged more consistently with Glu33 and Asp53, while Lys5 in LTNF10-MT formed contacts with Lys69 and Gly31.

In 4QEM svPLA_2_, Asp49 and Trp31 became more favorable contributors (–68.55 and –4.14 kJ/mol to –220.10 and –47.91 kJ/mol), highlighting their role in stabilizing the LTNF10 peptide interface. However, Lys69 and Lys115 in svPLA_2_ shifted destabilizing roles (–47.52 and –15.31 kJ/mol to +111.07 and +32.69 kJ/mol), suggesting that the mutation disrupted electrostatic balance in this binding pocket (Table S11). The peptide dynamics revealed Leu1 in LTNF10-MT becoming more stabilizing (–13.59 to –40.64 kJ/mol), Lys2 in LTNF10-MT shifted from an unfavorable contribution (+18.45 kJ/mol) to a favorable contribution (–56.04 kJ/mol), while Lys5 in LTNF10-MT showed an unfavorable contribution (+82.88 kJ/mol) when compared to Asp5 in LTNF10(−130.11) (Figure 10 and Table S15). Hydrogen bond analysis (Table S19) showed reduced persistence of Leu1 and Lys2 in LTNF10-MT with catalytic residues such as Asp49 and His48, while Lys5 in LTNF10-MT gained contacts with Lys69, but these were less stabilizing overall. Binding free energy calculations using MM/PBSA (Table 4) showed a shift from –706.96 to –588.11 kJ/mol in 2QHD and −701.185 to −495.135 kJ/mol in 4QEM, suggesting weaker interactions in LTNF10-MT-svPLA_2_ complexes compared to LTNF10-svPLA_2_ complexes in Viperidae. This result is supported by binding affinity predictions by PRODIGY and Boltz-2 (Table 5), which report a slight decrease in binding affinities.

The PCA-derived free energy landscapes presented in Figure 12 illustrate the conformational diversity of LTNF10 and LTNF10-MT with svPLA s from Elapidae. For the wild-type LTNF10 complexes (1U4J-LTNF10 and 2WQ5-LTNF10), the landscapes reveal relatively stable low-energy basins, suggesting that LTNF10 shows limited conformational freedom in the binding site of svPLA_2_. In contrast, the mutant LTNF10-MT complexes (1U4J-LTNF10-MT and 2WQ5-LTNF10-MT) display much broader low-energy regions, which suggests enhanced conformational flexibility. Such observations imply that the mutation in LTNF10 promotes structural flexibility while maintaining strong intermolecular interactions with svPLA_2_. This might help LTNF10-MT peptide to dynamically stabilize key contacts with catalytic residues and the calcium-binding loop of svPLA s from Elapidae.

**Figure 12.**
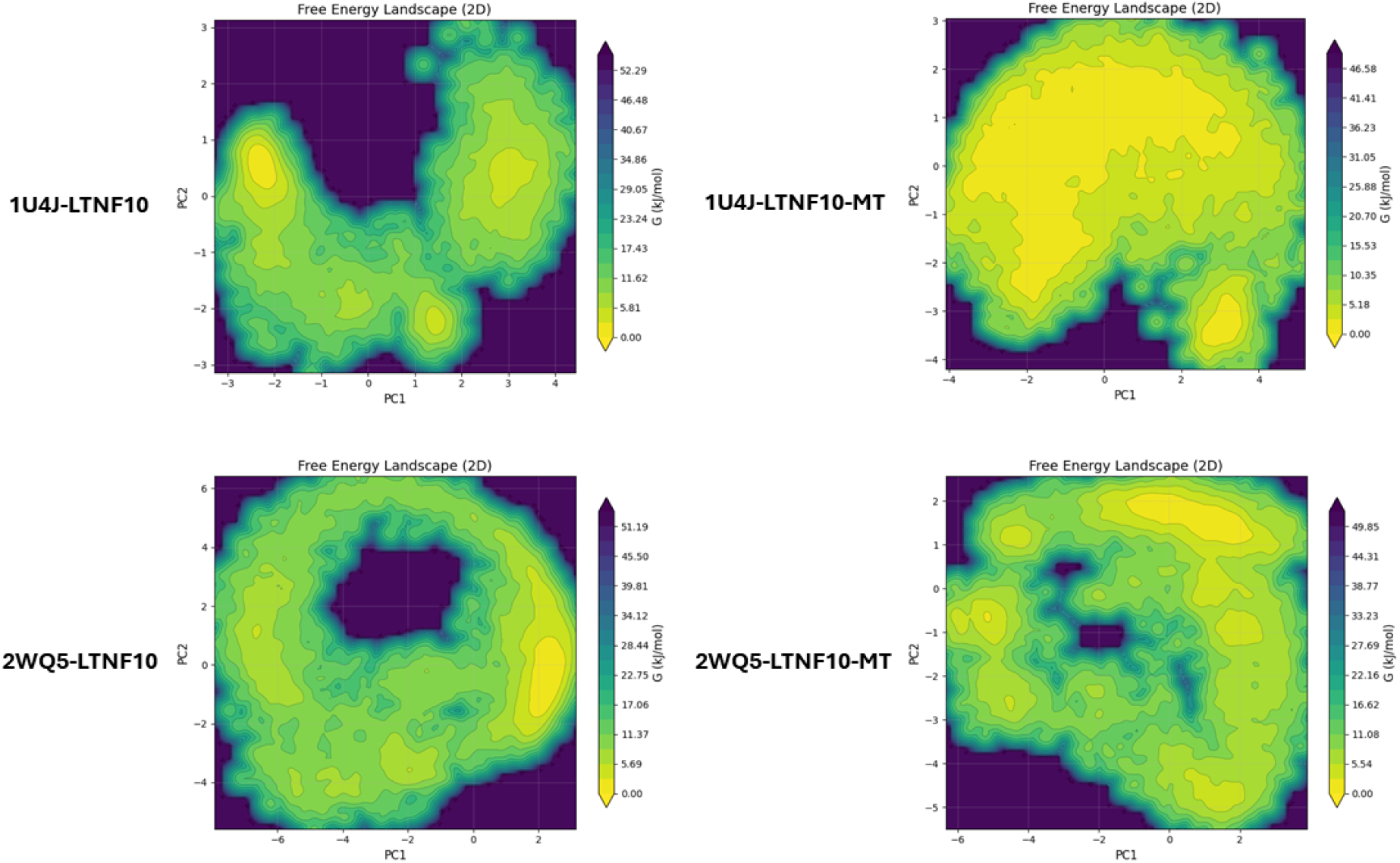
PCA (Principal component analysis) Derived Two Dimensional Free Energy Landscapes (ΔG, kJ/mol) of LTNF10 and LTNF10 MT Peptides from Elapidae. It shows conformational dynamics for LTNF10 and LTNF10 MT from Elapidae. The two dimensional free energy landscapes are plotted along the first two principal components (PC1 and PC2), with contour maps illustrating conformational basins and transitions observed during MD simulations. The color scale represents the relative free energy (ΔG, kJ/mol), ranging from low energy states (yellow) to high energy states (dark purple). The top row shows Free energy landscapes for the 1U4J with LTNF10 (left) and LTNF10 MT (right). The bottom row shows Free energy landscapes for the 2WQ5 with LTNF10 (left) and LTNF10 MT (right). Comparison between LTNF10 and LTNF10-MT peptides highlights shifts in conformational flexibility and energy minima, reflecting the impact of mutation on LTNF10–svPLA interactions from Elapidae.

The PCA-derived free energy landscapes presented in Figure 13 illustrate the conformational diversity of the LTNF10 and LTNF10-MT with svPLA s from Viperidae. They reveal a distinct impact of mutation in LTNF10 on conformational stability and binding energetics. In the LTNF10 complexes (2QHD-LTNF10 and 4QEM-LTNF10), the landscapes are characterized by relatively wider low-energy basins, suggesting enhanced conformational sampling with stable binding with svPLA_2_ from Viperidae. These well-defined minima reflect that LTNF10 is conformationally flexible yet energetically optimized for interaction with the svPLA_2_ from Viperidae. In contrast, the mutant LTNF10-MT complexes (2QHD-LTNF10-MT and 4QEM-LTNF10-MT) display conformationally restricted low-energy regions, indicative of decreased conformational flexibility upon mutation in LTNF10. Therefore, the point mutation in LTNF10 (D5K) destabilizes key interactions with catalytic and calcium-binding residues. Thus, the mutation D5K shows opposite effects in LTNF10-svPLA_2_ binding in Viperidae.

**Figure 13.**
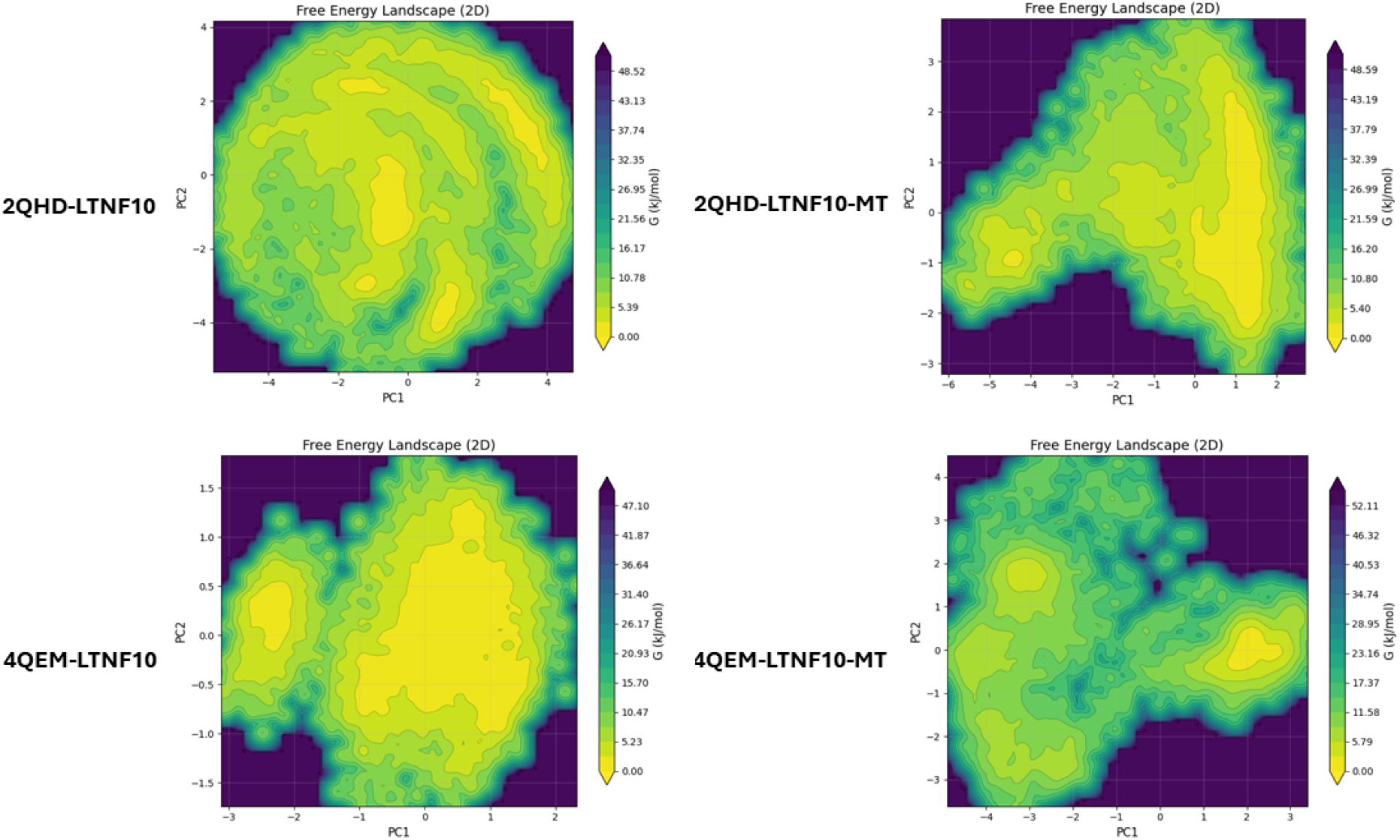
PCA (Principal component analysis) Derived Two Dimensional Free Energy Landscapes (ΔG, kJ/mol) of LTNF10 and LTNF10 MT Peptides from Viperidae. It shows conformational dynamics for LTNF10 and LTNF10 MT from Viperidae. The two dimensional free energy landscapes are plotted along the first two principal components (PC1 and PC2), with contour maps illustrating conformational basins and transitions observed during MD simulations. The color scale represents the relative free energy (ΔG, kJ/mol), ranging from low energy states (yellow) to high energy states (dark purple). The top row shows Free energy landscapes for the 2QHD with LTNF10 (left) and LTNF10 MT (right). The bottom row shows Free energy landscapes for the 4QEM with LTNF10 (left) and LTNF10 MT (right). Comparison between LTNF10 and LTNF10-MT peptides highlights shifts in conformational flexibility and energy minima, reflecting the impact of mutation in LTNF10–svPLA interactions from Viperidae.

These results suggest that LTNF10-MT-svPLA_2_s interactions in Elapidae benefit from Aspartic acid to Lysine substitution at the 5^th^ position of LTNF10 (D5K). This is supported by stronger electrostatic and hydrogen bond formation.

To understand the difference in binding pocket volume of svPLA_2_ bound with LTNF10 and LTNF10-MT, we calculated the mean pocket volume for each of LTNF10 and LTNF10-MT svPLA_2_ complexes (Table 6 and Figures S28–S29). We observed a significant reduction in binding pocket volume for the svPLA_2_ Group I (1U4J and 2WQ4: Elapidae) complexes (−49.7% and −58.1%, respectively), consistent with a more tightly packed, collapsed binding cavity relative to the LTNF10 bound svPLA_2_ complexes. We observed that Group IIA svPLA_2_ (4QEM: Viperidae) showed negligible reduction in binding pocket volume in LTNF10-MT bound 4QEM (−1.1%), while another Group IIB svPLA_2_ (2QHD: Viperidae) with Ser49 substitution at the catalytic site, showed a comparatively small reduction in LTNF10-MT bound form compared to Group I (−25.3%) (Figure 14, Table 6). These differences in pocket volume pertain to the trends observed MM/PBSA binding free energies (Table 4), supporting a structural basis for the contrasting effects of peptide flexibility on binding affinity between Group I and Group II svPLA2s.

**Figure 14.**
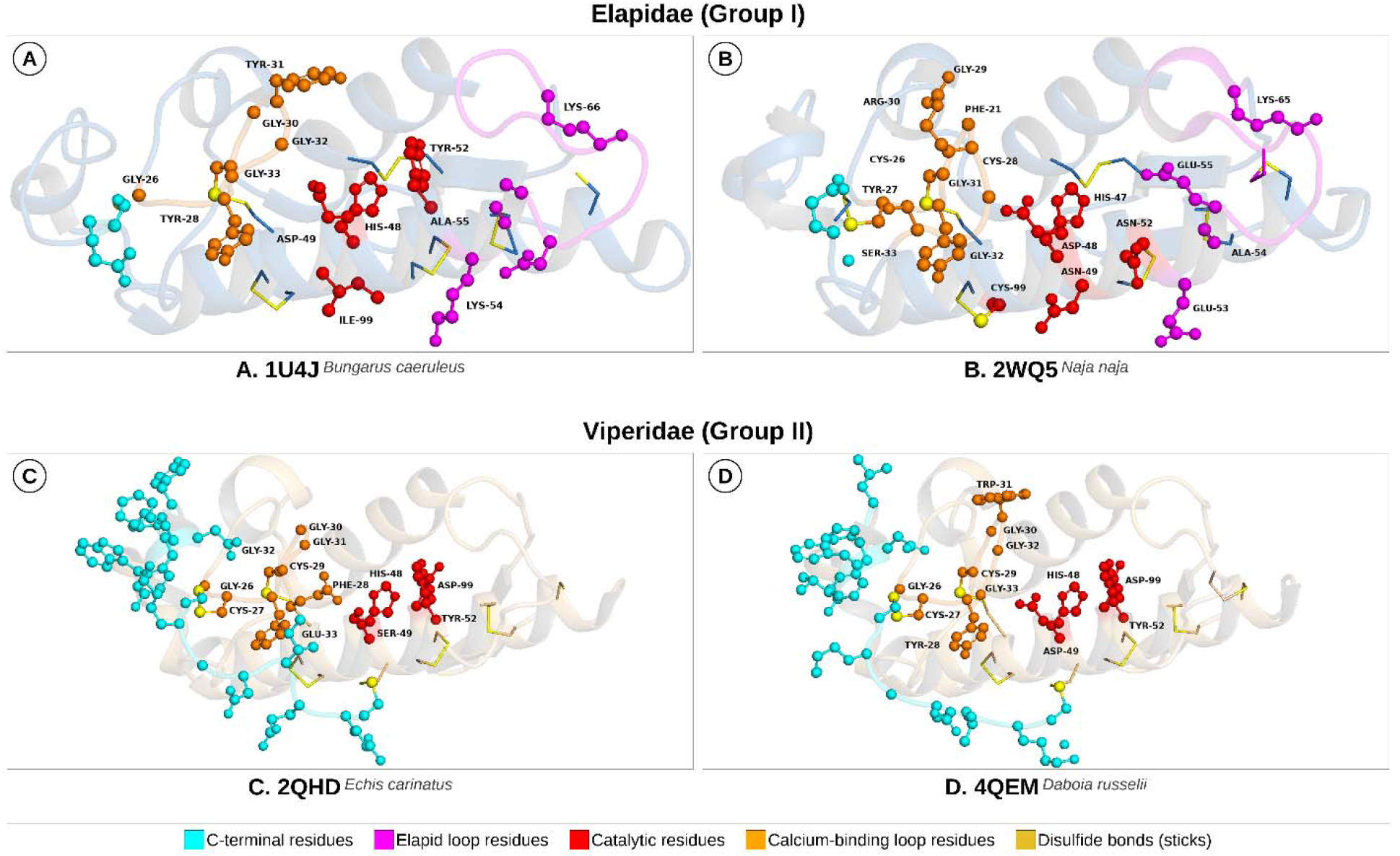
Structural features of the four svPLA2 targets: (A) 1U4J (Bungarus caeruleus) and (B) 2WQ5 (Naja naja), representing Group I (Asp49) Elapidae; (C) 2QHD (Echis carinatus) and (D) 4QEM (Daboia russelii), representing Group II Viperidae. Calcium-binding loop residues are shown in orange, catalytic residues in red, the elapid-loop region in magenta, and C-terminal residues in cyan; yellow-orange sticks denote disulfide bonds. The elapid-loop (magenta) is a defining feature of the Group I svPLA_2_. 1U4J (A) and 2WQ5 (B) have Ile99 and Cys99 substitutions instead of the standard Asp99, which forms the catalytic-dyad with His48, while 2QHD (C) have a Ser49 substitution in place of the canonical catalytic Asp49.

**Table 6.** Mean svPLA2 binding-pocket volume in the LTNF10- and LTNF10-MT-bound states.

| **Complex** | **Family** | **Mean pocket volume (Å³)** |  | **Δ Volume (%)** |
|---|---|---|---|---|
|  |  | **sv-PLA2-LTNF10** | **svPLA2-LTNF10-MT** |  |
| 1U4J (Indian krait) | Elapidae | 416.06 ± 91.06 | 209.43 ± 88.68 | –49.7% |
| 2WQ5 (Indian cobra) | Elapidae | 553.48 ± 194.89 | 231.94 ± 114.64 | –58.1% |
| 2QHD (Sindh saw-scaled viper) | Viperidae | 193.91 ± 79.88 | 144.79 ± 34.88 | –25.3% |
| 4QEM (Russell’s viper) | Viperidae | 308.13 ± 129.05 | 304.83 ± 104.71 | –1.1% |

Group I (Elapidae) svPLA_2_ have a conserved Asp49 catalytic residue, a calcium-binding loop and an elapid loop (distinct in Elapidae). An electronegative pocket can favourably interact with substituted Lys5 through electrostatic interactions with Asp49 in the catalytic region (1U4J and 2WQ5; Table S16-19). The Asp5 to Lys5 substitution also favoured improved interactions of Leu10 in LTNF10-MT with svPLA_2_ (Table S12-13).

Group II svPLA_2_ (Viperidae) possess more diverse structural features. Generally, Asp49 and its catalytic activity may be retained (Group IIA:4QEM) or it may be substituted to Ser49/Lys49 (Group IIB:2QHD), rendering it catalytically inactive. It has extended C-terminal residues for stability and lacks an additional disulfide bond formed with Elapid loop (i.e. absent in Viperidae), which gives it more structural diversity. It also has Asp99 that forms the catalytic dyad. In 2QHD (Viperidae), Lys5 in LTNF10-MT was observed to interact unfavourably with Ser49, but improved interactions were observed for Leu10 and Met4 in LTNF10-MT (Table S14). The pocket volume in LTNF10 bound 2QHD (Viperidae) was already compact, but the reduction in pocket volume in LTNF10-MT bound form suggests increased participation of other residues in molecular interactions (Leu10 and Met4; Table S18). Lys5 in the binding pocket tends to stay unfavourable, while it might also indirectly influence the increase in molecular interactions for Leu10 and Met4 (Table S18). On the other hand, in 4QEM (Viperidae), we again see favourable interactions and participation by Asp5 in LTNF10, but its substitution to Lys5 leads to unfavourable interactions, while Lys2, Ala3 and Met4 were observed to have a significant increase in favourable interactions (Table S15). Lys5 in proximity to two negatively charged residues (Asp49 and Asp99; Figure 14) might be the reason for its reduced participation. So, when the pocket volume doesn’t show any significant change, other LTNF10-MT residues Lys2, Ala3 and Met4 compensated for the lost interactions.

## Discussion

In two structurally and chemically different ligands, LTNF10 (peptide) and Varespladib (small-molecule) (24,83), The interactions observed in both cases show similar binding patterns with the hydrophobic channel, calcium-loop region, and catalytic site, which suggests inhibition potential of LTNF10 and LTNF10-MT peptides (Table S22). In the present study, we considered Varespladib as a reference inhibitor molecule to confirm the key residues in the binding region for further detailed analysis of LTNF10 binding. Therefore, comparative analysis of binding free energies for Varespladib and LTNF10 was not performed.

The most promising mutation in LTNF10 (Asp5 to Lys5) formed more stabilizing interactions. In svPLA_2_-LTNF10-MT complexes from Indian Cobra and Common Krait, which belong to Elapidae, where the mutated residue Lys5 forms new stable hydrogen bonds, leading to enhanced binding as confirmed by MD simulations and MM/PBSA analysis. On the other hand, there was a reduction in binding affinities upon mutation from Asp5 to Lys5 in Viperidae. This can also be observed from the loss of hydrogen bond interactions in Viperidae. The difference in binding profile of Elapidae and Viperidae svPLA complexes following mutation of the LTNF10 peptide can also be rationalized in light of their distinct structural and functional classifications. Elapidae svPLA s belong to the Group I class of Phospholipase A_2_, characterized by a conserved Asp49 catalytic residue and a well defined calcium binding loop that supports robust enzymatic activity and neurotoxic function. Within this framework, LTNF10-MT peptide shows enhanced conformational diversity and flexibility, enabling the peptide to adapt dynamically to the flexible active site environment in svPLA_2_ and maintain favourable binding energetics. In contrast, svPLA from Viperidae are members of the Group II class of Phospholipase A_2_, which exhibit greater structural heterogeneity, including variation at Ser49 that abolishes enzymatic activity but confers direct myotoxic effect through membrane disruption. In svPLA_2_ from Viperidae, the same mutation (D5K) leads to a reduction in binding affinity, which can be explained by their broader structural diversity and alternative modes of toxicity. This divergence underscores how lineage specific structural variation in the svPLA_2_ binding site. Our findings highlight the evolutionary distinction between Group I and Group II PLA s, where differences in catalytic site and functional roles shape the impact of LTNF10 mutation on binding affinity and conformational landscapes. The enhanced binding affinity of LTNF10-MT in svPLA_2_ from Elapidae suggests that the incorporated Asp5 to Lys5 mutation confers target-specific optimization and improved binding affinity.

The pocket-volume analysis offers a structural rationale for the divergent effect of Asp5 to Lys5 on binding affinity. In the Group I (Asp49) Elapidae complexes (1U4J and 2WQ5), the LTNF10-MT’s greater conformational flexibility (PCA, Figure 12) along with reduction in pocket volume, enables closer packing in the catalytic region and calcium-binding loop, which leads to more persistent electrostatic contacts with Lys5 (Tables S9, S16–S17), consistent with the improved MM/PBSA binding free energies for LTNF10-MT in these complexes and suggesting an interesting possibility of enthalpy-entropy compensation (Table 4). In the case of 2QHD, a Group IIB variant, it is mechanistically distinct, as the Ser49 substitution makes it catalytically inactive. It primarily acts through membrane disruption rather than hydrolysis. The pocket-volume reduction in 2QHD’s svPLA2-LTNF10-MT complex (25.3%) reflects this change in structural diversity. Replacing acidic Asp49 with neutral Ser49 shifts the pocket’s electrostatic microenvironment, partially offsetting the favourable electrostatic gain from Asp5 to Lys5 (via Glu33, Table S10), also corroborated by the reduced MM/PBSA binding free energies. While in 4QEM, a Group IIA variant, the catalytic and catalytic dyad residues Asp49 and Asp99, respectively, were retained; therefore, Lys5 from LTNF10-MT, in the vicinity of two negatively charged residues, reduced its participation in interactions while also promoting interactions with Lys2, Ala3 and Met4.

Therefore, these observations indicate that the differential effects of Asp5 to Lys5 mutation on binding affinity of LTNF10-MT with Group I and Group II svPLA_2_ were primarily driven by group-specific structural and dynamical differences in binding and active site regions, and resultant differences in LTNF10-MT flexibility upon binding to svPLA_2_ from these two groups.

## Conclusions

The comprehensive structure-guided mutation screening and optimization successfully enhanced the binding affinity of LTNF10 toward svPLA toxins from the Group I Elapidae family. The selected Asp5 to Lys5 mutation led to enhanced conformational flexibility, improved binding energetics, and structural complementarity, leading to the formation of stable complexes with Group I svPLA_2_. These findings highlight the importance of the active site charge microenvironment, binding affinity and conformational flexibility in computational design and engineering of peptide inhibitors against snake venom enzyme variants from diverse snake species. The LTNF10 and LTNF10-MT peptide candidates can be promising candidates for further in-vitro experimental validation and therapeutic translation against snake venom svPLA_2_ enzymes.

## Data availability

The data underlying this article are available in the article and in its online supplementary material.

## Declaration of generative AI and AI-assisted technologies for schematic conceptual illustrations

The author(s) used Claude Sonnet 5.5 to prepare Figure 8 for schematic representation of the results. After using this tool/service, the author(s) reviewed and edited the content as needed and take(s) full responsibility for the content of the published article.

## Supporting information

https://mahindraecolecentrale-my.sharepoint.com/:u:/g/personal/bipin_singh_mahindrauniversity_edu_in/IQAeX4TqPqSvRLKd9djg7UNRAavigwuoZ8EdePYakYQRqlc?e

