## Supplementary material for "Target Specific Conformational Flexibility and Electrostatic Complementarity Modulate LTNF10 Peptide Binding to Phospholipase A2 Variants from Snake Venoms": https://mahindraecolecentrale-my.sharepoint.com/:u:/g/personal/bipin_singh_mahindrauniversity_edu_in/IQAeX4TqPqSvRLKd9djg7UNRAavigwuoZ8EdePYakYQRqlc?e

**Table S1**. List of svPLA2 3D structures available in RCSB PDB Database.

| **S. No.** | **PDB ID** | **Year** | **Origin** | **Family** | **Species** | **Sequence Length** | **Resolution (Å)** |
|---|---|---|---|---|---|---|---|
| 1 | 2RD4 | 2007 | India | Elapidae | *Naja sagittifera* | 119 | 2.9 |
| 2 | 1MF4 | 2003 | India | Elapidae | *Naja sagittifera* | 119 | 1.9 |
| 3 | 1ZM6 | 2005 | India | Elapidae | *Naja sagittifera* | 119 | 2.6 |
| 4 | 1T37 | 2004 | India | Elapidae | *Naja sagittifera* | 119 | 2.6 |
| 5 | 1A3D | 1998 | India | Elapidae | *Naja naja* | 119 | 1.8 |
| 6 | 1A3F | 1998 | India | Elapidae | *Naja naja* | 119 | 2.65 |
| 7 | 2QHE | 2007 | India | Viperidae | *Echis carinatus* | 122 | 2 |

| | | | | | | | |
|---|---|---|---|---|---|---|---|
| 8 | 3BJW | 2007 | India | Viperidae | *Echis carinatus* | 122 | 2.3 |
| 9 | 2QHD | 2007 | India | Viperidae | *Echis carinatus* | 122 | 1.95 |
| 10 | 1OZ6 | 2003 | India | Viperidae | *Echis carinatus* | 120 | 2.6 |
| 11 | 2H4C | 2007 | India | Viperidae | *Daboia siamensis* | 122 | 2.6 |
| 12 | 1OXL | 2004 | India | Viperidae | *Daboia russelii* | 121 | 1.8 |
| 13 | 4QF7 | 2014 | India | Viperidae | *Daboia russelii* | 121 | 1.48 |
| 14 | 4QEM | 2014 | India | Viperidae | *Daboia russelii* | 121 | 1.2 |
| 15 | 4QF8 | 2014 | India | Viperidae | *Daboia russelii* | 121 | 1.65 |
| 16 | 5VET | 2018 | India | Viperidae | *Daboia russelii* | 121 | 2 |
| 17 | 2QVD | 2007 | India | Viperidae | *Daboia russelii* | 121 | 1.93 |
| 18 | 1OQS | 2003 | India | Viperidae | *Daboia siamensis* | 122 | 1.9 |

| | | | | | | | |
|---|---|---|---|---|---|---|---|
| 19 | 1KPM | 2002 | India | Viperidae | *Daboia russelii* | 121 | 1.8 |
| 20 | 1JQ9 | 2002 | India | Viperidae | *Daboia russelii* | 121 | 1.8 |
| 21 | 3H1X | 2009 | India | Viperidae | *Daboia russelii* | 121 | 1.4 |
| 22 | 2ARM | 2005 | India | Viperidae | *Daboia russelii* | 121 | 1.23 |
| 23 | 1JQ8 | 2002 | India | Viperidae | *Daboia russelii* | 121 | 2 |
| 24 | 1FE5 | 2001 | India | Elapidae | *Bungarus caeruleus* | 118 | 2.45 |
| 25 | 2OSN | 2007 | India | Elapidae | *Bungarus caeruleus* | 118 | 2.5 |
| 26 | 1DPY | 2000 | India | Elapidae | *Bungarus caeruleus* | 118 | 2.45 |
| 27 | 1G2X | 2003 | India | Elapidae | *Bungarus caeruleus* | 118 | 2.5 |
| 28 | 1U4J | 2004 | India | Elapidae | *Bungarus caeruleus* | 118 | 2.18 |
| 29 | 1TC8 | 2004 | India | Elapidae | *Bungarus caeruleus* | 118 | 2.7 |

| | | | | | | | |
|---|---|---|---|---|---|---|---|
| 30 | 2WQ5 | 2010 | India | Elapidae | *Naja naja* | 119 | 1.65 |
| 31 | 1Y75 | 2005 | India | Elapidae | *Naja sagittifera* | 118 | 2.3 |
| 32 | 1XXW | 2005 | India | Elapidae | *Naja sagittifera* | 119 | 2.7 |
| 33 | 1TD7 | 2004 | India | Elapidae | *Naja sagittifera* | 119 | 2.5 |
| 34 | 1S6B | 2004 | India | Elapidae | *Naja sagittifera* | 119 | 1.6 |
| 35 | 1YXH | 2005 | India | Elapidae | *Naja sagittifera* | 126 | 1.86 |

**Table S2.** List of peptides collected for early analysis of their properties and binding potential.

| **Peptide Name** | **Reported Target Protein** | **Peptide Sequence** |
|---|---|---|
| LTNF15 | SVMP | LKAMDPTPPLWIKTE |
| LTNF10 | SVMP | LKAMDPTPPL |
| Synthetic design | PLA2 | VIAL |
| Synthetic design | PLA2 | VAFRS |
| Synthetic design | PLA2 | LAIYS |
| Synthetic design | PLA2 | FLSYK |

| | | |
|---|---|---|
| Synthetic design | 3FTX | [PEG]6-YMWDGWYM-NH2e |
| Turmerin | Not specified | >sp\|P85278\|TURM_CURLO Turmerin (Fragment) OS=Curcuma longa OX=136217 PE=1 SV=2 LCPLDVLQLSSELLDIDGNEVEASRILSDITAF GGIRCPLTVVQSRGIGTIISSPYRFIA EGHPLSLKDMDGWFRVSDDEFNNYK |
| snakin-z | Not specified | >tr\|A0A6P3YX70\|A0A6P3YX70_ZIZJJ snakin-2-like OS=Ziziphus jujuba OX=326968 GN=LOC107406494 PE=3 SV=1 MMQKYDGNIEKSSLDGKIDCGGACAARCQ KSSRPRLCKRACGTCCQRCNCVPPGTAGQE VCPCYAALTTHGGRRKCP |
| snaking-z | Not specified | >tr\|A0A6P3ZSR4\|A0A6P3ZSR4_ZIZJJ snakin-2-like OS=Ziziphus jujuba OX=326968 GN=LOC107419922 PE=3 SV=1 MAIRLLLLMALLLICLAQVSPDSKIEDQETH VVVRGTNRRLFPFLDCGGLCKRRCSLHSR PNVCIRACGTCCIRCKCVPPGTAGNREMCGR CYTDMTTHGNRTKCP |
| MP-4 | PLA2 | >tr\|A0A158RFS0\|A0A158RFS0_MUCPR MP-4 OS=Mucuna pruriens OX=157652 PE=1 SV=1 KNDAEPVIDTDGNPLLHRGKYYIMPSNWGP PGGGLRLGKTRNLNCPVTVLQDYNEAINGL PVKFNIREILPRTIFTDTELNIEFTEKPNCAEN RAWSLFKGDDRGHKARVGIGGSNGHPG GEMLRGGFYGEQHGLRNGTYKLVFCRDGSS TCLDVGRYGNREGRRLGLSEAGELGVGFEK AGGGN |
| 95Mat5 | 3FTX | [W/Y]YxxGxY motif |
| JB006-free | PLA2 | H-DHWVWGWNYQYQPQEWHTES-OH |

**Elapidae**

1. *Bungarus caeruleus* (1U4J)

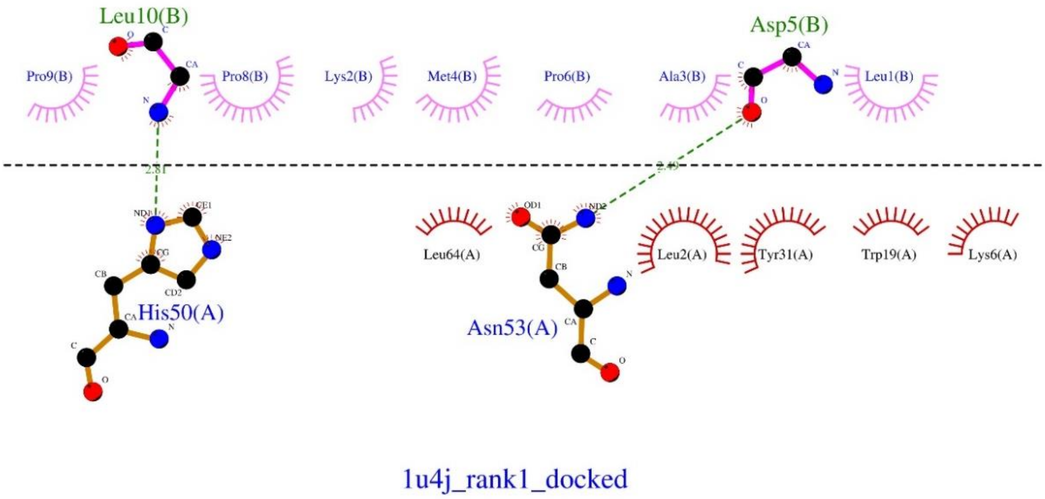


**Figure S1.** Interactions between PLA2 from *Bungarus caeruleus* with LTNF10.

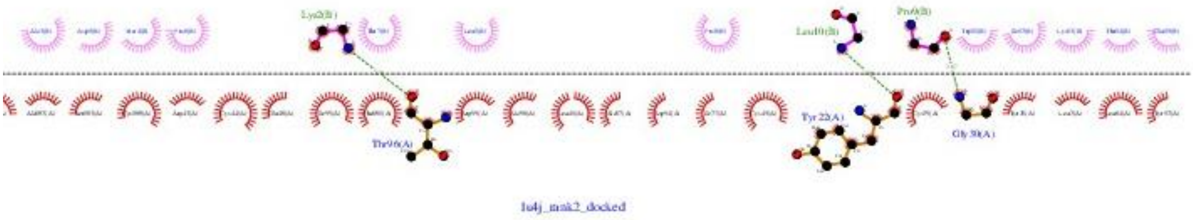

**Figure S2**. Interactions between PLA2 from *Bungarus caeruleus* with LTNF15.

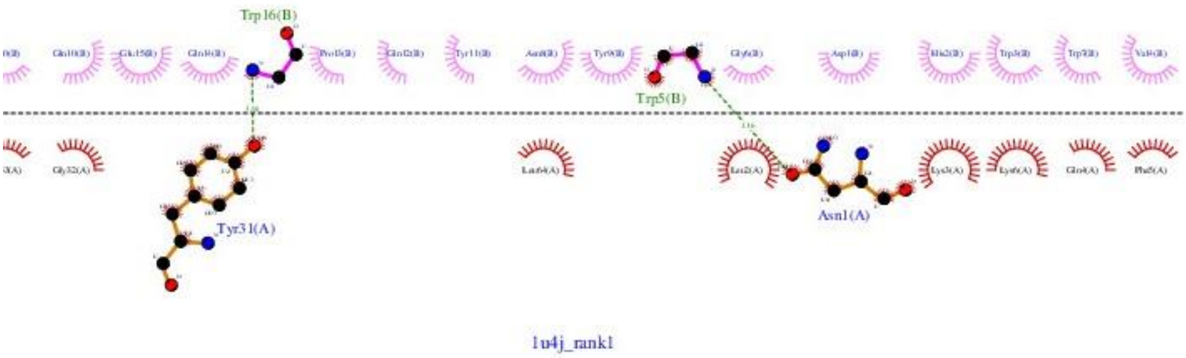


**Figure S3.** Interactions between PLA2 from *Bungarus caeruleus* with MP-4.

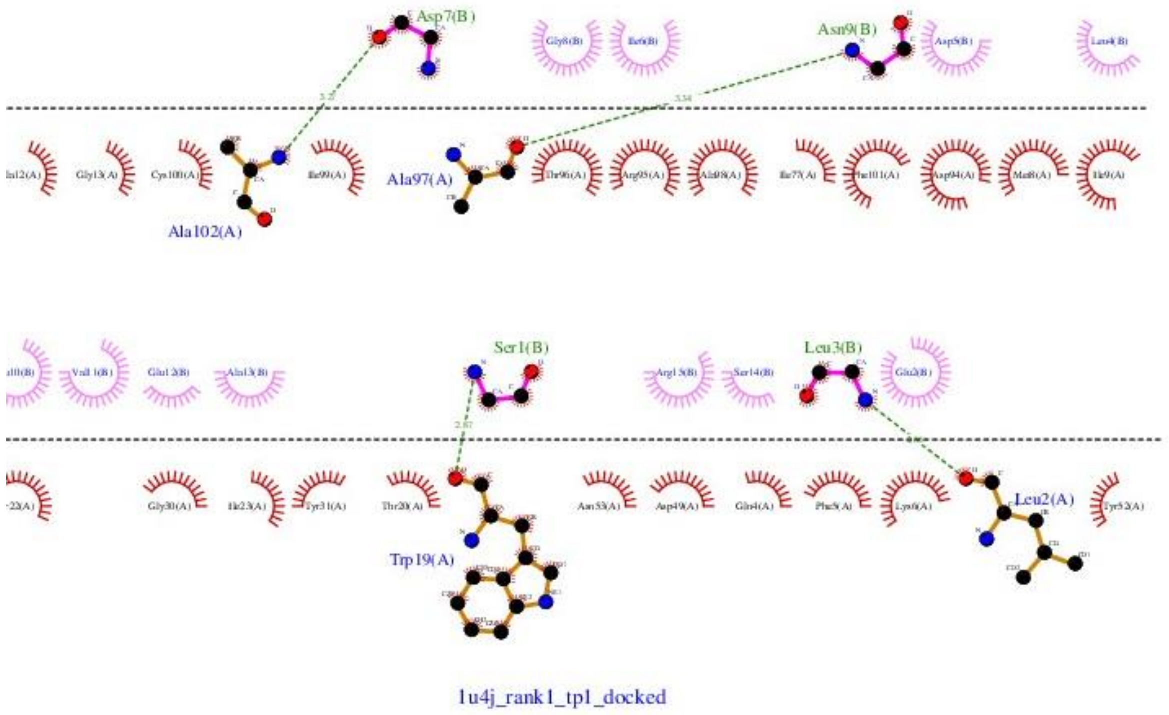


**Figure S4**. Interactions between PLA2 from *Bungarus caeruleus* with Turmerin derived peptide candidate 1 (TP1).

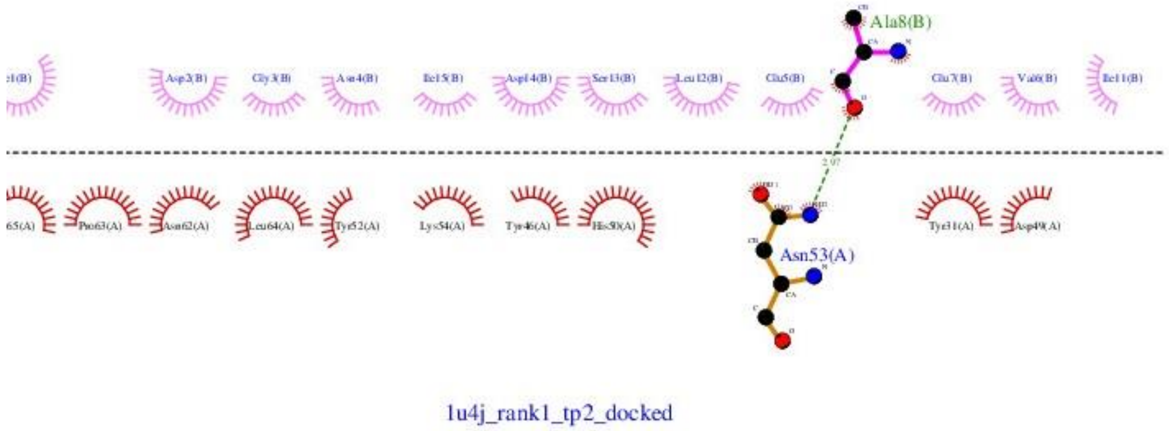


**Figure S5.** Interactions between PLA2 from *Bungarus caeruleus* with Turmerin derived peptide candidate 2 (TP2).

2. *Naja naja* (2WQ5)

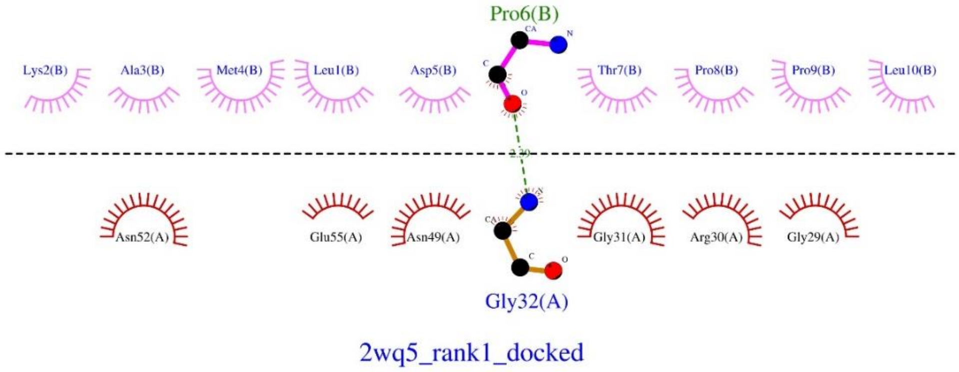


**Figure S6.** Interactions between PLA2 from *Naja naja* with LTNF10.

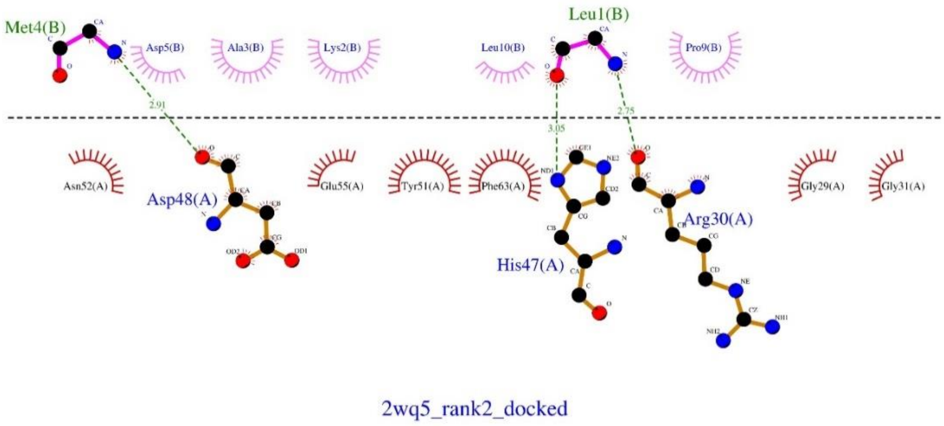


**Figure S7.** Interactions between PLA2 from *Naja naja* with LTNF15.

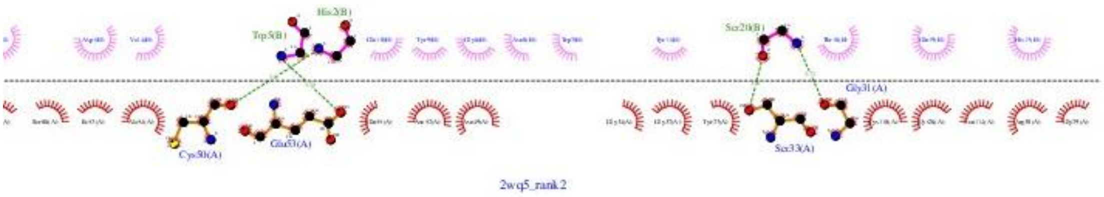


**Figure S8.** Interactions between PLA2 from *Naja naja* with MP-4.

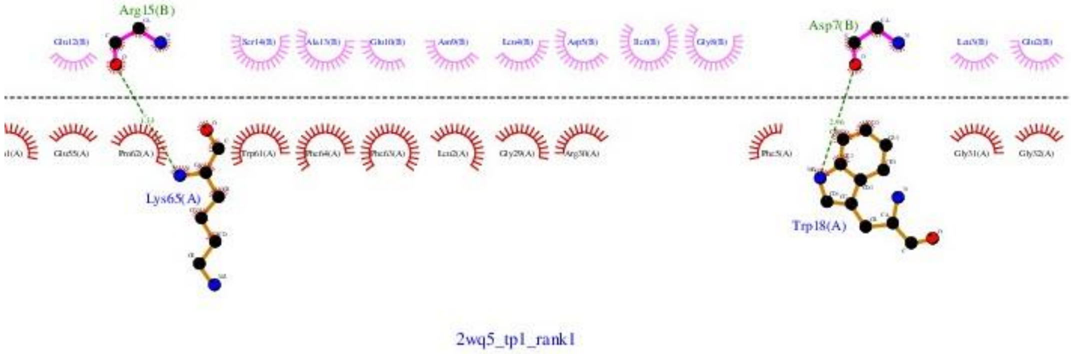


**Figure S9.** Interactions between PLA2 from *Naja naja* with Turmerin derived peptide candidate 1 (TP1).

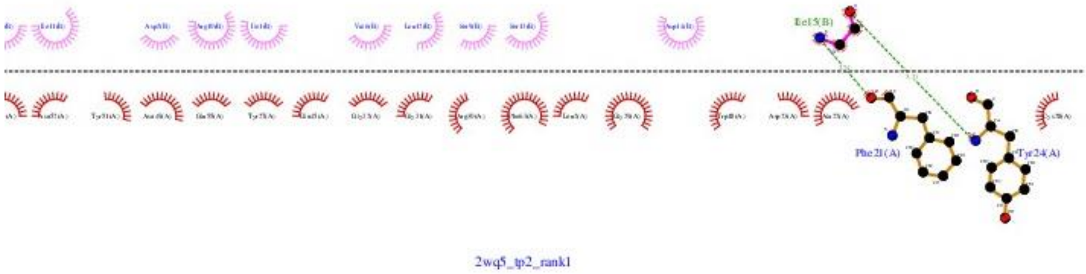


**Figure S10.** Interactions between PLA2 from *Naja naja* with Turmerin derived peptide candidate 2 (TP2).

**Viperidae**

1. *Daboia russelii pulchella* (4QEM)

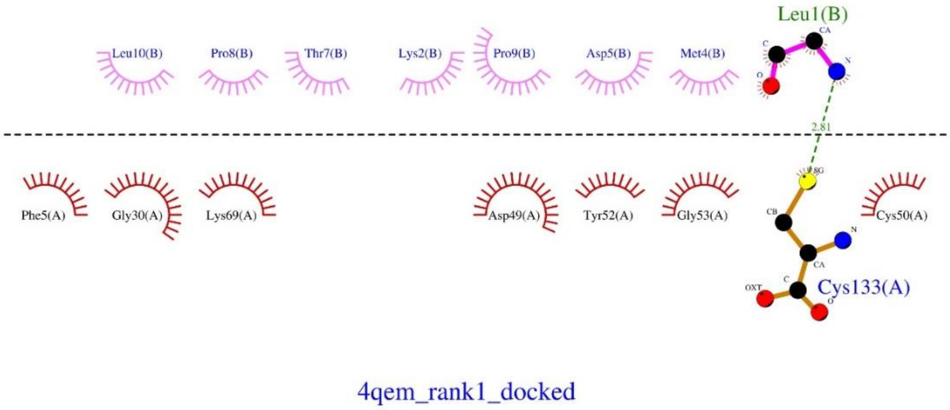


**Figure S11.** Interactions between PLA2 from *Daboia russelii pulchella* with LTNF10.

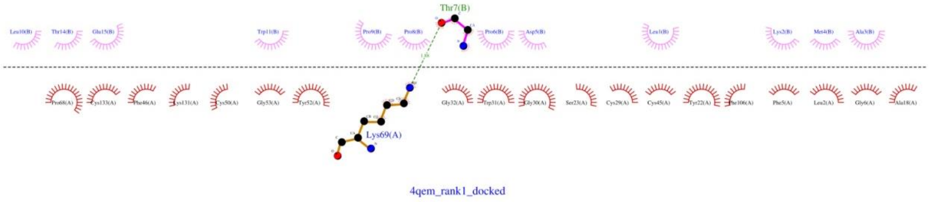


**Figure S12.** Interactions between PLA2 from *Daboia russelii pulchella* with LTNF15.

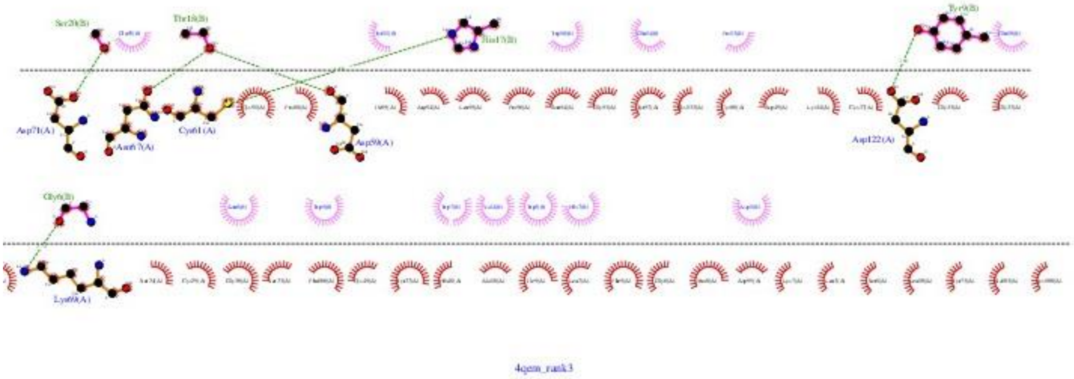


**Figure S13.** Interactions between PLA2 from *Daboia russelii pulchella* with MP-4.

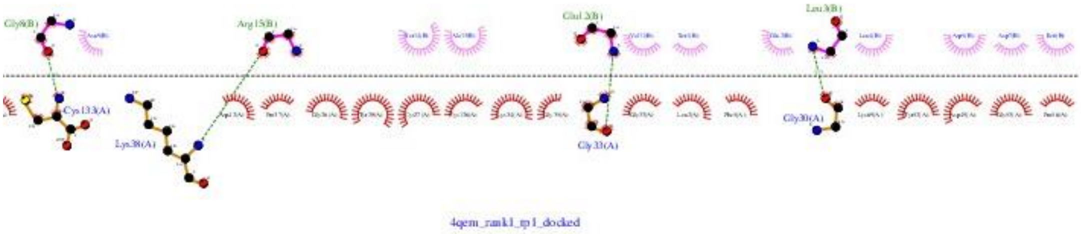


**Figure S14**. Interactions between PLA2 from *Daboia russelii pulchella* with Turmerin derived peptide candidate 1 (TP1).

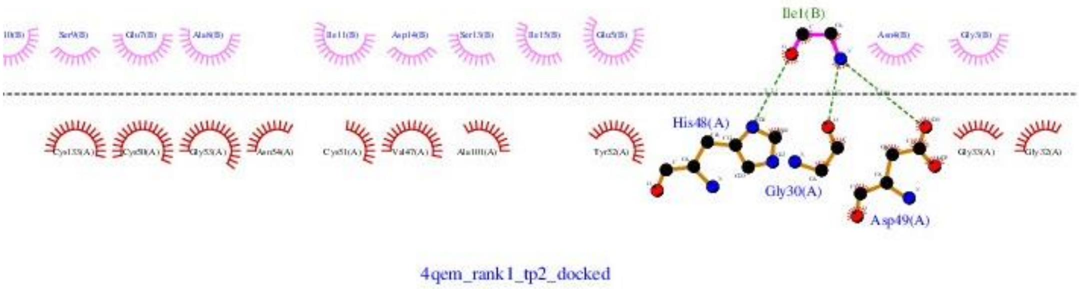

**Figure S15.** Interactions between PLA2 from *Daboia russelii pulchella* with Turmerin derived peptide candidate 2 (TP2).

2. *Echis carinatus* (2QHD)

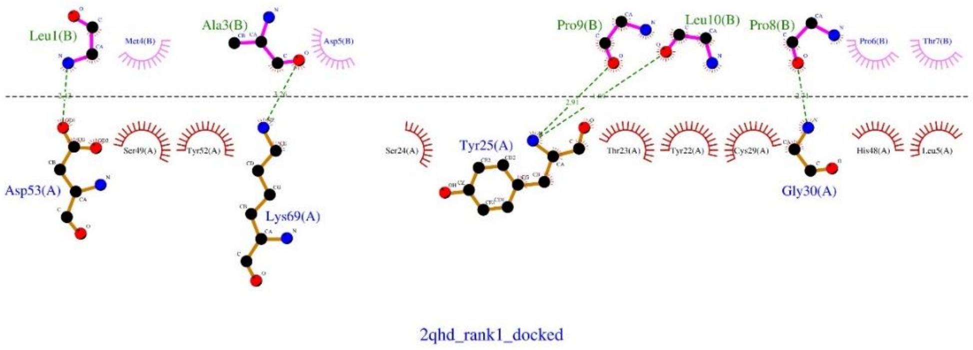


**Figure S16.** Interactions between PLA2 from *Echis carinatus* with LTNF10.

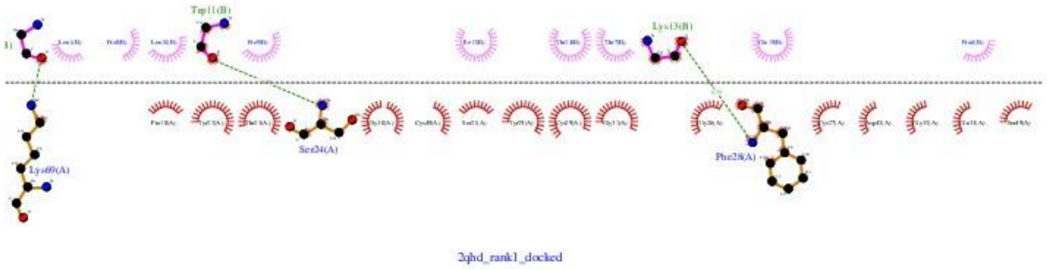


**Figure S17.** Interactions between PLA2 from *Echis carinatus* with LTNF15.

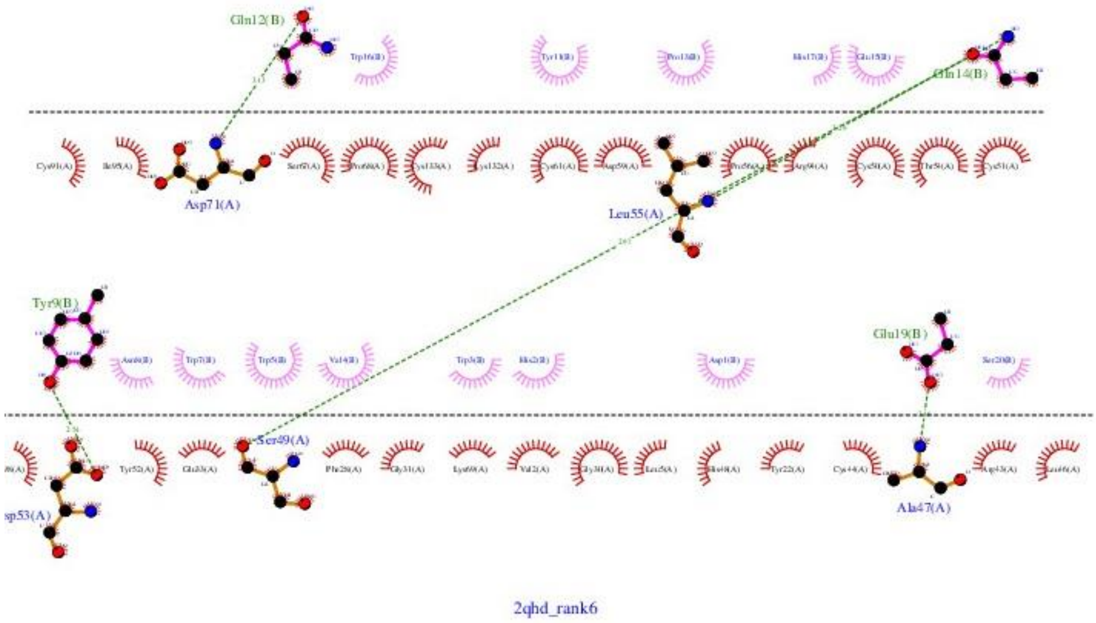


**Figure S18.** Interactions between PLA2 from *Echis carinatus* with MP-4.

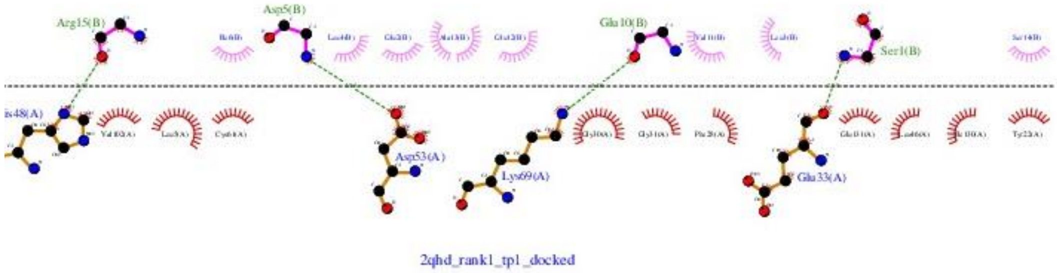


**Figure S19.** Interactions between PLA2 from *Echis carinatus* with Turmerin derived peptide candidate 1 (TP1).

**Table S3.** Positional scanning results from FoldX: predicted ΔΔG values (kcal/mol) for single-point mutations at selected residues in LTNF10 complex with svPLA₂ structures (PDB IDs 1U4J, 2WQ5, 2QHD, 4QEM).

| MUTATION | Change in binding affinity upon mutation(kcal/mol) | | | |
|---|---|---|---|---|
| | 1U4J | 2WQ5 | 2QHD | 4QEM |
| METB4M | 0 | 0 | 0 | 0 |
| METB4G | 2.25649 | -1.54067 | 1.08212 | -0.38213 |
| METB4A | 1.25728 | -1.18014 | 0.510642 | 0.969499 |
| METB4L | -0.41501 | 0.212318 | 0.415309 | 0.31286 |
| METB4V | 1.38325 | 0.596679 | 2.01443 | 3.64246 |
| METB4I | 0.803097 | 0.632345 | 3.10965 | 4.00555 |
| METB4P | 3.67952 | -1.83891 | 3.89205 | 5.13577 |
| METB4R | 0.417846 | -0.2544 | -0.28713 | 0.770786 |
| METB4T | 2.37314 | 0.169064 | 2.3121 | 4.38346 |
| METB4S | 1.98022 | -0.75754 | 1.63751 | 1.12334 |
| METB4C | 1.17892 | -0.08664 | 0.605988 | 0.974095 |
| METB4M | -3.27E-13 | -2.84E-14 | 0.012978 | 0 |
| METB4K | 0.295168 | -0.30052 | 0.578719 | 0.561961 |
| METB4E | 1.98187 | -0.31784 | 1.89855 | 1.01772 |
| METB4Q | 1.37663 | -0.52532 | 0.98642 | 0.727466 |
| METB4D | 2.20448 | 0.364927 | 2.12597 | 1.49969 |
| METB4N | 1.36141 | -0.44227 | 1.58458 | 1.54049 |
| METB4W | 0.834574 | 0.208386 | 0.60585 | 1.85339 |
| METB4Y | 0.668636 | 0.280871 | -0.19563 | 0.865686 |
| METB4F | 0.31895 | 0.290511 | -0.29275 | 0.672531 |
| METB4o | 1.59544 | 0.388939 | 0.960608 | 2.26022 |
| METB4y | 0.648286 | 0.31505 | 0.705014 | 0.330887 |
| METB4p | 4.85151 | 1.72757 | 6.42472 | 6.08586 |
| METB4s | 2.91934 | 0.753069 | 3.93402 | 1.87129 |
| METB4h | 3.83696 | -2.34349 | 3.70732 | 13.9466 |
| ASPB5D | 0 | 0 | 0 | 0 |
| ASPB5G | -0.27558 | -1.04454 | -4.34889 | -7.92883 |
| ASPB5A | -0.33328 | -0.60276 | -3.4308 | -7.71884 |
| ASPB5L | -0.3879 | -1.84596 | -1.59214 | -2.87335 |
| ASPB5V | 1.42215 | -1.18944 | -3.08171 | -5.45163 |
| ASPB5I | 1.50701 | -1.5116 | -2.76447 | -0.16414 |
| ASPB5P | 1.52839 | -0.33826 | -4.23863 | 0.339785 |
| ASPB5R | -0.37663 | -1.39079 | -2.47985 | 3.3445 |
| ASPB5T | 1.11939 | -0.82656 | -2.6259 | -5.91672 |
| ASPB5S | -0.80555 | -1.00673 | -3.3094 | -6.25237 |
| ASPB5C | -0.67733 | -0.91337 | -3.47501 | -7.34121 |
| ASPB5M | -0.508 | -1.86279 | -2.53136 | -1.81301 |
| ASPB5K | -0.36087 | -1.58482 | -2.09601 | -2.60031 |
| ASPB5E | 1.26294 | -0.97732 | -0.09325 | 3.16422 |

| | | | | |
|---|---|---|---|---|
| ASPB5Q | 0.095342 | -1.14401 | -0.52964 | 0.418325 |
| ASPB5D | -0.64344 | -1.42E-14 | 2.84E-14 | 2.74216 |
| ASPB5N | -0.26011 | -0.4654 | -2.23383 | -2.09041 |
| ASPB5W | -0.15644 | -2.06011 | 1.95164 | 39.8194 |
| ASPB5Y | -0.55522 | -1.92127 | -0.84952 | 36.2547 |
| ASPB5F | -0.61793 | -2.00512 | -1.00502 | 20.9814 |
| ASPB5e | -0.66363 | 0.215093 | -2.12232 | 25.2898 |
| ASPB5y | -0.53034 | -2.93597 | -1.69955 | 38.5604 |
| ASPB5p | 4.78554 | -0.64636 | 6.43706 | 20.0825 |
| ASPB5s | 1.88946 | -0.53428 | 2.34223 | 9.2626 |
| ASPB5h | 8.34912 | 2.22867 | -2.89159 | 5.50877 |
| THRB7T | 0 | 0 | 0 | 0 |
| THRB7G | -3.44739 | -0.77481 | -2.97511 | -1.28739 |
| THRB7A | -3.51588 | -0.13596 | -3.02358 | -1.95767 |
| THRB7L | -2.41101 | -1.01519 | -1.94029 | 2.12439 |
| THRB7V | -1.39011 | -0.63454 | -1.50953 | -0.77486 |
| THRB7I | -3.19098 | -1.0895 | -1.30065 | 1.46239 |
| THRB7P | 1.41451 | 1.45009 | -5.87667 | -0.98353 |
| THRB7R | -3.56052 | -0.23954 | -1.15657 | 1.22483 |
| THRB7T | -1.36442 | 0 | 0 | 0.005725 |
| THRB7S | -2.53933 | 0.23011 | -1.09032 | -0.97371 |
| THRB7C | -4.75111 | 0.225191 | -2.72227 | -1.81536 |
| THRB7M | -3.4958 | -0.54213 | -3.39042 | -1.52148 |
| THRB7K | -0.63704 | -0.84872 | -1.02691 | 0.474501 |
| THRB7E | -0.97852 | -0.40723 | -0.39256 | 0.983786 |
| THRB7Q | -0.38946 | -0.85146 | -0.85659 | 1.08761 |
| THRB7D | -0.21304 | -0.20309 | 0.5865 | 0.409055 |
| THRB7N | -1.03386 | -0.46477 | -1.70899 | 0.236331 |
| THRB7W | 2.03427 | -0.18831 | 8.59577 | 11.7601 |
| THRB7Y | 2.29984 | -0.86702 | 12.3581 | 13.7108 |
| THRB7F | 2.35221 | -0.90349 | 7.89727 | 10.6673 |
| THRB7H | 1.74434 | -0.46514 | 5.53237 | 49.3385 |
| THRB7y | 6.80529 | -0.82311 | 12.9178 | 15.1618 |
| THRB7p | 3.45383 | 1.11815 | 7.63758 | 7.71887 |
| THRB7s | -0.65533 | 0.280064 | 0.872155 | 1.82564 |
| THRB7h | 1.22725 | 5.04981 | -4.79505 | 0.401185 |

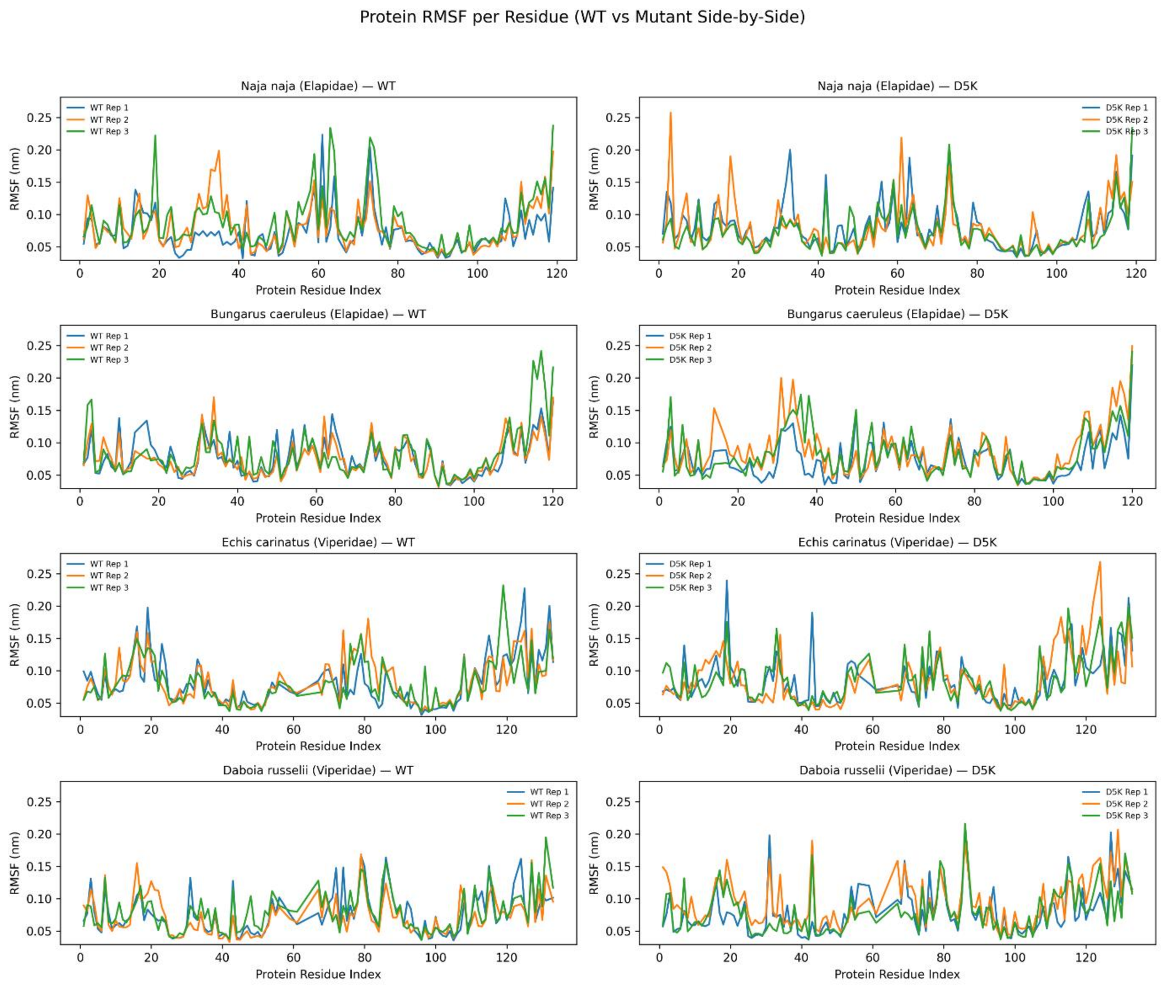


**Figure S20.** Residue-Wise RMSF Profiles of svPLA₂ from Elapidae and Viperidae in Complexes with LTNF10 and LTNF10-MT

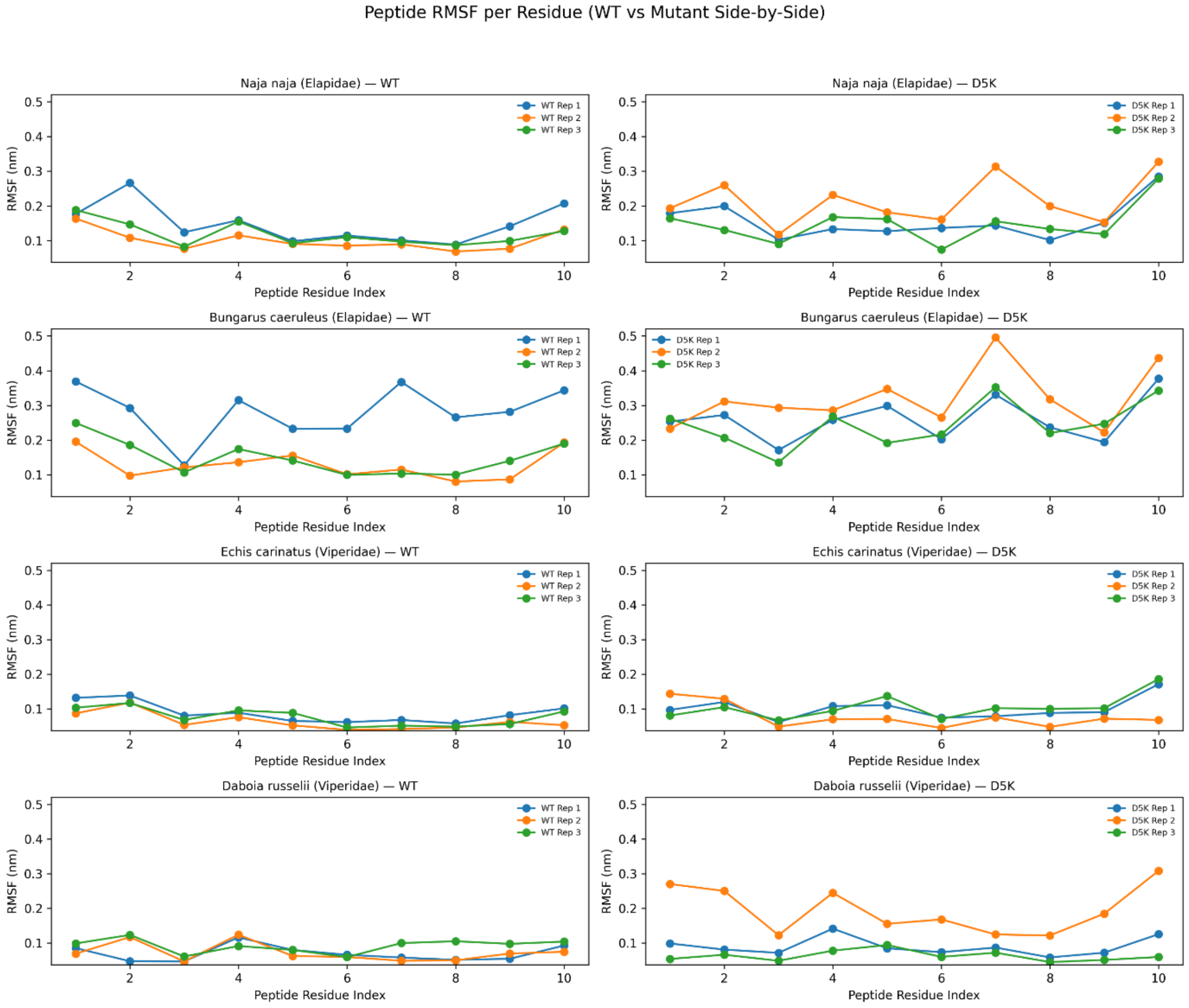


**Figure S21.** Residue-Wise RMSF Profiles of LTNF10 and LTNF10-MT Peptides in Complexes with svPLA$_2$ from Elapidae and Viperidae.

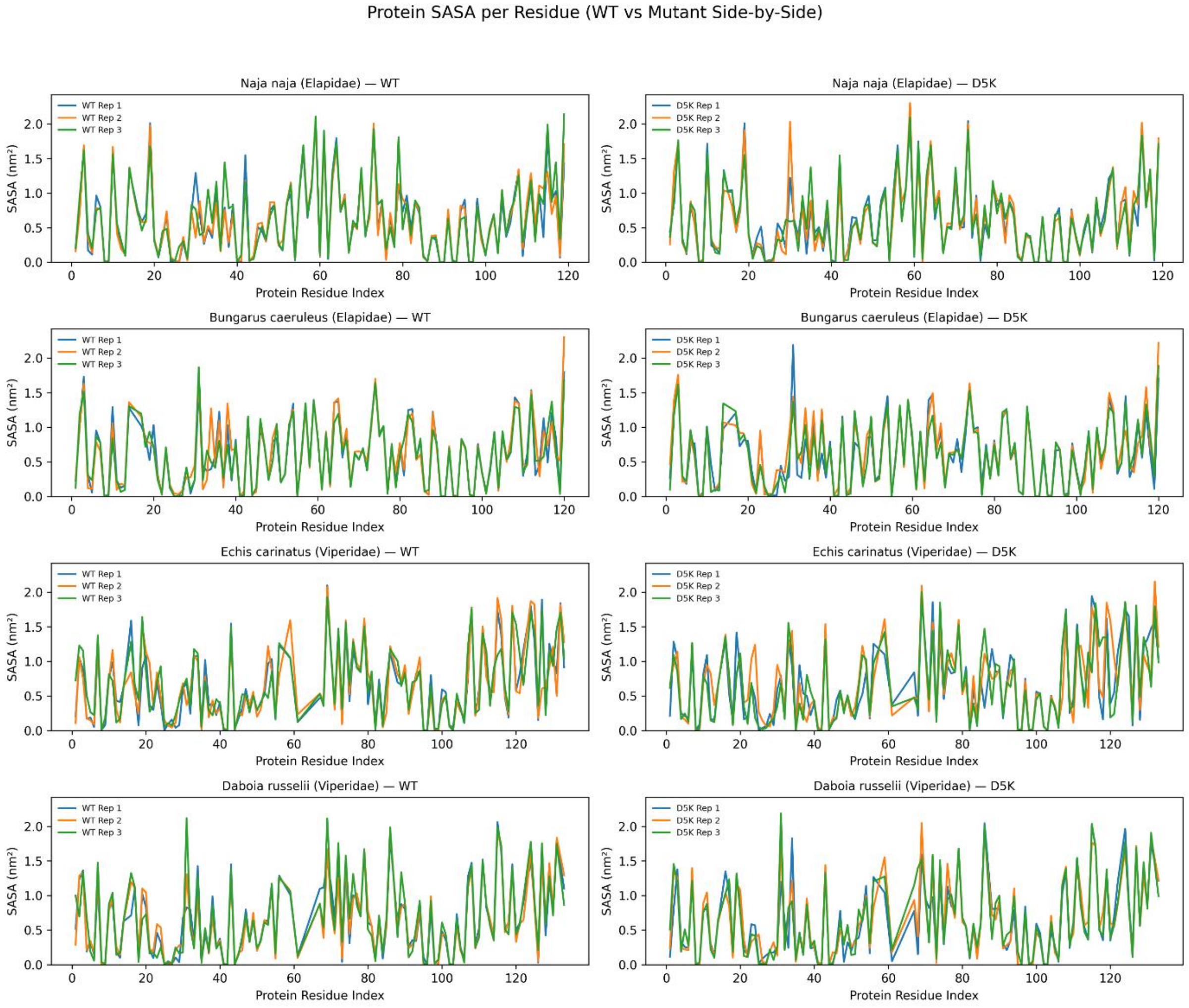


**Figure S22.** Residue-wise Solvent Accessible Surface Area (SASA) Profiles of svPLA2 from Elapidae and Viperidae when binding with LTNF10 and LTNF10-MT.

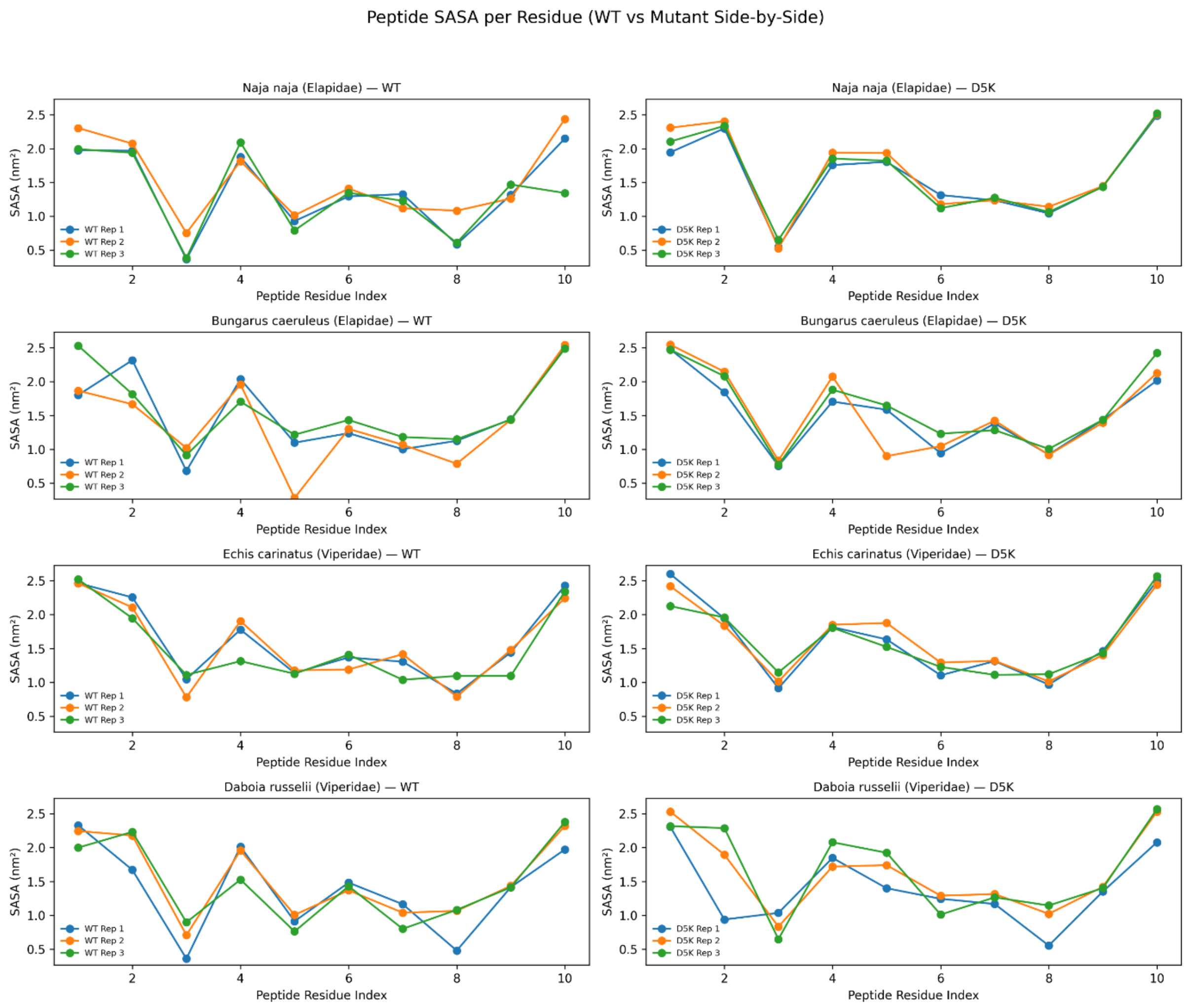


**Figure S23.** Residue-wise Solvent Accessible Surface Area (SASA) LTNF10 and LTNF10-MT peptides when binding with svPLA2.

**Table S4.** Average RMSF (nm) and SASA ($nm^2$) values in svPLA$_2$ 1U4J (Elapidae).

| | **RMSF (avg. in nm)** | | **SASA (avg in nm$^2$)** | |
|---|---|---|---|---|
| **Residues in svPLA** | **svPLA-LTNF10** | **svPLA-LTNF10-MT** | **svPLA-LTNF10** | **svPLA-LTNF10-MT** |
| LEU2 | 0.112233 | 0.0883 | 1.067 | 1.274667 |
| ILE9 | 0.055933 | 0.053767 | 0.016667 | 0.028667 |
| THR18 | 0.0813 | 0.070033 | 0.798 | 0.818667 |
| SER19 | 0.0798 | 0.071867 | 0.745 | 0.872667 |
| TYR20 | 0.071967 | 0.0732 | 0.821 | 0.728333 |
| ILE21 | 0.066833 | 0.0616 | 0.279333 | 0.252667 |
| CYS27 | 0.054767 | 0.069833 | 0.06 | 0.190667 |
| GLY28 | 0.056 | 0.083767 | 0.371333 | 0.372 |
| TYR29 | 0.054633 | 0.0642 | 0.136667 | 0.21 |
| GLY30 | 0.082 | 0.091167 | 0.306 | 0.554 |
| GLY31 | 0.137467 | 0.146 | 1.695 | 1.665667 |

| | | | | |
|---|---|---|---|---|
| SER32 | 0.095533 | 0.122833 | 0.258333 | 0.496333 |
| GLY33 | 0.089667 | 0.136067 | 0.422667 | 0.464 |
| HIS46 | 0.066233 | 0.0879 | 1.044333 | 1.034667 |
| ASP47 | 0.050667 | 0.052367 | 0.730333 | 0.731667 |
| TYR50 | 0.095 | 0.11915 | 0.979333 | 1.011 |
| LEU62 | 0.1109 | 0.1041 | 0.921 | 0.925333 |
| ILE63 | 0.070633 | 0.071033 | 0.178667 | 0.157333 |
| LYS64 | 0.112467 | 0.102933 | 1.259333 | 1.162 |
| THR65 | 0.096967 | 0.081367 | 1.327333 | 1.378 |
| TYR66 | 0.075767 | 0.067567 | 0.63 | 0.644 |
| SER67 | 0.067233 | 0.0808 | 0.907 | 0.921 |
| TYR68 | 0.046767 | 0.046667 | 0.151 | 0.147667 |
| THR69 | 0.067433 | 0.058867 | 0.628333 | 0.620333 |
| CYS70 | 0.060133 | 0.0615 | 0.572667 | 0.575667 |
| ALA113 | 0.074133 | 0.0769 | 0.575 | 0.387667 |
| SER114 | 0.1015 | 0.108567 | 0.419333 | 0.474 |
| THR115 | 0.157233 | 0.149533 | 0.882333 | 0.979667 |

**Table S5.** Average RMSF (nm) and SASA ($nm^2$) values in 2WQ5 (Elapidae).

| | RMSF (avg. in nm) | | SASA (avg in $nm^2$) | |
|---|---|---|---|---|
| **Residues in svPLA** | **svPLA-LTNF10** | **svPLA-LTNF10-MT** | **svPLA-LTNF10** | **svPLA-LTNF10-MT** |
| LEU2 | 0.101933 | 0.102933 | 0.849333333 | 1.023667 |
| ILE9 | 0.061133 | 0.062233 | 0.018 | 0.029667 |
| TRP18 | 0.096667 | 0.121133 | 0.635333333 | 0.797667 |
| TRP19 | 0.148467 | 0.101967 | 1.886333333 | 1.822667 |
| ASP20 | 0.061467 | 0.07 | 0.329333333 | 0.399 |
| PHE21 | 0.054933 | 0.056267 | 0.083666667 | 0.081333 |
| TYR27 | 0.064433 | 0.0624 | 0.32 | 0.446 |
| CYS28 | 0.0568 | 0.052667 | 0.037333333 | 0.307667 |
| GLY29 | 0.091033 | 0.0777 | 0.765666667 | 0.307 |
| ARG30 | 0.103567 | 0.0894 | 0.805 | 1.28 |
| GLY31 | 0.094567 | 0.0976 | 0.652333333 | 0.583 |
| GLY32 | 0.093467 | 0.1029 | 0.329666667 | 0.293333 |
| SER33 | 0.123767 | 0.127967 | 0.689 | 0.863667 |
| HIS47 | 0.046533 | 0.047733 | 0.309666667 | 0.293 |
| ASP48 | 0.0595 | 0.077033 | 0.766666667 | 0.768333 |
| TYR52 | 0.090467 | 0.072967 | 0.845666667 | 0.842 |
| PHE63 | 0.141067 | 0.127833 | 1.231666667 | 1.163 |
| PHE64 | 0.153967 | 0.1193 | 1.73 | 1.707 |
| LYS65 | 0.070867 | 0.074933 | 0.737 | 0.712667 |
| THR66 | 0.060967 | 0.0656 | 0.949666667 | 0.977333 |
| TYR67 | 0.048067 | 0.0484 | 0.147666667 | 0.132333 |
| SER68 | 0.062867 | 0.0692 | 0.556333333 | 0.527 |

| | | | | |
|---|---|---|---|---|
| TYR69 | 0.059733 | 0.063133 | 0.512333333 | 0.514667 |
| LYS70 | 0.1045 | 0.098767 | 1.291666667 | 1.256 |
| LEU113 | 0.1086 | 0.1125 | 1.024 | 0.894 |
| LYS114 | 0.100767 | 0.110133 | 0.755333333 | 0.772 |

**Table S6.** Average RMSF (nm) and SASA (nm$^2$) values in 2QHD (Viperidae).

| | RMSF avg in nm | | SASA avg in nm$^2$ | |
|---|---|---|---|---|
| Residues in svPLA | svPLA-LTNF10 | svPLA-LTNF10-MT | svPLA-LTNF10 | svPLA-LTNF10-MT |
| VAL2 | 0.075833 | 0.086533 | 1.114 | 1.108667 |
| ILE9 | 0.065433 | 0.068667 | 0.151 | 0.041333 |
| PRO18 | 0.100367 | 0.102967 | 0.254333 | 0.096667 |
| PHE19 | 0.163467 | 0.175067 | 1.355333 | 1.011 |
| PRO20 | 0.122033 | 0.092067 | 1.025667 | 1.001667 |
| SER21 | 0.108067 | 0.088 | 0.536667 | 0.303333 |
| TYR19 | 0.054433 | 0.0527 | 0.092333 | 0.057 |
| PHE28 | 0.0672 | 0.061333 | 0.125 | 0.140667 |
| CYS29 | 0.0556 | 0.057933 | 0.326333 | 0.179667 |
| GLY30 | 0.0699 | 0.086667 | 0.639 | 0.631667 |
| GLY31 | 0.079067 | 0.088833 | 0.603667 | 0.746333 |
| GLY32 | 0.0694 | 0.0777 | 0.381333 | 0.225 |
| GLU33 | 0.107133 | 0.127367 | 1.113 | 1.356333 |
| HIS47 | 0.047133 | 0.052833 | 0.544 | 0.57 |
| SER48 | 0.043833 | 0.048 | 0.317 | 0.304667 |
| TYR52 | 0.049467 | 0.059633 | 0.562 | 0.566667 |
| SER67 | 0.077367 | 0.0749 | 0.512 | 0.599667 |
| PRO68 | 0.074467 | 0.066067 | 0.405667 | 0.324667 |
| LYS69 | 0.098567 | 0.1113 | 2.034333 | 2.000667 |
| THR70 | 0.097833 | 0.093233 | 1.142 | 1.181 |
| ASN113 | 0.067033 | 0.105967 | 0.494667 | 0.572667 |
| LYS114 | 0.099967 | 0.101067 | 0.676 | 0.656333 |
| LYS115 | 0.130567 | 0.1735 | 1.580333 | 1.521 |

**Table S7.** Average RMSF (nm) and SASA (nm$^2$) values in 4QEM (Elapidae).

| | RMSF Avg in nm | | SASA Avg in nm2 | |
|---|---|---|---|---|
| Residues in svPLA | svPLA-LTNF10 | svPLA-LTNF10-MT | svPLA-LTNF10 | svPLA-LTNF10-MT |
| LEU2 | 0.082667 | 0.108167 | 1.072 | 1.204 |
| ILE9 | 0.054933 | 0.072833 | 0.018 | 0.018 |
| ALA18 | 0.082133 | 0.089833 | 0.089333 | 0.146 |
| ILE19 | 0.0956 | 0.122933 | 0.928667 | 1.240333 |
| PRO20 | 0.099933 | 0.105567 | 0.870667 | 0.916667 |
| SER21 | 0.085167 | 0.088967 | 0.202667 | 0.23 |

| | | | | |
|---|---|---|---|---|
| TYR27 | 0.039867 | 0.052333 | 0.008667 | 0.057333 |
| TYR28 | 0.042 | 0.050867 | 0.188333 | 0.199333 |
| CYS29 | 0.042333 | 0.0497 | 0.161667 | 0.189333 |
| GLY30 | 0.056967 | 0.0604 | 0.374 | 0.220667 |
| TRP31 | 0.0947 | 0.140267 | 1.419 | 1.733333 |
| GLY32 | 0.068567 | 0.069067 | 0.64 | 0.351333 |
| GLY33 | 0.0622 | 0.064533 | 0.286333 | 0.602333 |
| HIS47 | 0.062233 | 0.063367 | 0.726667 | 0.641667 |
| ASP48 | 0.063667 | 0.050867 | 0.335 | 0.235667 |
| TYR52 | 0.060367 | 0.062233 | 0.633 | 0.583 |
| ASN67 | 0.1067 | 0.116533 | 0.955 | 0.955333 |
| PRO68 | 0.072467 | 0.088067 | 0.681333 | 0.653333 |
| LYS69 | 0.0986 | 0.131067 | 1.756 | 1.722333 |
| SER70 | 0.088333 | 0.099133 | 0.982333 | 0.982667 |
| SER113 | 0.077033 | 0.092033 | 0.609 | 0.528333 |
| LYS114 | 0.070733 | 0.081867 | 0.396333 | 0.450667 |
| LYS115 | 0.134467 | 0.149133 | 1.995333 | 1.938667 |

**Table S8.** Decomposition energy contributions of significant residues in svPLA$_2$ from 1U4J (Elapidae) during complex formation with LTNF10 and LTNF10-MT, calculated using MM/PBSA (values in kJ/mol).

| svPLA$_2$ residue | svPLA$_2$-LTNF10 | svPLA$_2$-LTNF10-MT |
|---|---|---|
| LEU-2 | -2.25767 | 0.375667 |
| ILE-9 | 1.526333 | 2.315667 |
| THR-20 | 0.141 | -1.231 |
| SER-21 | -1.31433 | -1.727 |
| TYR-22 | -0.19233 | -0.81067 |
| ILE-23 | -1.03133 | -1.90833 |
| CYS-29 | -3.95333 | -5.22867 |
| GLY-30 | -3.37333 | -8.04067 |
| TYR-31 | -5.11633 | -11.0173 |
| GLY-32 | 0.518 | 0.25 |
| GLY-33 | 0.829 | -4.87833 |
| SER-34 | 0.077333 | -2.434 |
| GLY-35 | -1.04633 | -4.56067 |
| HIS-48 | -2.92967 | -5.12633 |
| ASP-49 | -42.9933 | -183.058 |
| TYR-52 | -3.01267 | 1.669 |
| LEU-64 | -7.43133 | -1.37667 |
| ILE-65 | -0.873 | 1.006667 |
| LYS-66 | 1.779667 | 61.35467 |
| THR-67 | 0.941333 | -0.24933 |
| TYR-68 | -0.70933 | 0.647333 |
| SER-69 | -0.11533 | -0.57433 |
| TYR-70 | -0.29667 | 0.053333 |

| | | |
|---|---|---|
| THR-71 | -0.00667 | -0.276 |
| CYS-72 | 0.039333 | 0.740667 |
| ALA-115 | 0.412 | 0.206333 |
| SER-116 | -0.32333 | -3.49867 |
| THR-117 | 0.025667 | -1.025 |

**Table S9.** Decomposition energy contributions of significant residues in svPLA₂ from 2WQ5 (Elapidae) during complex formation with LTNF10 and LTNF10-MT, calculated using MM/PBSA (values in kJ/mol).

| svPLA$_2$ residue | svPLA$_2$-LTNF10 | svPLA$_2$-LTNF10-MT |
|---|---|---|
| LEU-2 | 0.098 | 0.884667 |
| ILE-9 | 0.790333 | 0.917 |
| TRP-18 | -1.37333 | -1.07833 |
| TRP-19 | -2.22033 | -0.997 |
| ASP-20 | -31.0653 | -45.4867 |
| PHE-21 | -0.27933 | -1.377 |
| TYR-27 | -1.46467 | -7.291 |
| CYS-28 | -1.66167 | 0.234333 |
| GLY-29 | -2.77567 | 3.366333 |
| ARG-30 | 44.88067 | 60.52033 |
| GLY-31 | 0.61 | -5.30267 |
| GLY-32 | -0.97233 | -4.415 |
| SER-33 | -0.28933 | 0.450333 |
| HIS-47 | -0.964 | -4.035 |
| ASP-48 | -27.2687 | -149.111 |
| TYR-51 | 0.64 | 0.225667 |
| PHE-63 | -0.13633 | 3.684333 |
| PHE-64 | -0.03033 | 2.029667 |
| LYS-65 | 7.742333 | 71.15233 |
| THR-66 | 0.256667 | -1.686 |
| TYR-67 | -0.01033 | 1.604667 |
| SER-68 | -0.226 | -0.61333 |
| TYR-69 | -0.03533 | 0.552333 |
| LYS-70 | 7.135667 | 40.61367 |
| LEU-114 | -5.46567 | -0.59667 |
| LYS-115 | -73.647 | 47.97467 |

**Table S10.** Decomposition energy contributions of significant residues in svPLA$_2$ from 2QHD (Viperidae) during complex formation with LTNF10 and LTNF10-MT, calculated using MM/PBSA (values in kJ/mol).

| svPLA$_2$ residue | svPLA$_2$-LTNF10 | svPLA$_2$-LTNF10-MT |
|---|---|---|
| VAL-2 | -6.32533 | -4.34367 |
| ILE-9 | -2.64 | -0.375 |
| PRO-18 | 0.062333 | -0.14467 |

| | | |
|---|---|---|
| PHE-19 | -4.682 | -7.09733 |
| PRO-20 | 0.238 | -6.14333 |
| SER-21 | -0.60233 | -7.002 |
| TYR-22 | -11.6537 | -10.1873 |
| PHE-28 | -0.46333 | -3.77733 |
| CYS-29 | -7.80633 | -5.257 |
| GLY-30 | -3.84233 | -6.22533 |
| GLY-31 | 1.685 | -12.132 |
| GLY-32 | -2.16333 | -7.71467 |
| GLU-33 | -53.9997 | -195.309 |
| HIS-48 | 4.625 | -2.83267 |
| SER-49 | -8.03167 | 1.063 |
| TYR-52 | -7.21 | -10.5937 |
| SER-67 | 0.752333 | 5.064333 |
| PRO-68 | -1.26533 | -2.01533 |
| LYS-69 | -53.199 | 137.3057 |
| THR-70 | 1.758 | 3.27 |
| ASN-114 | 1.547667 | 0.994 |
| LYS-115 | -20.6183 | 28.02733 |
| LYS-116 | -11.1593 | 36.31933 |

**Table S11.** Decomposition energy contributions of significant residues in $svPLA_2$ from 4QEM (Viperidae) during complex formation with LTNF10 and LTNF10-MT, calculated using MM/PBSA (values in kJ/mol).

| **$svPLA_2$ residue** | **$svPLA_2$-LTNF10** | **$svPLA_2$-LTNF10-MT** |
|---|---|---|
| LEU-2 | -9.90267 | -3.129 |
| ILE-9 | -2.273 | -0.865 |
| ALA-18 | -1.67633 | -0.12233 |
| ILE-19 | -6.532 | -4.535 |
| PRO-20 | -0.11 | -1.099 |
| SER-21 | -0.03 | -0.97967 |
| TYR-22 | -0.55167 | -1.06333 |
| TYR-28 | -3.74533 | -9.252 |
| CYS-29 | -2.02933 | -6.82333 |
| GLY-30 | -4.406 | -7.505 |
| TRP-31 | -4.137 | -47.9093 |
| GLY-32 | -7.81867 | -8.14367 |
| GLY-33 | -4.76833 | -3.26967 |
| HIS-48 | -1.47833 | -6.51833 |
| ASP-49 | -68.551 | -220.105 |
| TYR-52 | -3.49167 | -4.89767 |
| ASN-67 | -2.33367 | 3.462 |
| PRO-68 | 1.617 | -11.505 |
| LYS-69 | -47.5243 | 111.0697 |
| SER-70 | 1.671333 | 1.293667 |

| | | |
|---|---|---|
| SER-114 | 0.760333 | -0.12433 |
| LYS-115 | -15.3153 | 32.68867 |
| LYS-116 | -5.33333 | 40.01333 |

**Table S12.** Decomposition energy contribution of residues from LTNF10 and LTNF10-MT peptides in binding with svPLA2 from 1U4J (Elapidae) calculated using MM/PBSA. Values are in kJ/mol.

| LTNF10 Residue | Decomposition energy | LTNF10-MT Residue | Decomposition energy |
|---|---|---|---|
| ALA-3 | 3.756 | ALA-3 | -4.00967 |
| ASP-5 | 62.07833 | LEU-1 | -57.5143 |
| LEU-1 | 20.09867 | LEU-10 | 12.98067 |
| LEU-10 | -30.7313 | LYS-2 | -74.685 |
| LYS-2 | -92.1597 | LYS-5 | -67.5137 |
| MET-4 | -23.709 | MET-4 | -11.69 |
| PRO-6 | -2.14033 | PRO-6 | 1.406333 |
| PRO-8 | -9.49167 | PRO-8 | 1.926667 |
| PRO-9 | -2.76667 | PRO-9 | -1.94567 |
| THR-7 | -5.51667 | THR-7 | 0.266333 |

**Table S13.** Decomposition energy contribution of residues from LTNF10 and LTNF10-MT peptides in binding with svPLA2 from 2WQ5 (Elapidae) calculated using MM/PBSA. Values are in kJ/mol.

| LTNF10 Residue | Decomposition energy | LTNF10-MT Residue | Decomposition energy |
|---|---|---|---|
| ALA-3 | 1.4 | ALA-3 | -1.75 |
| ASP-5 | 13.444 | LEU-1 | -118.18 |
| LEU-1 | -68.155 | LEU-10 | 33.37067 |
| LEU-10 | 8.823667 | LYS-2 | -72.4543 |
| LYS-2 | -86.3343 | LYS-5 | -30.7143 |
| MET-4 | -0.285 | MET-4 | 1.754333 |
| PRO-6 | 2.83 | PRO-6 | -8.46033 |
| PRO-8 | 2.545667 | PRO-8 | 0.388333 |
| PRO-9 | -1.277 | PRO-9 | -3.34433 |
| THR-7 | 4.160333 | THR-7 | 0.250333 |

**Table S14.** Decomposition energy contribution of residues from LTNF10 and LTNF10-MT peptides in binding with svPLA2 from 2QHD (Viperidae) calculated using MM/PBSA. Values are in kJ/mol.

| LTNF10 Residue | Decomposition energy | LTNF10-MT Residue | Decomposition energy |
|---|---|---|---|
| ALA-3 | -13.6837 | ALA-3 | -8.35067 |
| ASP-5 | -98.4817 | LYS-5 | -10.009 |
| LEU-1 | -49.2033 | LEU-1 | -160.706 |
| LEU-10 | -152.553 | LEU-10 | 46.49767 |
| LYS-2 | -40.5597 | LYS-2 | -20.075 |
| MET-4 | -9.58567 | MET-4 | -14.687 |
| PRO-6 | -23.8497 | PRO-6 | -16.713 |

| | | | |
|---|---|---|---|
| PRO-8 | -10.3863 | PRO-8 | -13.6217 |
| PRO-9 | -7.81267 | PRO-9 | -10.075 |
| THR-7 | -1.46267 | THR-7 | -3.154 |

**Table S15.** Decomposition energy contribution of residues from LTNF10 and LTNF10-MT peptides in binding with svPLA2 from 4QEM (Viperidae) calculated using MM/PBSA. Values are in kJ/mol.

| LTNF10 Residue | Decomposition energy | LTNF10-MT Residue | Decomposition energy |
|---|---|---|---|
| LEU-1 | -13.5933 | ALA-3 | -3.88633 |
| LYS-2 | 18.45267 | LEU-1 | -40.6363 |
| ALA-3 | -4.57967 | LEU-10 | -80.1103 |
| MET-4 | -11.265 | LYS-2 | -56.0437 |
| ASP-5 | -130.106 | LYS-5 | 82.88333 |
| PRO-6 | 6.545333 | MET-4 | -13.5413 |
| THR-7 | -6.66833 | PRO-6 | -9.23233 |
| PRO-8 | -10.892 | PRO-8 | -16.0503 |
| PRO-9 | -8.363 | PRO-9 | -16.8323 |
| LEU-10 | -109.355 | THR-7 | 1.574667 |

**Table S16.** Hydrogen Bond existence percentage (%) in LTNF10 and LTNF10-MT Complexes with svPLA$_2$ from 1U4J (Elapidae). (Rep- Replicate)

| | Rep1 | | | | | Rep2 | | | | | Rep3 | | | | |
|---|---|---|---|---|---|---|---|---|---|---|---|---|---|---|---|
| | Donor | | Acceptor | | % Existence | Donor | | Acceptor | | % Existence | Donor | | Acceptor | | % Existence |
| 1U4J-LTNF10 | TYR31 | HA | MET124 | HE3 | 10.511 | HIS50 | HA | PRO129 | HD2 | 13.979 | LYS54 | HD2 | PRO128 | HD2 | 18.358 |
| | TYR31 | HA | ASP125 | HN | 16.517 | HIS50 | HA | LEU130 | HD23 | 14.429 | LYS54 | HD2 | LEU130 | HD22 | 22.378 |
| | TYR31 | HA | ASP125 | HA | 10.511 | ASN53 | HA | ASP125 | HB2 | 45.355 | LYS54 | HD2 | LEU130 | HD23 | 20.978 |
| | HIS50 | HA | LEU130 | HD22 | 10.811 | LYS54 | HD2 | LEU130 | HD23 | 13.639 | ASN57 | HA | THR127 | HG23 | 15.958 |
| | HIS50 | HA | LEU130 | HD23 | 12.162 | LEU121 | N | TYR28 | HH | 54.955 | LYS122 | HD2 | ASP49 | HN | 54.305 |
| | ASN53 | HA | ASP125 | HB2 | 81.081 | LEU121 | N | CYS29 | HB2 | 10.029 | LYS122 | HD2 | ASP49 | HA | 40.646 |
| | LYS54 | HD2 | LEU130 | HD22 | 14.865 | LEU121 | N | ASP49 | HN | 35.546 | ASP125 | N | ASN53 | HN | 71.983 |
| | LYS54 | HD2 | LEU130 | HD23 | 12.312 | LEU121 | N | ASP49 | HA | 30.957 | THR127 | N | ASN53 | HD22 | 14.169 |
| | LYS122 | HD2 | HIS48 | ND1 | 74.625 | LYS122 | HD2 | ASP49 | HN | 15.208 | | | | | |
| | LYS122 | HD2 | ASP49 | HA | 12.312 | LYS122 | HD2 | ASP49 | HA | 14.309 | | | | | |
| | ASP125 | N | ASP49 | HN | 45.195 | MET124 | N | ASN53 | HN | 15.188 | | | | | |
| | ASP125 | N | ASP49 | HA | 25.976 | ASP125 | N | ASP49 | HA | 13.619 | | | | | |
| 1U4J-LTNF10-MT | LYS122 | N | TYR31 | HH | 10.999 | TYR31 | HA | LYS125 | HZ3 | 36.696 | SER34 | HA | ALA123 | HB3 | 31.407 |
| | LYS122 | N | GLY32 | HA2 | 16.798 | SER116 | N | THR127 | HG23 | 11.959 | SER34 | HA | LYS125 | HZ3 | 11.519 |
| | LYS122 | HD2 | TYR28 | HH | 10.109 | LEU121 | N | ASP49 | HN | 38.306 | HIS50 | HA | PRO128 | HD2 | 11.379 |

| | | | | | | | | | | | | | | | |
|---|---|---|---|---|---|---|---|---|---|---|---|---|---|---|---|
| | LYS122 | HD2 | ASP49 | HN | 31.377 | LEU121 | N | ASP49 | HA | 38.106 | HIS50 | HA | PRO129 | HD2 | 33.027 |
| | LYS122 | HD2 | ASP49 | HA | 40.116 | LEU121 | N | ASN53 | HN | 17.808 | LYS54 | HD2 | LEU130 | HD22 | 24.438 |
| | THR127 | N | ASN53 | HN | 64.214 | LYS125 | HD2 | SER114 | HA | 12.999 | LYS54 | HD2 | LEU130 | HD23 | 32.037 |
| | | | | | | LYS125 | HD2 | SER114 | HG1 | 49.055 | LYS122 | HD2 | GLY33 | HA2 | 10.679 |
| | | | | | | THR127 | OG1 | SER116 | HG1 | 10.209 | LYS122 | HD2 | SER34 | HG1 | 17.178 |
| | | | | | | THR127 | OG1 | THR117 | HG23 | 13.229 | LYS122 | HD2 | GLN120 | HE21 | 13.099 |
| | | | | | | | | | | | LYS122 | HD2 | GLN120 | HE22 | 14.419 |
| | | | | | | | | | | | THR127 | N | TYR46 | HA | 50.195 |

**Table S17.** Hydrogen Bond existence percentage (%) in LTNF10 and LTNF10-MT Complexes with svPLA$_2$ from 2WQ5 (Elapidae). (Rep- replicate)

| | **Rep1** | | | | | **Rep2** | | | | | **Rep3** | | | | |
|---|---|---|---|---|---|---|---|---|---|---|---|---|---|---|---|
| | **Donor** | | **Acceptor** | | **% Existence** | **Donor** | | **Acceptor** | | **% Existence** | **Donor** | | **Acceptor** | | **% Existence** |
| **2WQ5-LTNF10** | ARG42 | HG2 | LEU129 | HD22 | 33.947 | LYS65 | HD2 | LEU129 | HD22 | 25.757 | ARG30 | HB1 | LEU129 | HD22 | 16.848 |
| | ARG42 | HG2 | LEU129 | HD23 | 15.328 | LYS65 | HD2 | LEU129 | HD23 | 21.678 | ARG30 | HB1 | LEU129 | HD23 | 38.646 |
| | ARG42 | HE | LEU129 | HD22 | 11.359 | LEU120 | N | ARG30 | HH22 | 22.038 | ARG30 | HE | LEU129 | HD22 | 38.086 |
| | ARG42 | HE | LEU129 | HD23 | 30.147 | LEU120 | N | ASP48 | HN | 38.266 | ARG30 | HE | LEU129 | HD23 | 27.677 |
| | ASN49 | HA | LEU120 | H3 | 16.768 | LEU120 | N | ASP48 | HA | 25.957 | LEU114 | N | LYS121 | HZ3 | 15.088 |
| | ARG94 | HE | ASP124 | HN | 23.438 | LYS121 | HD2 | ASN52 | HN | 12.319 | LYS121 | HD2 | ASP23 | HN | 35.906 |
| | ARG94 | HE | ASP124 | HA | 11.209 | LYS121 | HD2 | GLU55 | HB1 | 25.337 | LYS121 | HD2 | ASP23 | HA | 35.646 |
| | LEU120 | N | GLY31 | HA2 | 16.198 | LYS121 | HD2 | GLU55 | HB2 | 24.418 | | | | | |
| | LEU120 | N | ASP48 | HN | 26.697 | | | | | | | | | | |
| | LYS121 | HD2 | GLU53 | HB1 | 15.518 | | | | | | | | | | |
| | LYS121 | HD2 | GLU53 | HB2 | 13.659 | | | | | | | | | | |
| | LYS121 | HD2 | GLU55 | HB1 | 10.339 | | | | | | | | | | |
| | LYS121 | HD2 | GLU55 | HB2 | 10.019 | | | | | | | | | | |
| **2WQ5-LTNF10-MT** | TRP18 | NE1 | MET123 | HE3 | 26.207 | TRP18 | NE1 | ALA122 | HB3 | 20.168 | ARG42 | HB1 | LEU129 | HD22 | 28.337 |
| | ARG30 | HG2 | LEU129 | HD23 | 10.199 | LEU120 | N | CYS28 | HB2 | 22.568 | ARG42 | HB1 | LEU129 | HD23 | 11.169 |
| | ARG30 | HE | LEU129 | HD22 | 10.389 | LEU120 | N | GLY29 | HA2 | 17.908 | ARG42 | HG2 | LEU129 | HD22 | 16.528 |
| | ARG30 | HE | LEU129 | HD23 | 10.649 | LEU120 | N | ASP48 | HN | 25.857 | ARG42 | HG2 | LEU129 | HD23 | 14.779 |
| | LYS115 | HD2 | LEU129 | HD22 | 12.529 | LEU120 | N | ASP48 | HA | 16.048 | ARG42 | HE | LEU129 | HD22 | 37.896 |
| | LYS115 | HD2 | LEU129 | HD23 | 11.999 | LYS121 | HD2 | ASP48 | HN | 14.339 | ARG42 | HE | LEU129 | HD23 | 47.535 |
| | LEU120 | N | HIS47 | ND1 | 34.687 | LYS121 | HD2 | ASP48 | HA | 14.509 | GLN45 | HB2 | THR126 | HG23 | 19.678 |

| | | | | | | | | | | | | | | | |
|---|---|---|---|---|---|---|---|---|---|---|---|---|---|---|---|
| | LEU120 | N | ASP48 | HN | 17.248 | LYS121 | HD2 | ASN52 | HN | 10.179 | ASN49 | HA | THR126 | HG23 | 26.307 |
| | LEU120 | N | ASP48 | HA | 22.188 | LYS121 | HD2 | GLU55 | HB1 | 18.428 | THR126 | OG1 | SER33 | HG1 | 10.559 |
| | LYS121 | HD2 | ASP48 | HN | 18.768 | LYS121 | HD2 | GLU55 | HB2 | 18.068 | | | | | |
| | LYS121 | HD2 | ASP48 | HA | 18.558 | | | | | | | | | | |
| | MET123 | N | CYS28 | HB2 | 22.608 | | | | | | | | | | |
| | MET123 | N | GLY29 | HA2 | 17.308 | | | | | | | | | | |
| | LYS124 | N | GLY29 | HA2 | 21.068 | | | | | | | | | | |
| | LYS124 | HD2 | ASP23 | HN | 36.706 | | | | | | | | | | |
| | LYS124 | HD2 | ASP23 | HA | 23.708 | | | | | | | | | | |

**Table S18.** Hydrogen Bond existence percentage (%) in LTNF10 and LTNF10-MT Complexes with $svPLA_2$ from 2QHD (Viperidae)

| | **Rep1** | | | | | **Rep2** | | | | | **Rep3** | | | | |
|---|---|---|---|---|---|---|---|---|---|---|---|---|---|---|---|
| | **Donor** | | **Acceptor** | | **% Existence** | **Donor** | | **Acceptor** | | **% Existence** | **Donor** | | **Acceptor** | | **% Existence** |
| **2QHD - LTNF 10** | LYS69 | HD2 | ASP138 | HN | 18.408 | THR23 | OG1 | LEU143 | HD22 | 11.609 | TYR25 | N | LEU143 | HD23 | 10.959 |
| | LYS69 | HD2 | ASP138 | HA | 16.078 | SER24 | N | LEU143 | HD22 | 51.005 | LYS69 | HD2 | ASP138 | HN | 11.729 |
| | LEU134 | N | GLU33 | HB1 | 14.849 | SER24 | N | LEU143 | HD23 | 46.285 | LYS69 | HD2 | ASP138 | HA | 13.229 |
| | LEU134 | N | GLU33 | HB2 | 15.328 | TYR25 | N | LEU143 | HD22 | 33.517 | LEU134 | N | GLU33 | HB2 | 34.577 |
| | LEU134 | N | ASP53 | HN | 14.289 | TYR25 | N | LEU143 | HD23 | 19.628 | LEU134 | N | ASP53 | HN | 23.028 |
| | LYS135 | HD2 | THR23 | HG23 | 20.148 | GLY32 | N | MET137 | HE3 | 28.347 | LEU134 | N | ASP53 | HA | 19.308 |
| | LYS135 | HD2 | TYR25 | HH | 14.499 | TYR113 | HA | LEU143 | HD23 | 31.977 | LYS135 | N | GLU33 | HB2 | 29.207 |
| | LYS135 | HD2 | GLY26 | HA2 | 30.567 | LEU134 | N | GLU131 | HG2 | 12.009 | LYS135 | HD2 | GLY31 | HA2 | 16.088 |
| | LYS135 | HD2 | CYS29 | HB2 | 38.476 | LYS135 | N | ASP53 | HN | 14.929 | LYS135 | HD2 | GLU33 | HB1 | 25.327 |
| | ALA136 | N | GLY30 | HA2 | 18.928 | LYS135 | N | ASP53 | HA | 17.058 | LYS135 | HD2 | GLU33 | HB2 | 20.598 |
| | ALA136 | N | GLY31 | HA2 | 10.779 | LYS135 | HD2 | ASP53 | HN | 35.306 | | | | | |
| | MET137 | N | GLY30 | HA2 | 14.079 | LYS135 | HD2 | ASP53 | HA | 27.807 | | | | | |
| **2QHD - LTNF 10- MT** | LYS132 | N | LEU134 | H3 | 10.529 | SER21 | N | LEU143 | HD23 | 88.471 | THR23 | OG1 | PRO142 | HD2 | 17.228 |
| | LYS132 | HD2 | LYS135 | HZ3 | 15.158 | TYR22 | N | LEU143 | HD23 | 96.84 | LYS69 | HD2 | ALA136 | HB3 | 15.348 |

| | | | | | | | | | | | | | | | |
|---|---|---|---|---|---|---|---|---|---|---|---|---|---|---|---|
| | LEU134 | N | GLU131 | HB1 | 12.149 | THR23 | N | LEU143 | HD23 | 97.97 | LEU134 | N | GLU33 | HB1 | 18.318 |
| | LEU134 | N | GLU131 | HB2 | 14.749 | SER24 | N | LEU143 | HD22 | 14.209 | LEU134 | N | GLU33 | HB2 | 19.088 |
| | LYS135 | HD2 | GLU33 | HB1 | 29.527 | TYR25 | N | LEU143 | HD22 | 92.771 | LEU134 | N | ASP53 | HN | 23.588 |
| | LYS135 | HD2 | GLU33 | HB2 | 31.127 | GLY30 | N | PRO141 | HD2 | 77.912 | LEU134 | N | ASP53 | HA | 21.388 |
| | ALA136 | N | ILE130 | HD3 | 13.629 | LEU134 | N | ASP53 | HN | 24.808 | LYS135 | N | GLU33 | HB1 | 17.328 |
| | MET137 | N | ASP53 | HA | 52.785 | LEU134 | N | ASP53 | HA | 21.628 | LYS135 | N | GLU33 | HB2 | 15.378 |
| | LYS138 | HD2 | GLY31 | HA2 | 43.046 | LYS135 | HD2 | GLU33 | HB1 | 19.408 | LYS135 | HD2 | GLY31 | HA2 | 11.879 |
| | LYS138 | HD2 | GLU33 | HB1 | 44.726 | LYS135 | HD2 | GLU33 | HB2 | 24.918 | LYS135 | HD2 | GLU33 | HB1 | 29.287 |
| | LYS138 | HD2 | GLU33 | HB2 | 42.616 | LYS138 | HD2 | GLY31 | HA2 | 29.377 | LYS135 | HD2 | GLU33 | HB2 | 27.007 |
| | | | | | | LYS138 | HD2 | GLU33 | HB1 | 37.796 | LYS138 | HD2 | HIS48 | ND1 | 13.249 |
| | | | | | | LYS138 | HD2 | GLU33 | HB2 | 29.107 | LYS138 | HD2 | SER49 | HA | 31.367 |
| | | | | | | THR140 | N | HIS48 | ND1 | 63.844 | | | | | |

**Table S19.** Hydrogen Bond existence percentage (%) in LTNF10 and LTNF10-MT Complexes with svPLA$_2$ from 4QEM (Viperidae)

| | **Rep1** | | | | | **Rep2** | | | | | **Rep3** | | | | |
|---|---|---|---|---|---|---|---|---|---|---|---|---|---|---|---|
| | **Donor** | | **Acceptor** | | **% Existence** | **Donor** | | **Acceptor** | | **% Existence** | **Donor** | | **Acceptor** | | **% Existence** |
| **4QEM-LTNF10** | LEU1 | N | ASN77 | HN | 13.079 | LEU1 | N | GLY42 | HA2 | 44.736 | LEU1 | N | ASP59 | HN | 35.096 |
| | LYS2 | HD2 | HIS58 | ND1 | 26.027 | LEU1 | N | ASP59 | HN | 66.943 | LEU1 | N | ASP59 | HA | 33.257 |
| | LYS2 | HD2 | ASP59 | HN | 22.498 | LEU1 | N | ASP59 | HA | 69.493 | SER33 | HA | LEU10 | HD22 | 27.377 |
| | LYS2 | HD2 | ASP59 | HA | 13.669 | LYS2 | N | ASP59 | HN | 82.622 | SER33 | HA | LEU10 | HD23 | 30.017 |
| | SER33 | HA | PRO9 | HD2 | 10.909 | LYS2 | HD2 | CYS143 | HB1 | 22.368 | LYS79 | HD2 | ASP5 | HN | 18.618 |
| | GLY40 | N | LEU10 | HD23 | 28.517 | LYS2 | HD2 | CYS143 | HB2 | 23.338 | LYS79 | HD2 | ASP5 | HA | 23.658 |
| | LYS79 | N | LEU1 | H3 | 10.269 | ALA3 | N | ASP59 | HA | 69.233 | | | | | |
| | LYS79 | HD2 | ASP5 | HN | 12.059 | THR7 | OG1 | CYS39 | HB2 | 10.249 | | | | | |
| | | | | | | SER33 | HA | LEU10 | HD22 | 32.577 | | | | | |
| | | | | | | SER33 | HA | LEU10 | HD23 | 28.217 | | | | | |
| | | | | | | LYS79 | HD2 | ASP5 | HN | 31.227 | | | | | |
| | | | | | | LYS79 | HD2 | ASP5 | HA | 27.857 | | | | | |

| | | | | | | | | | | | | | | |
|---|---|---|---|---|---|---|---|---|---|---|---|---|---|---|
| **4QEM-LTNF10-MT** | SER23 | HA | THR140 | HG23 | 43.036 | TRP31 | NE1 | ALA136 | HB3 | 11.449 | LEU134 | N | TRP31 | NE1 | 12.939 |
| | GLY30 | N | PRO142 | HD2 | 18.898 | LYS69 | HD2 | LEU143 | HD22 | 12.789 | LEU134 | N | TRP31 | HH2 | 74.783 |
| | TRP31 | N | PRO142 | HD2 | 13.439 | LYS69 | HD2 | LEU143 | HD23 | 12.129 | LEU134 | N | ASP49 | HN | 44.536 |
| | LEU134 | N | ASP49 | HN | 10.419 | | | | | | LEU134 | N | ASP49 | HA | 75.582 |
| | LEU134 | N | ASP49 | HA | 15.918 | | | | | | LYS135 | N | ASP49 | HN | 48.225 |
| | LEU134 | N | TYR120 | HH | 25.027 | | | | | | LYS135 | N | ASP49 | HA | 30.357 |
| | LEU134 | N | ASP122 | HN | 30.457 | | | | | | LYS135 | HD2 | ASP49 | HN | 18.878 |
| | LEU134 | N | ASP122 | HA | 15.168 | | | | | | LYS135 | HD2 | ASN54 | HN | 44.596 |
| | | N | SER24 | HA | 21.128 | | | | | | LYS135 | HD2 | CYS133 | HB1 | 26.997 |
| | | | | | | | | | | | LYS135 | HD2 | CYS133 | HB2 | 24.718 |
| | | | | | | | | | | | ALA136 | N | ASP49 | HA | 40.676 |
| | | | | | | | | | | | MET137 | N | PRO68 | HD2 | 23.198 |
| | | | | | | | | | | | THR140 | OG1 | TYR52 | HA | 10.039 |
| | | | | | | | | | | | LEU143 | N | LEU2 | HD23 | 41.246 |

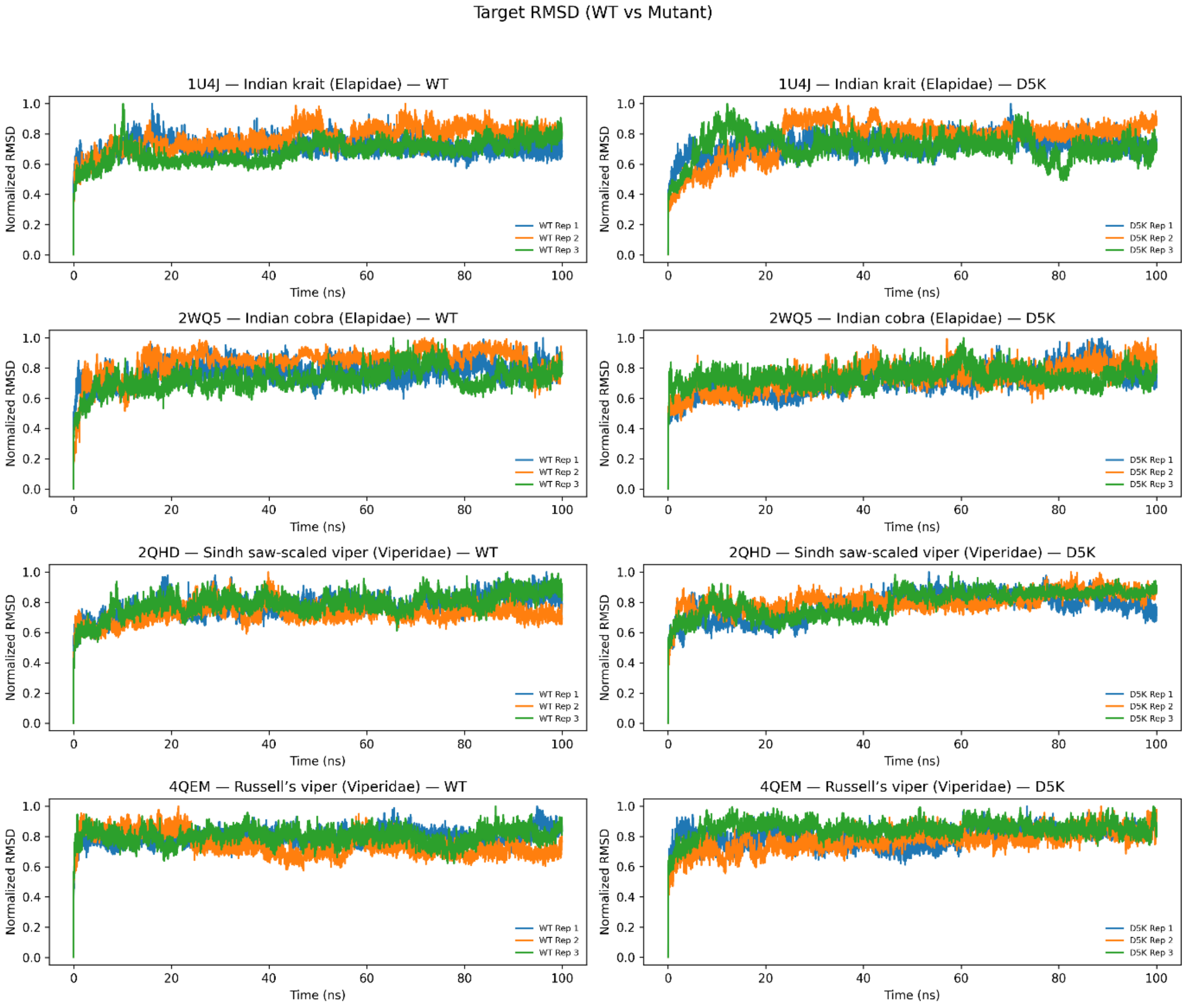


**Figure S24.** RMSD against backbone trajectories of svPLA$_2$ proteins with LTNF10 and LTNF10-MT across elapidae and Viperidae over 100ns.

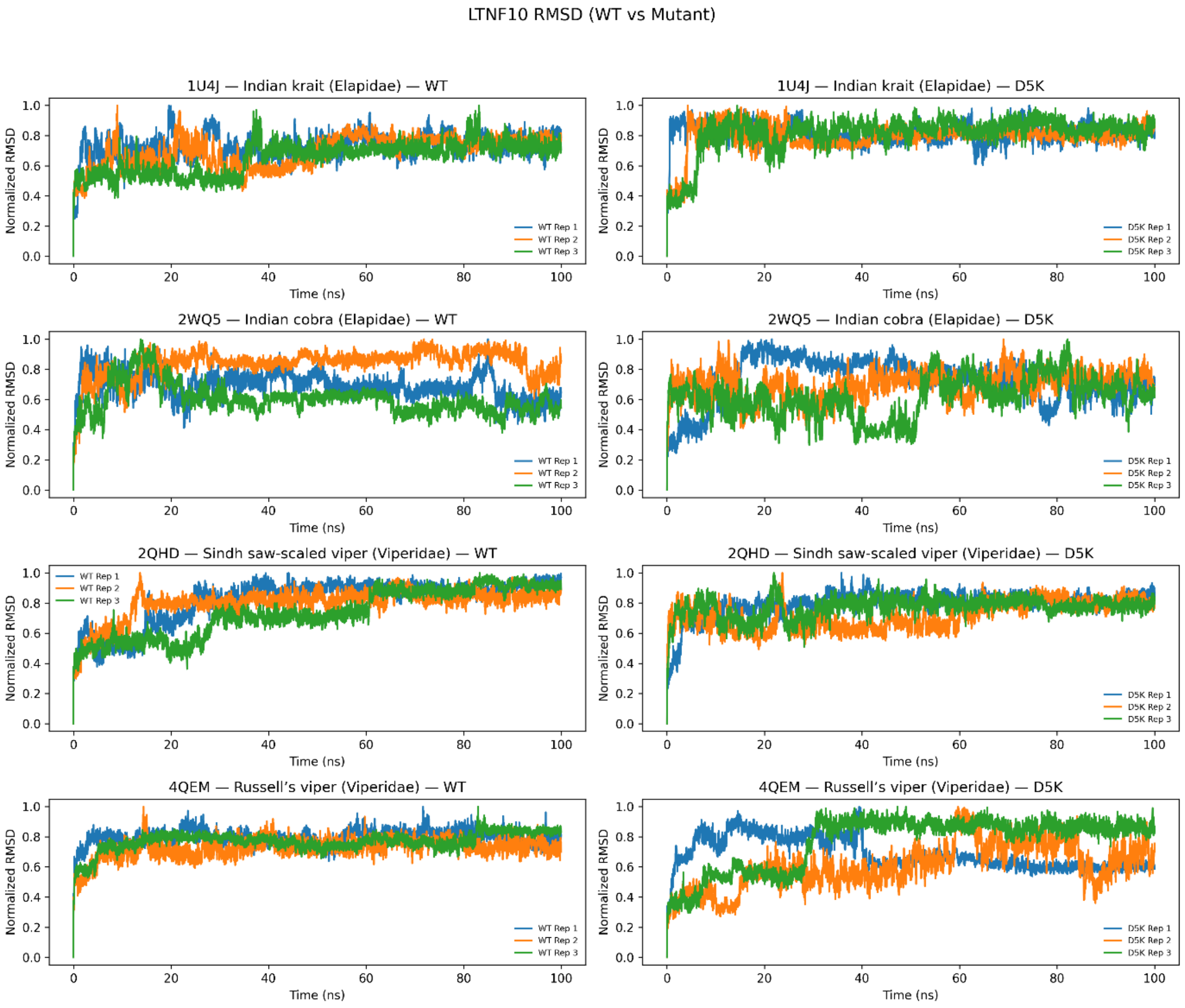


**Figure S25.** RMSD against backbone trajectories of LTNF10 and LTNF10-MT across elapidae and Viperidae over 100ns.

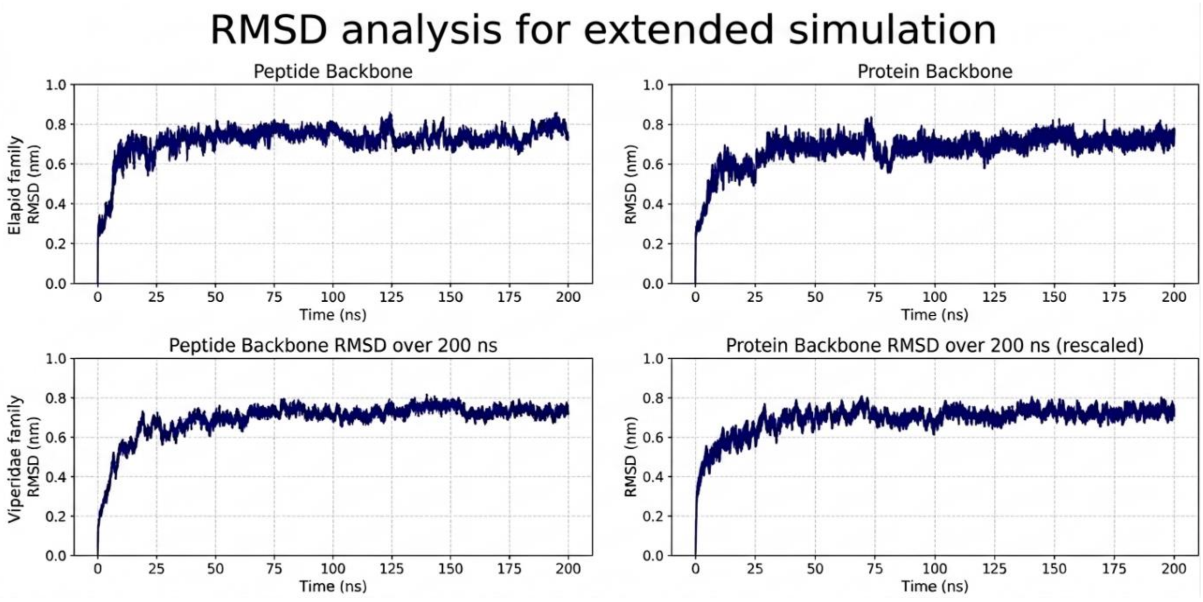


**Figure S26.** Backbone RMSD Trajectories of peptide and target backbone for Elapid (1U4J-LTNF10-MT) and Viperidae (4QEM-LTNF10-MT) complexes over extended simulation to 200ns.

**Table S20.** MM/PBSA binding free energy extended of simulation of 200ns compared with the original 100ns simulation. (1001 frames from the stable trajectory was used for both the analysis).

| System (Family) | ΔG_bind (kJ/mol) | |
|---|---|---|
| | 100ns | Extended to 200ns |
| 1U4J-LTNF10-MT (Elapidae) | -471.216 | -456.085 |
| 4QEM-LTNF10-MT (Viperidae) | -495.135 | -490.481 |

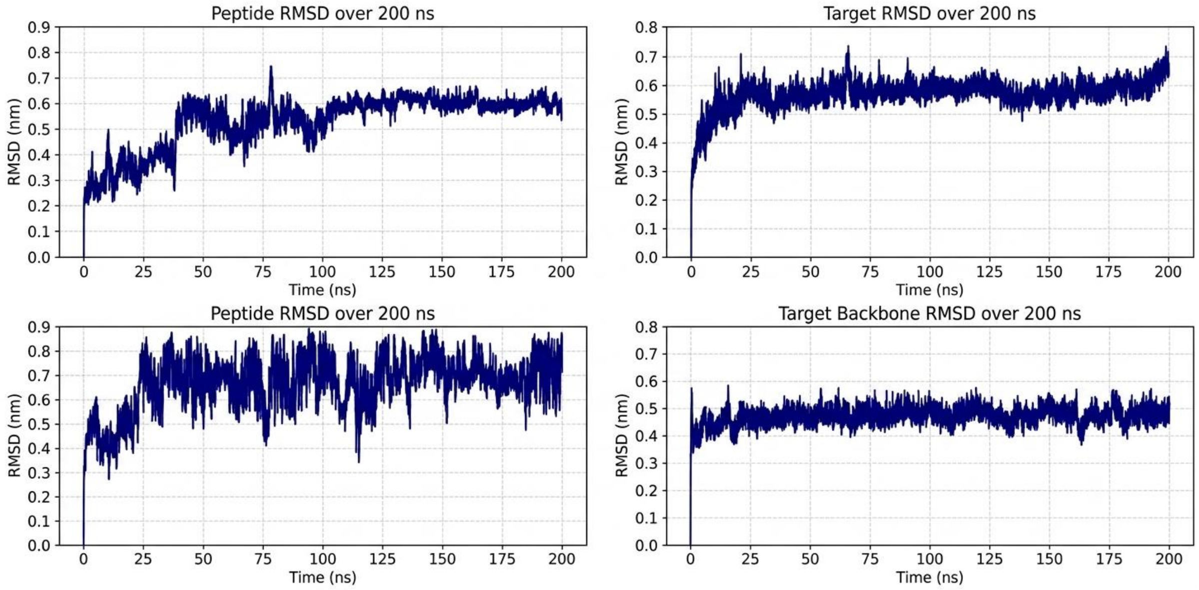


**Figure S27.** CHARMM36 validation runs (200 ns). Left column: peptide RMSD; right column: svPLA2 target RMSD. Top row: 1U4J (Elapidae); bottom row: 4QEM (Viperidae).

**Table S21.** MM/PBSA binding free energy for the CHARMM36 validation runs (1001 frames from the stable trajectory was used for both the analysis).

| System (Family) | ΔG_bind (kJ/mol) | |
|---|---|---|
| | CHARMM27 | CHARMM36 |
| 1U4J-LTNF10-MT (Elapidae) | -471.216 | -465.768 |
| 4QEM-LTNF10-MT (Viperidae) | -495.135 | -465.731 |

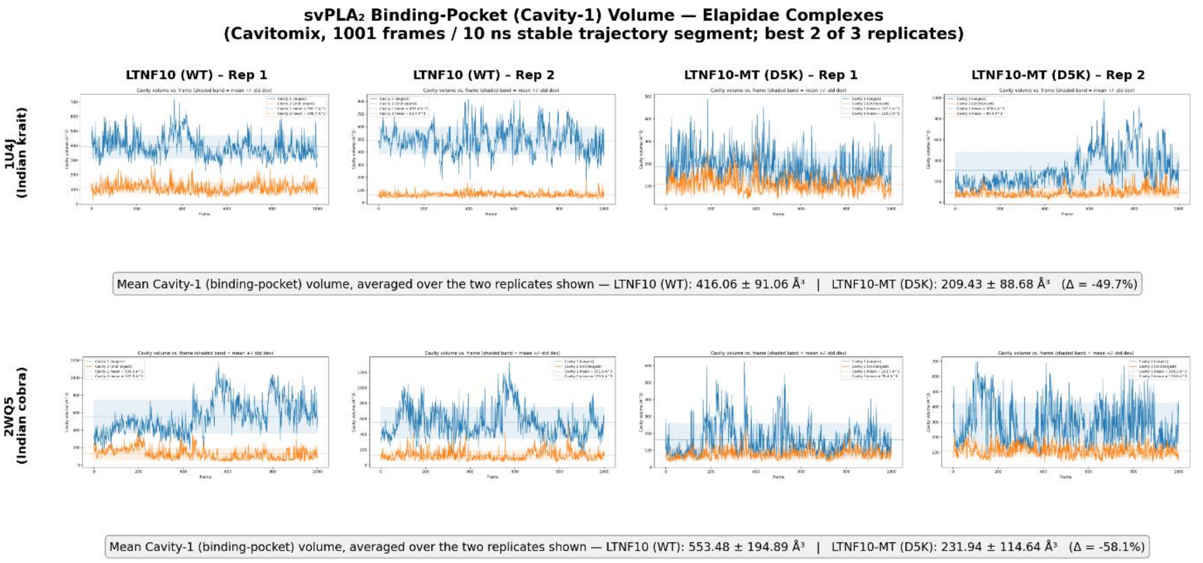


**Figure S28.** Cavitomix-derived binding-pocket (Cavity-1) volume dynamics of LTNF10/LTNF10-MT–svPLA$_2$ complexes in Elapidae.

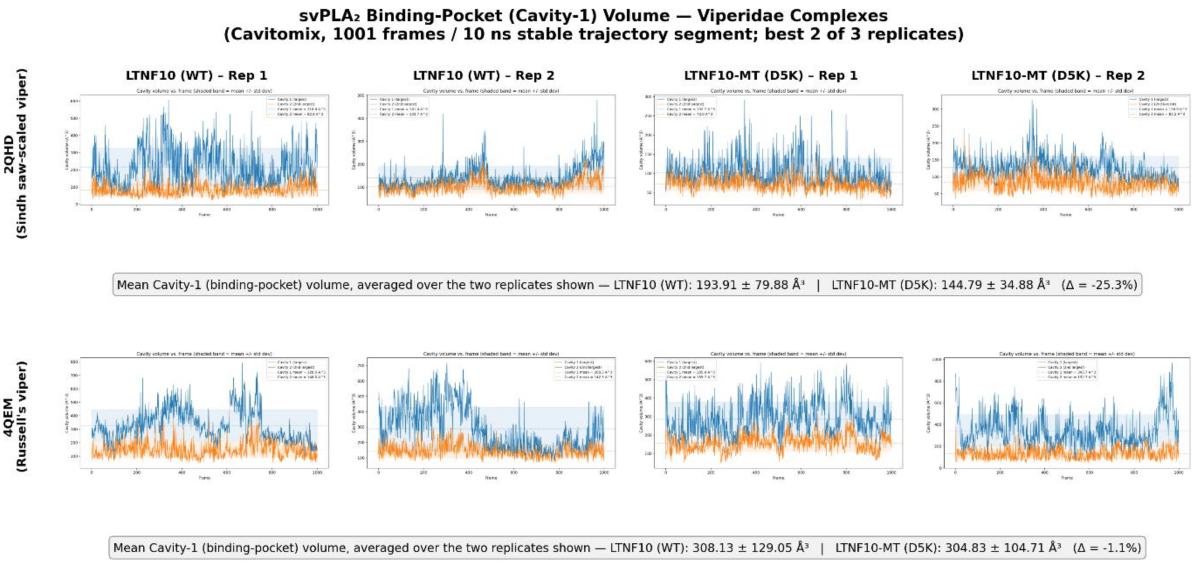


**Figure S29**. Cavitomix-derived binding-pocket (Cavity-1) volume dynamics of LTNF10/LTNF10-MT–svPLA$_2$ complexes in Viperidae

**Table S22.** Interaction similarity profile between LTNF10 and varespladib.

| svPLA$_2$ structural region | LTNF10 interactions | Varespladib interactions |
|---|---|---|
| N-terminal hydrophobic channel | Leu2, Ile9, Trp31, Tyr52, Asn52/53 | Leu2, Leu5, Ile9, Leu96 (6PWH, 7LYE) |
| Calcium-binding loop (~26–33) | Gly30, Gly32, Tyr25 | Asn27 backbone (6PWH, 7LYE); Gly31, Trp31 loop |

| | | |
|---|---|---|
| Catalytic site | His48, Asp49 | His47(ND1)/Lys48(NZ) (6PWH, 7LYE); His48/Lys49 (8DND) |

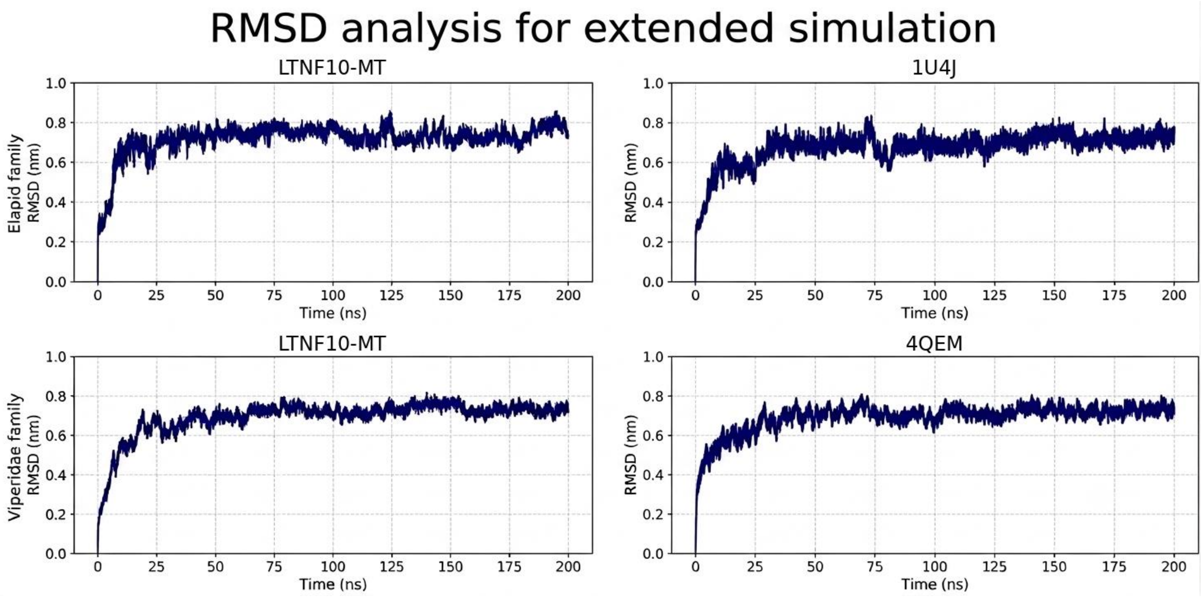


Figure S30. Backbone RMSD Trajectories of peptide and target backbone for Elapid (1U4J-LTNF10-MT) and Viperidae (4QEM-LTNF10-MT) complexes over extended simulation to 200ns.

**Table S23.** MM/PBSA binding free energy from an extended simulation of 200ns compared with the original 100ns simulation. (1001 frames from the stable trajectory was used for both the analysis).

| **System (Family)** | **ΔG_bind (kJ/mol)** | |
|---|---|---|
| | **100ns simuation** | **Extended simulation (200ns)** |
| 1U4J-LTNF10-MT (Elapidae) | -471.216 ± 61.227 | -456.085 ± 50.714 |
| 4QEM-LTNF10-MT (Viperidae) | -495.135 ± 59.169 | -490.481 ± 55.48 |
